# A pharmacological modality to sequester homomeric proteins

**DOI:** 10.1101/2025.03.05.641597

**Authors:** Ella Livnah, Ohad Suss, Adi Rogel, Atar Gilat, Yuval Abdan, Jose Villegas, Barr Tivon, Shira Albeck, Tamar Unger, Ofra Golani, Inna Goliand, David Margulies, Emmanuel D. Levy, Nir London

## Abstract

Molecules that can perturb protein-protein interactions have an immense impact on chemical biology and therapeutics. However, such compounds typically rely on accessory proteins to function, such as E3 ligases in the case of targeted degradation, which may restrict their target and tissue scope or lead to resistance. Here we alleviate the need for accessory proteins with a novel pharmacological modality to knock-down protein function. Our strategy exploits protein symmetry as a selective vulnerability, and is widely applicable owing to the ubiquitous nature of homomeric proteins in cellular systems. We target homomeric proteins with PINCHs (Polymerization Inducing Chimeras) - bifunctional molecules composed of two linked ligands that act as bridges between homomers and trigger their supramolecular assembly into insoluble polymers. We design PINCHs that achieve efficient polymerization of three homomeric targets. In cells, we observed that a PINCH targeting Keap1 exhibited a longer duration of action compared to its monomeric inhibitor, and a PINCH targeting BCL6 displayed selective and improved B cell toxicity compared to its monomeric parent. Our results highlight PINCHs as a novel and general strategy to modulate and knock out protein function.

## Introduction

In recent years, the induction of proximity between proteins^1–4^ has become a prevalent approach in chemical biology and drug discovery^5^. A number of modalities utilize bifunctional small molecules that bind a target protein on one end, and ‘hijack’ a cellular effector on the other end, to induce proximity between the two proteins and achieve a desired post-translational modification on the target. Prominent examples include PROTACs, to induce proteasomal degradation^6,7^, LYTACs to induce lysosomal degradation^8^, PHICs to induce phosphorylation^9^ and DUBTACs^10^ to induce stabilization via deubiqitination and RIPTACs^11^ to selectively target essential proteins. Monovalent proximity inducers, also known as molecular glues, are also gaining prominence^12,13^.

While targeted degradation by PROTACs offers several advantages over traditional inhibition^14^, these approaches also have a number of drawbacks. The bifunctional molecule has to bind two different molecular targets, which requires first and foremost, ligands for both proteins, but also for them to be co-expressed and co-localized in the relevant cellular compartment. In addition, some post-translational modifications (PTMs) have spatial restrictions, thus solely inducing proximity does not ensure successful modification or degradation of the target^15^. For example, PROTAC- or molecular glue-induced degradation of the target requires a correct positioning of the different components, in such a way that there is an available lysine on the surface of the protein for the E2 unit to ubiquitinate^7,16^. From a therapeutic perspective, resistance often stems from the recruited effector^17^. Despite the advances in proximity-based technologies to target proteins of interest^18^, a major challenge remains in expanding their use in a general way. There is a growing need for methods that can be rationally applied to a large range of proteins, minimizing the lengthy screening campaigns that are usually required for each new protein target.

Protein symmetry has long been leveraged to engineer molecular assemblies^19–22^. Recent work has suggested that small chemical changes at the surface of proteins can induce their polymerisation, provided that the protein target naturally forms a symmetric complex or homomer^23–26^. The robustness of this approach suggested that proteins are naturally ‘on the verge of’ oligomerization and formation of higher-order protein assemblies. Notably, monovalent small molecule modulators of a target’s oligomerization state have been mostly discovered serendipitously^27^; their mode of action typically discovered separately from their functional effect. One example is Trifluoroperazine, initially discovered as an inhibitor of S100A4, a member of the S100 protein family that has been associated with several human diseases^28^. This compound was eventually revealed to inhibit S100A4’s activity by stabilizing an inactive pentameric complex of it. Another example is a monomeric degrader of the transcription factor BCL6^29^, which was shown to polymerize the BTB domain of BCL6 and led to its proteasomal degradation. Recently, Diaz et al. reported a rationally designed trivalent inhibitor designed to capture three units of the Lipoprotein(a) K_IV_ domain in an inactive trimer^18^.

Here we rationally design small molecules that can take advantage of the properties of homomeric proteins to efficiently perturb their function. We aimed to achieve this by linking two identical target ligands with a flexible linker. A key design principle is that the bifunctional ligand should not bind two subunits of the same homomer simultaneously as this would form a closed, finite complex. Instead, if each ligand moiety binds a separate dimer, it would drive the assembly of an extended polymeric structure. This process effectively sequesters the protein in a polymeric state that can inhibit its normal function (Fig. 1a). Importantly, since protein homomers are common in nature, making up 25 to 50% of proteomes and exhibiting a variety of functionalities^30,31^, such an approach is expected to have a wide scope and be generally applicable.

**Figure 1.**
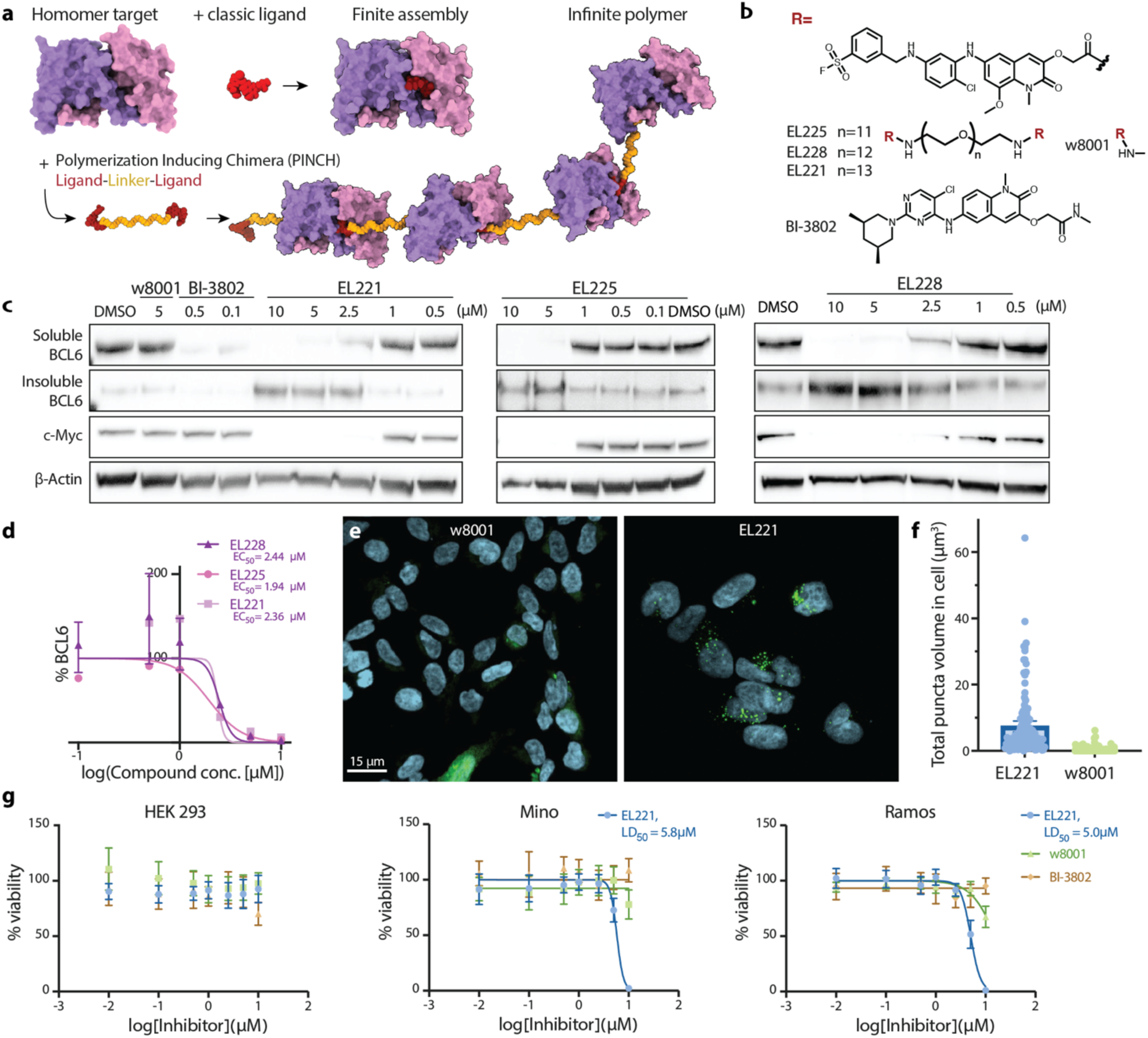
PINCHs precipitate BCL6 in cells and show cell type specific viability effects. **a.** Schematic representation of the PINCH mode of action. **b.** Structure of BCL6-targeting PINCHs. **c.** WB analyses of active BCL6-targeting PINCHs, 20 h treatment in Mino cells. **d.** WB band quantification and calculated EC_50_ values from n=3 independent experiments. Full gels are shown in Fig. S8 **e.** HEK-293 cells overexpressing BCL6(1-275)-GFP treated with parent ligand w8001 (left) and EL221 (right) at 5 μM, and **f.** the quantification of BCL-GFP ‘puncta’ in each treatment. Scale bars = 15 μm. (p<0.0001 in a two-tailed Mann-Whitney test from 223 cells analyzed in two independent experiments, additional images are shown in Fig. S6). One outlier was removed from the graph for clarity, but not from the statistical analysis. Error-bars show a 95% confidence interval. **g.** Viability of different cell lines after 48 h treatment with EL221, parent ligand w8001 and BCL6-polymerizer BI-3802 (representative results from two independent experimental replicates; n=6 technical replicates each).

We term these compounds PINCHs - Polymerization Inducing Chimeras, and present them in this work as a new modality for protein sequestration. We apply this approach to three target proteins: the homodimers BCL6 and Keap1, and the homotetramer LDHA. We show that PINCHs can induce polymerization *in vitro* and in cells, and precipitate the target proteins from the soluble phase to the insoluble phase. We show this polymerization is phenotypically different from inhibition or even degradation and can complement current approaches to perturb protein functions in cells.

## Results

### PINCH mediated accumulation of BCL6 in the cell’s insoluble phase

To evaluate the concept of PINCH as a novel modality, we initially focused on BCL6. The protein BCL6 is a homodimeric transcriptional repressor involved in B-cell regulation, and its overexpression is correlated with several B-cell non-Hodgkin lymphomas^32,33^. Thus, a pharmacological modality to knock down BCL6 is therapeutically relevant and makes it an excellent model system to test the PINCH approach. Several BCL6 inhibitors and degraders have been reported in recent years^34–36^. In addition, a BCL6 PROTAC that is currently under clinical trials demonstrated efficacy in multiple pre-clinical models of DLBCL^37^. Recently, BCL6 was also recruited by a new modality of bifunctional molecules to selectively repress alternative target genes^38,39^. Furthermore, it was discovered that BCL6 was able to polymerize via a small molecule glue^29^.

A covalent inhibitor of BCL6 that irreversibly binds a Tyrosine residue in the BTB domain dimeric interface was previously reported^40^. We used a close analog of this ligand (w8001; Fig. 1b) which showed rapid binding to recombinant BCL6 BTB domain by intact protein LC-MS (100% labeling at 2 μM protein/10 μM compound; 90 min; RT; Extended Data Fig. 1b). To create a PINCH targeting BCL6, we synthesized a compound (EL221; Fig. 1b) composed of two w8001 moieties linked by a PEG13 linker. Importantly, the linker had to be long enough to minimize potential steric clashes between bridged homodimers, yet short enough to forbid intra-molecular binding of subunits from the same homodimer.

We tested the effects of EL221 on BCL6 levels via Western Blot (WB) in Mino cells. At concentrations starting from 5 μM and after 20 h incubation we observed a reduction of BCL6 in the soluble fraction and an accumulation in the insoluble fraction (Fig. 1c). By contrast, the reported BCL6 polymerizing agent BI-3802^29^ showed no accumulation in the insoluble fraction, due to proteasomal degradation^7^.

### BCL6 PINCH induces different functional outcomes from its parent inhibitor and from a degrader

To assess the downstream effect of the PINCH-induced polymerization, we monitored the expression of c-Myc, a prominent target regulated by BCL6^41^. In the presence of EL221, BCL6 moved to the insoluble phase and c-Myc levels were markedly downregulated, implying that EL221 does hinder BCL6 cellular functions (Fig. 1c). Surprisingly, although BI-3802 downregulates BCL6, it showed no such effect on c-Myc, underscoring a potential difference in the mechanism of action of these modalities.

Proteomic analysis of EL221-treated cells revealed dozens of additional downregulated proteins (Fig. S1). We attribute this large and systemic effect to the centrality of BCL6 to B-cell biology as well as to the long incubation period (20 h).

Theoretically, using shorter linkers^42–44^, or a different linker chemistry in bifunctional molecules may affect the rigidity of the compound and its interactions with the protein, which could modulate the binding affinity. This motivated us to evaluate how changing the length of the PEG-based linker bridging the two binding moieties impacted the PINCH properties. We observed that the linker could be as short as 11 ethylene glycol units, but not less (Fig. 1b; Extended Data Fig. 1a). The similar potencies we observed for the longer linkers suggest that the PINCHs we examined were not stabilizing a neo BCL6-BCL6 interface but rather acted as simple bridges.

To further characterize whether EL221 induces the polymerization of BCL6, we visualized its cellular distribution following treatment in HEK293 cells expressing BCL6 (residues 1-275) fused to GFP. As cells were treated with EL221, the GFP signal accumulated into punctate structures, indicating that BCL6 formed higher-order assemblies. As a control, cells treated with the parent monomeric ligand (w8001) showed no such puncta. Instead, we observed a faint GFP signal spread throughout the cell (Fig. 1e). Indeed, the GFP puncta were significantly overrepresented in EL221 treated-cells compared to the monomeric ligand w8001 ( *p*< 0.001; Fig. 1f).

Finally, given that c-Myc is essential for B-cell viability, the marked downregulation in c-Myc expression following PINCH treatment (Fig. 1c) prompted us to evaluate its effect on cellular viability, and compare it to BI-3802 and w8001 (72 h). Neither compound affected the viability of HEK293 cells, which lack BCL6 expression. However, in Mino and Ramos B cells, both of which express BCL6, EL221 showed an LD_50_ of 5.8 μM and 5.0 μM respectively, while the degrader and inhibitor showed no visible effects (Fig. 1g). These results are in line with another report showing that BI-3802 exhibits viability effects on longer timescales^45^.

### Targeting Keap1 with the PINCH approach

To assess the broader applicability of the PINCH approach, we examined an additional system: the protein Keap1, a substrate adaptor for the E3 ubiquitin ligase complex Cul3-Rbx1, which plays a pivotal role in regulating Nrf2^46–48^. Nrf2 serves as a master regulator of the cellular anti-oxidative response, and proper functioning of the Keap1-Nrf2 pathway is essential in various diseases characterized by oxidative stress and inflammation^47,49,50^. Similarly to BCL6, Keap1 contains a BTB domain that mediates its homo-dimerization. Bardoxolone (BDX) is a reversible covalent binder of Cys151 in the BTB domain of Keap1 and is known to activate the Nrf2 pathway^51,52^. We therefore designed PINCHs targeting Keap1 based on Bardoxolone, two units of which were coupled via a PEG13 linker (EL229; Fig. 2a). Testing EL229 by WB showed it reduced Keap1 in the cell’s soluble fraction and induced its accumulation in the insoluble fraction (Fig. 2b).

**Figure 2.**
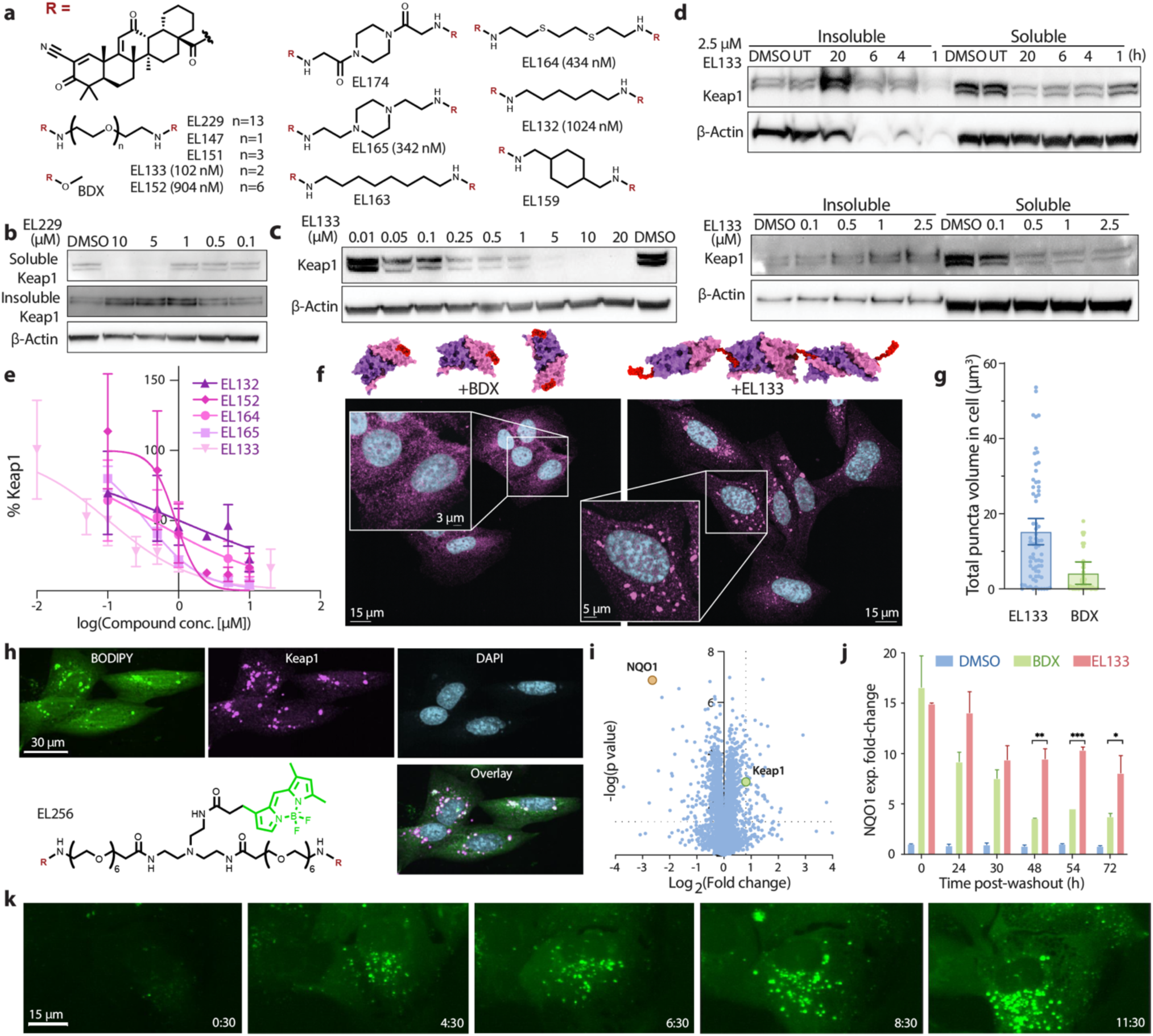
Keap1 PINCHs precipitate Keap1 in cells and have a long duration of action. **a.** Chemical structures of Keap1-targeting PINCHs. EC_50_ values for the active compounds are shown in parentheses. **b.** WB of EL229’s effect on Keap1 in OCI-AML2 cells’ soluble and insoluble phases; 20 h treatment. Full gels are shown in Fig. S10. **c.** WB analysis of additional active Keap1-targeting PINCHs in OCI-AML2 cells; 20 h. **d.** EL133 affects Keap1 in the soluble and insoluble OCI-AML2 cell phases, in a time- and dose-dependent manner. **e.** WB band quantification and calculated EC_50_ values from n=3 independent experiments. **f.** Immunostaining of U2OS cells following treatment with 250nM Bardoxolone (left) or 250nM EL133 (right). Keap1 is colored in Magenta; DAPI staining in Cyan. **g.** Quantification of Keap1 ‘puncta’ in the two treatments shows a significant difference in volume (*p*<0.0001, two-tailed Mann-Whitney test, n=129 cells analysed in 2 independent experiments, additional images are shown in Fig. S7). Error-bars show 95% confidence interval. **h.** Immunostaining of U2OS cells following a 20 hour treatment with 2.5 μM BODIPY-conjugated PINCH EL256, and its structure. Keap1 is colored in Magenta; DAPI staining in Cyan. **i.** Global proteomics analysis of OCI-AML2 cells soluble lysate, showing significant downregulation of Keap1 following treatment with EL133 (2.5 μM; 20 h), compared to DMSO. The dotted lines mark a 1.75-fold reduction in protein level (vertical) and *p*=0.05 significance (horizontal). **j.** rt-PCR analysis of NQO1 expression after a 20h treatment of OCI-AML2 cells with DMSO, EL133 or BDX at 250 nM, followed by washing of the cells (representative results following two independent experimental replicates with two technical replicates each). Error-bars show the standard deviation. **k.** Lattice light-sheet microscope live imaging of U2OS cells, immediately following treatment with 2.5 μM EL256. Time post-treatment is indicated in hr:min.

As for BCL6, we varied the linker length and composition (Fig. 2a; Extended Data Fig. 2a), which revealed a pronounced structure-activity relationship (SAR). EL152, based on an intermediary length PEG6, showed similar activity to EL229 (Extended Data Fig. 2a, EC_50_=904 nM). EL133, based on a PEG2, was the most potent compound with EC_50_=102 nM (Fig. 2c) and led to the accumulation of Keap1 in the insoluble phase at a concentration as low as 500 nM (Fig. 2d). Moreover, this accumulation occurred in a dose- and time-dependent manner (Fig. 2d). However, similar PEG-based linkers EL147 and EL151 based on PEG1 and PEG3 respectively, showed no activity, highlighting that the PINCH modality can be highly sensitive to linker length (Extended Data fig 2a). Given this apparent sensitivity, we designed several compounds based on the PEG2 design. First, EL164 differed with a sulfur atom replacing the PEG-oxygen, and showed good activity in cells, downregulating Keap1 with an EC_50_ value of 434 nM (Extended Data Fig. 2a). Replacing the middle ethylene glycol unit with a piperazine (EL165) also retained activity in cells (EC_50_ of 342 nM). In another design, EL132, we used a 6-carbon alkyl linker and downregulated Keap1 with an EC_50_ of 1024 nM. By contrast, three close analogs of these active compounds: EL163 based on an 8-carbon alkyl linker (the same length as EL133), EL159 which is the same length as EL132 but is conformationally restricted, and EL174 which differs from EL165 by two carbonyl groups, showed no activity (Extended Data Fig. 2a). This subtle SAR mirrors similar trends in bifunctionals such as PROTACs in which small changes in the linker can dramatically impact cellular potency, in part due to cooperative neo-PPIs induced by the bifunctional compound^43,53,54^.

Analyzing the crystallographic symmetry of the Keap1-Bardoxolone complex (PDB: 4CXT; Fig. S2A, B), we identified a neighboring Keap1 monomer from an adjacent unit cell positioned such that the Bardoxolone binding sites are close. We could model EL133 acting as a bridge across this crystallographic interface of two Keap1 dimers (which is not the biological dimer interface; Fig. S2C, D). This suggests a binding mode through which EL133 could stabilize a low-energy, non-native interface for Keap1, and therefore bind cooperatively. This may also explain the observed particularity in the linker length and chemical characteristics that allow efficient polymerization of Keap1.

### PINCH-mediated downregulation is selective and proteasome independent

To evaluate the selectivity of EL133, we performed quantitative proteomics while focusing either on the entire cellular protein content or using only the soluble fraction after cell lysis. Subjecting the entire lysate to proteomics showed no significant reduction in Keap1 levels compared to DMSO-treated controls (Extended Data Fig. 2b), likely due to the disassembly of the Keap1 polymer in the insoluble fraction, enabling its detection. By contrast, when we lysed cells using a RIPA buffer and focused solely on the soluble fraction, Keap1 was significantly downregulated in the treated sample relative to the DMSO-treated control (Fig. 2i). Moreover, only a few other proteins were down-regulated (Dataset S1), demonstrating selectivity of EL133 for Keap1.

Keap1 is also a substrate adapter protein to a Culling-ring E3 ligase complex, and was shown to induce degradation of targets to which it is recruited^55–58^. Thus, the Keap1 downregulation we observed may result from its self-degradation in a homo-PROTAC manner. Such self-targeting was previously observed for other E3 ligases^59–61^. To test this possibility, we initially treated cells either with MLN-4924 (a NEDDylation inhibitor), or with Bortezomib (a proteasome inhibitor), followed by EL133. Inhibiting either of these targets did not rescue Keap1 downregulation by EL133 (Extended Data Fig. 2c), implying that EL133 downregulates Keap1 in a proteasome-independent manner.

We visualized the sub-cellular localization of endogenous Keap1 following PINCH treatment by immunofluorescence in U2OS cells. Similarly to BCL6, we observed bright punctate structures in the EL133-treated cells and not in those treated with the parent ligand Bardoxolone (*p*<0.001; Fig. 2f, g).

To follow the cellular localization of the PINCH molecules themselves, we produced a BODIPY-conjugated PINCH, (EL256; Fig. 2h). Its design was based on the long linker of EL229 to minimize interference of the fluorophore with polymerization. A western-blot analysis of EL256-treated cells confirmed it had retained such ability (Extended Data Fig. 2a). Immunostaining EL256-treated U2OS cells revealed that endogenous Keap1 puncta were co-localized with the BODIPY signal (Fig. 2h), implying incorporation of the PINCH into the Keap1 assemblies, and further supporting our proposed mechanism of action for this modality. Taken together, these data indicate that the PINCH downregulates Keap1 by triggering its polymerization into non-soluble, supramolecular assemblies.

The co-localization also suggests that we may follow the compound’s fluorescent signal as a proxy for the formation of Keap1 polymers in cells. We utilized fluorescent live-U2OS imaging and EL256 to follow the protein’s higher-order assembly formation in real-time. This revealed an initial slow uptake of the compound by the cells, followed by puncta formation at 4.5 hours post-treatment (Fig. 2k). Over the course of several additional hours we observed the formation and expansion of additional puncta (See Movies S1, S2, S3).

### Keap1 PINCHs are longer acting than classic monomer inhibitors

We observed that PINCHs could drive a target protein into forming large macromolecular assemblies, reducing its levels in the soluble fraction and potentially inhibiting its function. This unique mechanism of action prompted us to explore whether it is linked to distinct pharmacodynamic properties, particularly in relation to the duration of action. NQO1 is a well-established downstream target of Nrf2^62,63^ and is therefore expected to be upregulated when Keap1 activity decreases. NQO1 was the most significantly upregulated protein in our soluble proteomics experiment (Fig. 2i). We measured NQO1 expression levels over a 72-hours time course by quantitative PCR after subjecting cells to either DMSO, Bardoxolone, or EL133 for 20 hours, followed by washout. At the time of washout both the Bardoxolone-treated and the EL133-treated cells showed a similar ∼15-fold increase in the expression level of NQO1. However, 48 h post-treatment, NQO1’s expression in the Bardoxolone-treated cells returned closer to its original level whereas it remained higher in the EL133-treated cells (48h *p*=0.0069; 54h *p*=0.0007; 72h *p*=0.0370; Fig. 2j). This result suggests a long-lasting effect of the PINCH and differentiates this modality from traditional inhibition of the target. We performed the same comparison with Omaveloxolone (OMVX), a derivative of Bardoxolone which has recently received FDA approval for the treatment of Friedrich’s Ataxia^64^, in which it had similar effects on NQO1 expression as Bardoxolone (Fig. S3). This observation highlights a potential advantage of the PINCH modality relative to other inhibition modes, whereby the duration of action is typically limited by clearance of the compound or half-life of the protein.

### In vitro characterization of PINCH-induced polymerization

Experiments in cells involve a complex milieu that may affect the PINCH activity. In order to characterize the mechanism of action of PINCHs independently of potential confounding variables, we evaluated the Keap1-PINCH *in vitro* with a purified recombinant BTB domain. Additionally, since Keap1 and BCL6 both involve a BTB domain, we also synthesized and similarly evaluated a PINCH targeting LDHA, a structurally unrelated homotetramer.

We used Dynamic Light Scattering (DLS) to follow the polymerization of Keap1 BTB domain in the presence of EL133. After about four hours, we observed large molecular species forming that did not appear in samples containing only Keap1 or EL133 (Fig 3a). Since Bardoxolone is a *reversible* covalent binder of Keap1, we hypothesized that EL133-induced polymer formation should be reversible as well, such that addition of monomeric Bardoxolone could dissolve the large assemblies. Indeed, adding two equivalents of Bardoxolone led to the disassembly of Keap1 polymers (Fig 3a). Surprisingly, this phenotype did not reproduce in the cellular environment: When treating cells with EL133 for 20 h and then adding a large excess of Bardoxolone no change in Keap1’s levels in the soluble phase occurred, as seen by WB (Fig 3b).

**Figure 3.**
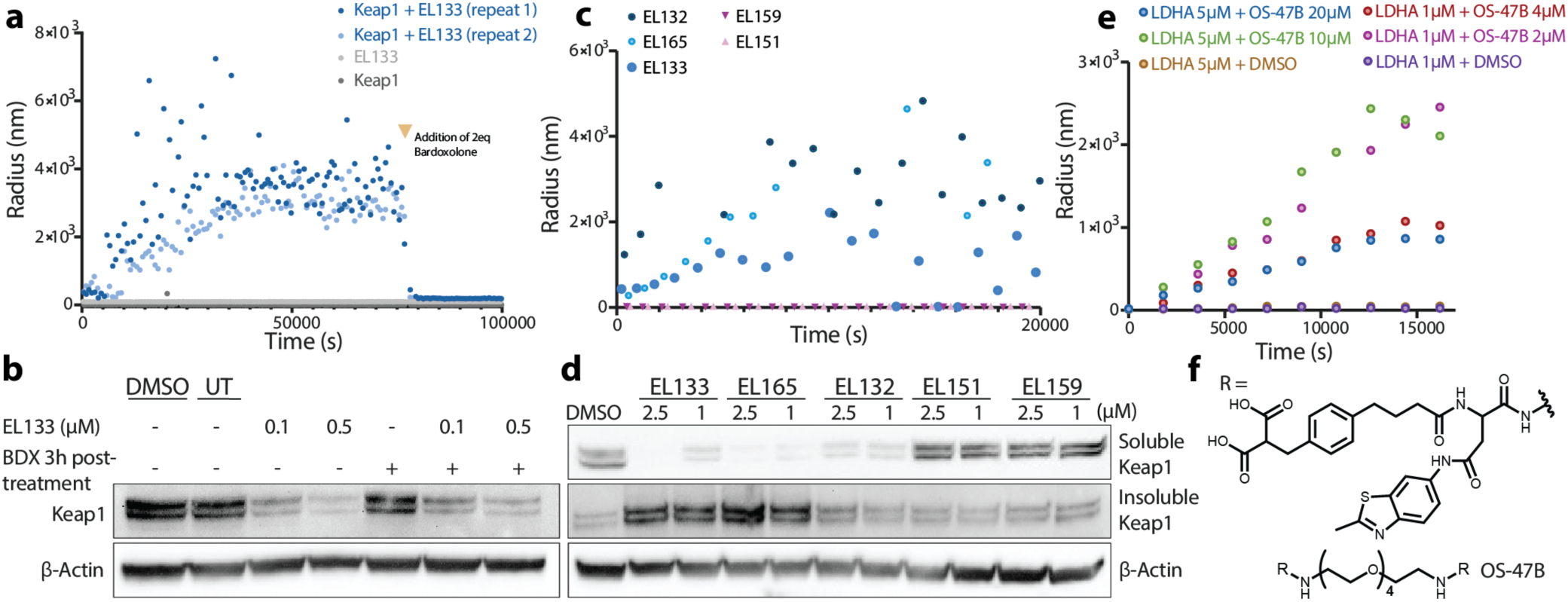
PINCHs induce target polymerization *in vitro*. **a.** Dynamic light scattering experiments following recombinant Keap1-BTB (50 μM) and EL133 (50 μM) over time at room temperature. **b.** WB of 20 h EL133-treated OCI-AML2 cells with and without post-treatment with excess BDX (5 μM). Full gels are available in Fig. S12. **c.** Dynamic light scattering experiments following recombinant Keap1-BTB (50 μM) and additional Keap1-targeting PINCHs (50 μM) over time at room temperature, and **d.** These PINCHs’ respective WB analysis in OCI-AML2 cells (20 h). **e.** Dynamic light scattering experiments following recombinant LDHA incubated with OS-47B over time at room temperature. Each LDHA bears four binding sites, and each PINCH bears two binding moieties (e.g., 5 μM LDHA= 20 μM binding sites). **f.** Chemical structure of LDHA-targeting PINCH OS-47B.

We tested additional Bardoxolone-based PINCHs with recombinant Keap1 in the same manner and observed the formation of large species over time with EL165 and EL132, but not with EL151 and EL159 (Fig 3c), mirroring these compounds’ activities in cells (Fig 3d). In addition, pre-incubating Keap1 with Bardoxolone prior to adding EL133 prevented the formation of large Keap1 structures, showing that these are Bardoxolone-binding-site dependent (Fig. S4A).

Finally, we applied the PINCH strategy to Lactate dehydrogenase A (LDHA) which forms a homotetrameric structure. We functionalized a parent inhibitor developed by Ward et al^65^ to synthesize an LDHA-targeting PINCH (OS-47B; Fig. 3f). DLS measurements of recombinant LDHA incubated with the PINCH revealed the formation of larger species over time (Fig 3e). A molar ratio of 1:1 in binding moiety to binding site resulted in the largest *in vitro* species, nonetheless, a polymerization effect was also observed in 2-fold excess in favour of either binding pocket or binding ligand. However, an excess of binding moieties (up to 8-fold) led to progressively smaller assemblies, presumably due to saturation of the LDHA binding sites (Fig. S4B). Also, sub-concentrations of PINCH (down to 1:8 binding moieties to binding pockets) led to smaller assemblies, as expected from a lower concentration of “crosslinks”.

### PINCH induced assemblies are long lived with minimal toxicity

A question that remains open is what happens to the PINCH-induced polymers that form and precipitate in the cells and how does the cell clear these formations, if at all. To explore this, we treated OCI-AML2 cells with EL133 for 20 hours to induce Keap1’s polymerization, followed by washing the cells and probing them over several time points (Extended Data Fig. 3a). After treatment with 500 nM EL133, Keap1 is absent from the cell’s soluble phase and remains accumulated in the insoluble phase for up to the 96 hours that we tested. It is interesting to note that BCL6 polymers do show gradual clearance in the same time frame, in a similar washout experiment, despite the fact that soluble BCL6 remains undetected (Extended Data Fig. 3b).

To assess whether the presence of the aggregate was toxic, we performed viability experiments in BDX or EL133 treated cells, following washout. We measured LD_50_ values three days and six days post wash (Extended Data Fig. 3c). The LD_50_ of EL133 comprising two units of BDX (254 nM after 3 days) was lower than that of BDX (956 nM after 3 days). However, the LD_50_ values after 3 or 6 days of EL133 treatment remained similar (294 nM after 6 days of EL133). These results suggest the presence of the aggregates themselves do not incur additional toxicity over Bardoxolone’s inherent toxicity^66^.

### A Hetero-PINCH simultaneously downregulates Keap1 and BCL6

In certain scenarios, the simultaneous elimination of more than one target is of therapeutic or biological interest. Accordingly, we propose that two homomeric targets could be induced to co-precipitate by a single PINCH. To illustrate this concept, we synthesized a PINCH bearing Bardoxolone and w8001 on either end (Fig. 4a), to respectively bind Keap1 and BCL6 dimers. Theoretically, this molecule should induce the formation of a polymer consisting of Keap1 and BCL6 dimers in an alternative fashion.

**Figure 4.**
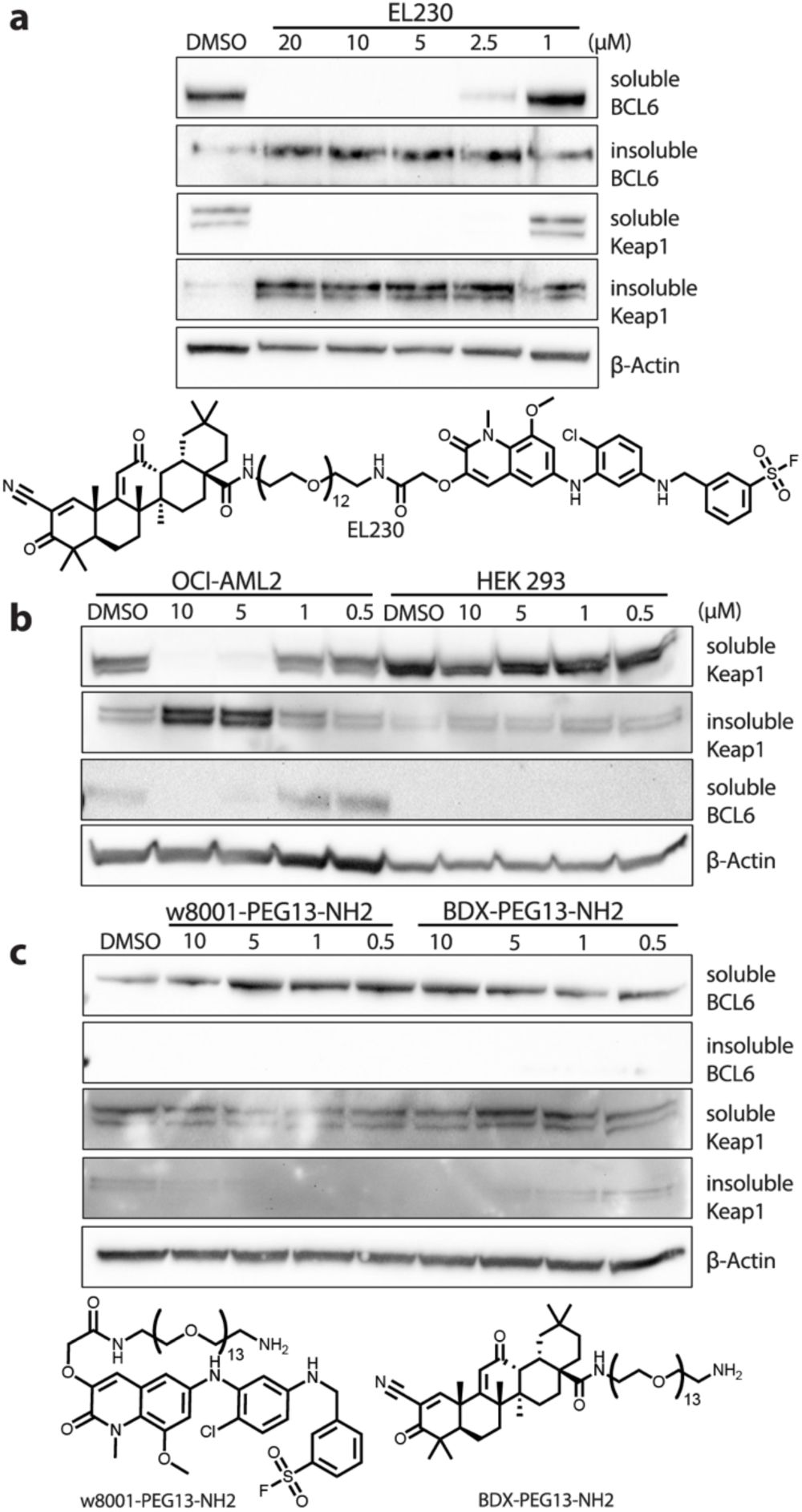
Hetero-PINCHs can simultaneously co-precipitate two targets. **a.** WB and structure of EL230, 20h treatment in Mino cells. Full gels are available in Fig. S13. **b.** EL230 tested in a BCL6-positive cell line (OCI-AML2) and in a BCL6-absent cell line (HEK 293), to show inactivity in the latter. **c.** Monomeric versions of EL230, based on BCL6- and Keap1-ligands with linkers, in Mino cells.

We tested the effect of the hetero-PINCH, EL230, in Mino cells that express high levels of BCL6, and we observed a reduction of both Keap1 and BCL6 levels in the soluble phase, and their corresponding accumulation in the insoluble phase, in a dose-dependent manner (Fig. 4a). To test the dependence of the hetero-PINCHs activity on the presence of both target proteins, we tested EL230 in two additional cell lines, OCI-AML2 and HEK293 cells, as the latter does not express BCL6. In OCI-AML2 cells Keap1 and BCL6 were downregulated simultaneously, while Keap1 remained unchanged in HEK293 cells (Fig. 4b). In addition, monomeric versions of the compounds, consisting of one ligand attached to a linker did not induce such effects on either target in Mino cells (Fig. 4c).

## Discussion

We introduce a novel pharmacological modality for inactivating protein targets through a mechanism that relies on their induced sequestration by polymerization. We applied this approach against two homodimeric proteins in cells and against a homotetramer protein *in vitro*.

This modality is reminiscent of PROTACs, which also involve bifunctional ligands and induce the degradation of a protein target. However, PROTACs require binding to a specific E3 ligase, and thereby necessitate its co-expression and co-localization with the target protein of interest. By contrast, PINCHs do not require such additional machinery besides the target itself. Additionally, while PROTACs require a precise interaction geometry to promote ubiquitination, the PINCH modality, in principle, does not require a highly specific interaction geometry to promote high-order assembly and sequestration. This relaxed requirement can be exploited to scout initial activity using long linkers (Fig. 1b, Fig. 2b), while optimization of the linker length and chemical nature for maximal potency could be performed later (Fig. 2c and Extended Data Fig. 2a).

While PINCHs are associated with relaxed constraints in terms of the subunit recognition geometry, they do require the bifunctional ligand to meet two conditions: (i) act as a bridge between two oligomers, and (ii) minimize the formation of finite oligomers such as dimer of dimers (Fig. S5) or more generally closed rings. Taking into account the homo-oligomer structure can help in target selection and ligand design for both considerations. Additionally, we anticipate homo-oligomers with dihedral symmetry will be less susceptible to form closed geometries^19,25,67,68^. As an example, a recent dimeric binder of Keap1 was reported to inhibit it in cells, without sequestration from the soluble fraction^69^. On top of these geometric constraints, concentration is an important property to consider, whereby a ligand concentration that is too low or too large would inhibit high-order assembly through the ‘hook’ effect.^70,71^ Additionally, although we show examples for non-covalent (LDHA), reversible-covalent (Keap1) and irreversible covalent (BCL6) PINCHs, the role of covalency in promoting efficient polymerization, and its interplay with ligand-target stoichiometry and kinetics remain to be fully characterized, as was done for other covalent proximity inducing modalities^72–74^.

Phenotypically, the PINCHs tested in cells exhibited different functional outcomes compared to their monomeric parent inhibitors. BCL6 PINCHs showed complete elimination of c-MYC, which was not seen with either the parent monomeric inhibitor, or degrader (Fig. 1c). Similarly, at 72h only the PINCH induced a viability effect (Fig. 1g) without apparent toxicity in cells not expressing the target. Additionally, Keap1 PINCHs showed significantly longer duration of action when compared to the monomeric Bardoxolone (Fig. 2j), perhaps due to the persistence of the aggregate following compound washout. While the precise mechanisms underlying these differences warrant further investigation, these results clearly demonstrate that PINCHs can complement other pharmacological modalities.

The scope of this technology might seem limited by the requirement for symmetrical oligomeric targets. However, our recent analysis^75^ suggests 20% of the human proteome fulfills this requirement as homo-oligomers. Moreover, an additional 30% of the proteome participates in symmetric hetero-oligomers that can in principle be targeted by PINCHs. The requirement for symmetry also adds a dimension of selectivity for PINCHs. While the monomeric ligand recruiter might not be very selective (like Bardoxolone for example), only a fraction of its off-targets would also fulfill the symmetry and geometry requirements for polymerization. This may be reflected in the relatively few targets downregulated by EL133 (Fig. 2i).

To conclude, we present here a novel modality for eliminating a target protein from the cellular soluble phase, by inducing its polymerization. This approach can target a large fraction of the human proteome, and complement existing modalities. Its simple design-principles should facilitate its adoption and application against a variety of new targets, expanding our pharmacological toolbox along with the space of targetable proteins.

## Supporting information

Supplemental Figures and Chemistry

Supplemental proteomics dataset

## Extended Data Figures

**Extended Data Figure 1.**
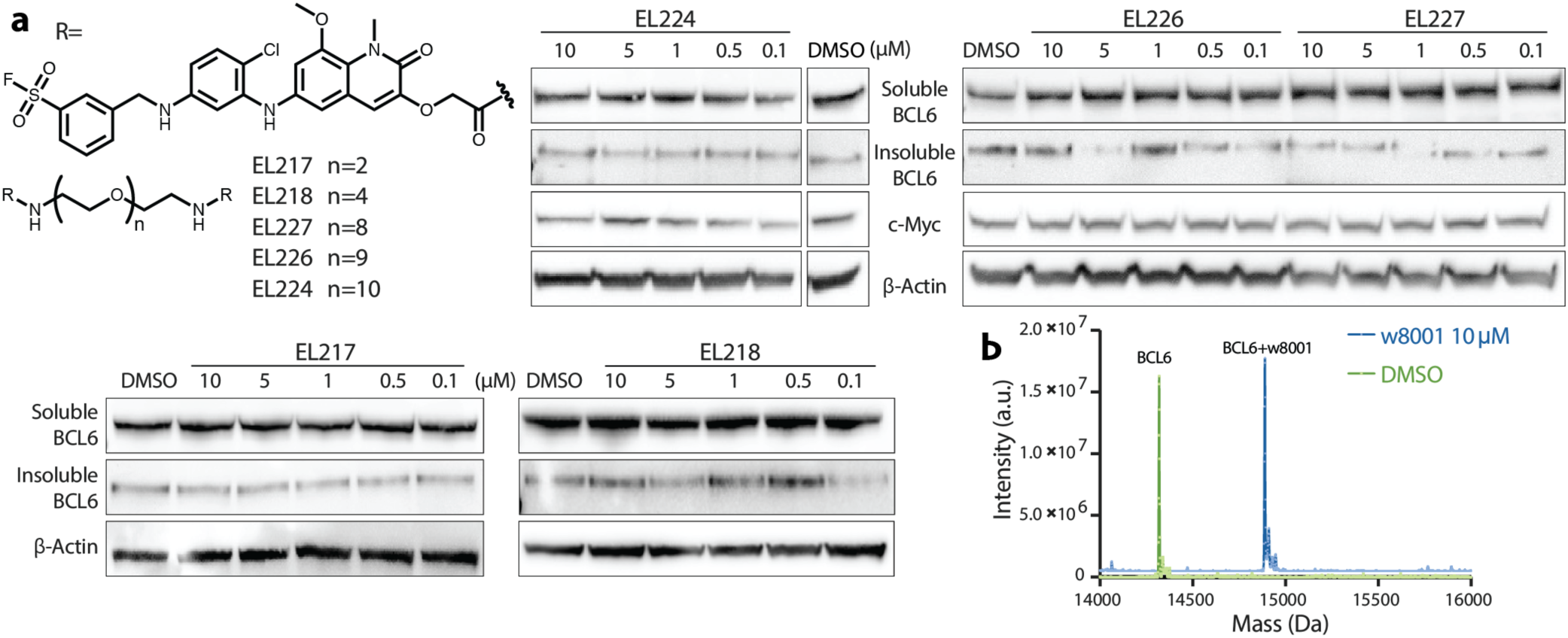
SAR of BCL6 targeting PINCHs. **a.** Structures of inactive BCL6-targeting PINCHs and their respective WB analyses in Mino cells at 20 h treatments. Full gels are available in Fig. S9. **b.** Intact LC-MS analysis of 2μM BTB-BCL6 after incubation with DMSO/w8001 for 1 hour at room temperature.

**Extended Data Figure 2.**
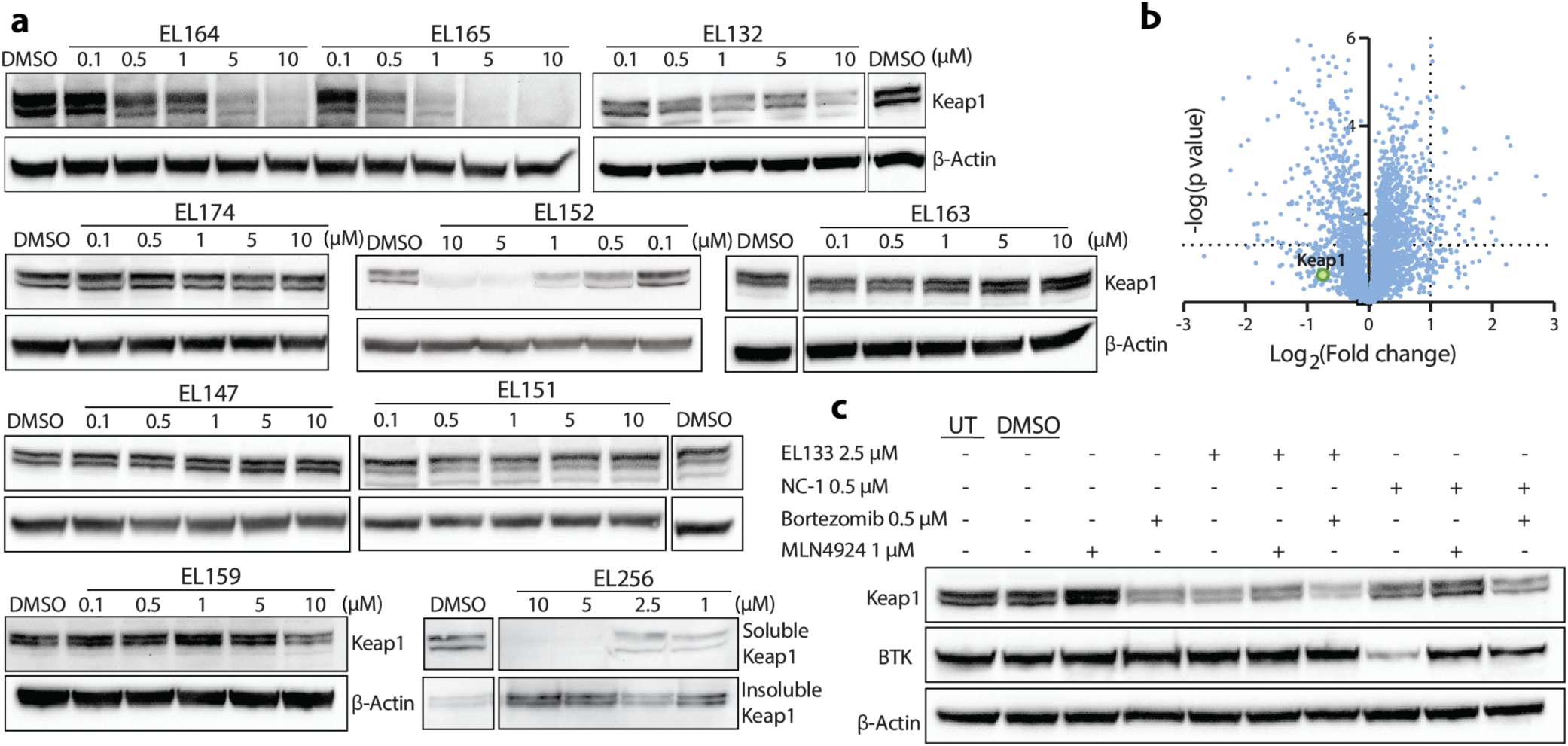
SAR and mechanistic characterization of Keap1 PINCHs. **a.** WB analyses of Keap1-targeting PINCHs in OCI-AML2 cells at 20 h treatments. All full gels are available in Fig. S10,11. **b.** Global proteomics analysis of OCI-AML2 cells treated with 2.5μM EL133/DMSO, lysed by sonication prior to sample analysis. **c.** OCI-AML2 cells pre-treated with Bortezomib or MLN4924 for 1 hour, then with EL133 or NC-1, a BTK-targeting PROTAC^76^, for an additional 7 hours.

**Extended Data Figure 3.**
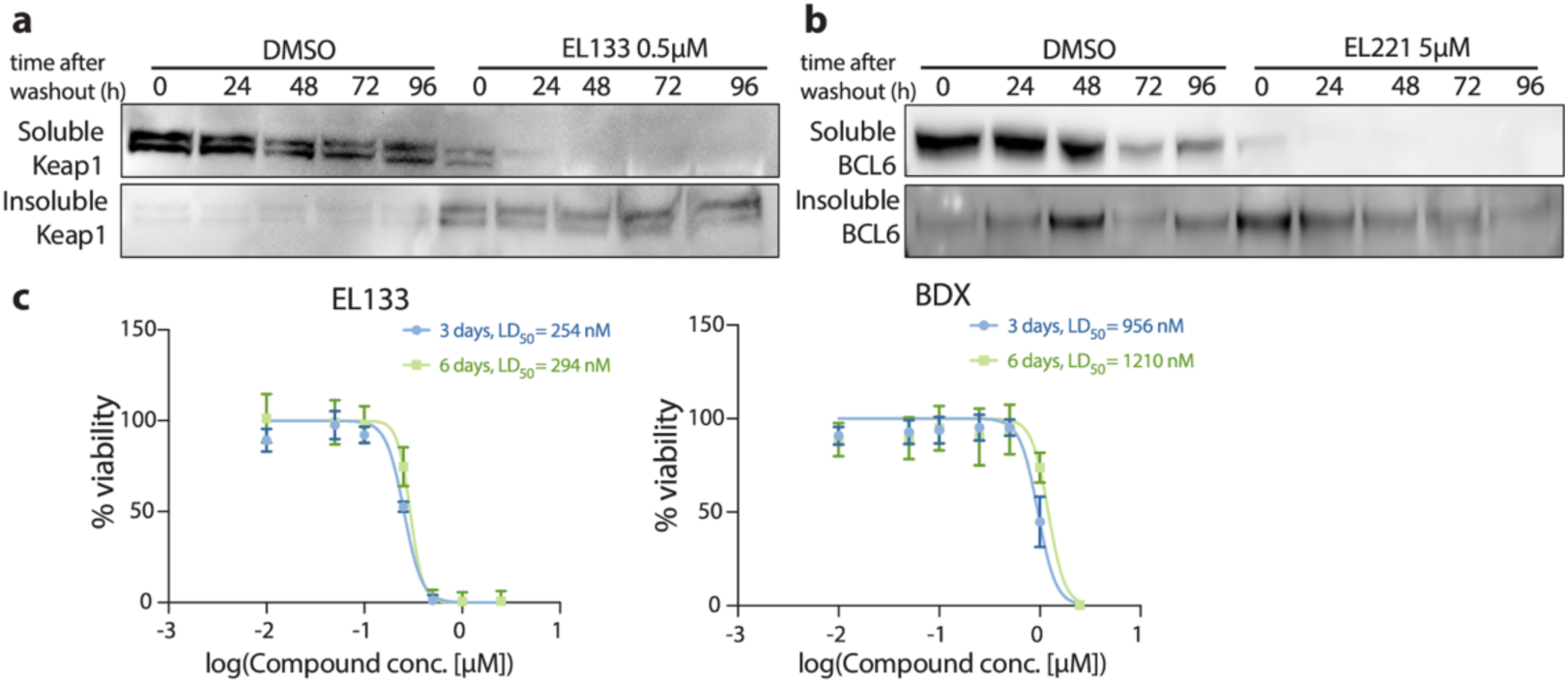
Duration of PINCH induced precipitation and effects on toxicity. **a.** WB of OCI-AML2 cells treated with 500nM EL133 for 20 h followed by washing of the cells, re-seeding and collected at different times after washout. Full gels are available in Fig. S12. **b.** WB of Mino cells treated with 5μM EL221 for 20 h followed by washing of the cells. **c.** Viability of U2OS cells at 3 and 6 days after treatment with BDX or EL133, (n=6 technical replicates).

## Methods

### Western Blotting

Cells were incubated with DMSO at 0.1% or with compound in concentrations ranging 10-20,000 nM for 20 hours unless indicated differently. Following incubation, lysis was performed in RIPA buffer and the samples were measured for total protein quantification by BCA assay (Thermo Fisher Scientific, 23225). 40 μg of each sample was loaded and run on an SDS-PAGE gel, then transferred onto a nitrocellulose membrane using a Trans-Blot Turbo system (Bio-Rad) and blocked using a 5% BSA in TBS-T (w/v) solution for 1 h at room temperature. Incubation with primary antibodies (mouse anti-BCL-6 (sc-7388, Santa Cruz, 1:500, overnight at 4 °C), mouse anti-Keap1 (sc-365626, Santa Cruz, 1:500, overnight at 4 °C), rabbit anti-c-Myc (#5605, Cell Signaling, 1:1000, overnight at 4°C), rabbit anti-p62 (ab155686, Abcam, 1:1000, overnight at 4 °C) and mouse anti-beta-actin (#3700, Cell Signaling, 1:1000, 1 h at room temperature) was performed, then the membranes were washed and incubated with an HRP-conjugated secondary antibody (mouse #7076/rabbit #7074, Cell Signaling) at room temperature for 1 hour. Imaging of Chemiluminescence signal was performed in a ChemiDoc XRS+ instrument (Biorad).

### Label-Free Quantitative Proteomics

Cells were incubated with DMSO at 0.1% or with compound for 20 hours unless indicated differently, in quadruplicates. Following incubation, the cells were lysed in either a 5% SDS and 50mM Ammonium Bicarbonate solution or in a mixed detergent buffer (Cells were lysed using detergent based buffer containing the following: (50mM HEPES pH 8.0, 1% SDS, 1% Triton X-100, 1% NP-40, 1% Tween 20, 1% sodium deoxycholate, 50mM NaCl, 5mM EDTA, 1% (w/v) glycerol), and probe-sonicated for the shearing of DNA. In samples where only soluble-phase proteomics was done, gentler RIPA-buffer lysis was performed, followed by discarding of the insoluble protein fraction. Samples were then reduced using DTT, followed by reaction with iodoacetamide. The labeled proteins were then precipitated on a dispersion of glass beads, followed by overnight trypsin digestion and purification of peptides. Label-free quantitative proteomics was applied for comparison of protein abundance between DMSO and compound-treated samples. The instrumentation includes nano flow liquid chromatography (nanoAcquity) coupled to tandem, high resolution mass spectrometry (Q Exactive HF). The data will be processed using MaxQuant^77^ or FragPipe^78,79^. The mass spectrometry proteomics data have been deposited to the ProteomeXchange Consortium via the PRIDE^80^ partner repository with the dataset identifiers PXD054711 and PXD054669, for Dataset S1 first and second sheets, respectively.

### Purification of KEAP1_S172A (48-180)

KEAP1 BTB domain (48-180) encoding a single mutation S172A was cloned into pET28-bdSumo vector generating His-bdSumo-KEAP1_ S172A. A 7.5 L culture of BL21(DE3) was induced with 200 µM IPTG and grown at 18°C overnight. The cells were harvested and lysed by a cooled cell disrupter (Constant Systems) in lysis buffer (50 mM Tris pH 8, 150 mM NaCl, 1 mM DTT, 10 mM imidazole and 10% glycerol) containing 0.2 mg/ml lysozyme, 20 µg/ml DNase, 1 mM MgCl_2_, 1 mM phenylmethylsulfonyl fluoride (PMSF) and protease inhibitor cocktail. After clarification by centrifugation (38,000 g for 30 min), the lysate was incubated with 5 ml Ni beads (Adar Biotech, prewashed with lysis buffer) for 1 h at 4°C. After removing the supernatant by centrifugation, the beads were washed 3 times with 50 ml lysis buffer and once with lysis buffer containing 50 mM imidazole (1 mM TCEP replacing the DTT). KEAP1 eluted from the beads by incubation of the beads with 20 ml cleavage buffer (50 mM Tris pH 8, 0.15 M NaCl, 1 mM DTT, 2 mM MgCl_2_, 10% glycerol and bdSumo protease) for 2 h at room temperature. The supernatant containing the cleaved KEAP1 was concentrated and applied to a size exclusion column (HiLoad_16/60_Superdex200, GE Healthcare) equilibrated with 150 mM NaCl, 25 mM Tris-HCl pH 8.0, and 1 mM TCEP. Fractions containing pure KEAP1 were pooled, concentrated and frozen (−80°C) in aliquots

### Purification of BCl6 BTB domain (5-129)

The BCL6 BTB domain (5-129) was cloned into the pET28-bdSumo vector generating His-bdSumo-BCL6_BTB. The cell culture, lysis and capture on Ni beads were all performed as described for KEAP1 BTB with the exception that PBS was used as the buffer for lysis and Ni capture. Cleaved BCL6 eluted from the beads by incubation of the beads with 20 ml cleavage buffer (PBS supplemented with bdSumo protease) for 2 h at room temperature. The supernatant containing the cleaved BCL6 was pooled and diluted x3 with 50 mM Tris pH 8 and loaded onto a Tricorn Q 10/100 GL anion exchange column. Pure BCL6 eluted at 250 mM NaCl when a linear gradient to 1 M NaCl was applied. The pooled pure protein was supplemented with 5 mM DTT and flash frozen (−80°C) in aliquots.

### Dynamic Light Scattering

DLS samples were prepared in the appropriate Keap1 buffer, to which 0.01% Tween was added, along with the respective compound from a DMSO stock to obtain a final DMSO concentration of 2%. Samples were prepared at room temperature and measured at a constant temperature of 25^0^C. All samples were passed through 0.1 μm filters, then were placed in a clear-bottom black 384-well plate. DLS data were recorded using a DynaproPlate Reader III (Wyatt Technology). Data was processed with the supplied DYNAMICS software.

### Quantitative Real-Time PCR (qPCR) for NQO1 expression

Cells were treated with either DMSO or compound at 250 nM, followed by washing out the treated media after 20 h and reseeding in fresh media. Cells were then collected at several time points post washout. Total RNA was extracted using RNeasy mini kit (QIAGEN, 74104). RNA concentration and purity was determined by Nanodrop, and 500 – 1000 ng RNA per sample were used for cDNA synthesis using High-Capacity cDNA Reverse Transcription Kit (Applied Biosystems, 4368814). 73 qPCR (10 µl reaction in a 96-well plate format) was performed with Fast-SYBR Green Master Mix (Applied Biosystems, 4385612), using 1-5 ng cDNA and the following primers (ordered from IDT): NQO1 Fw: CCGTGGATCCCTTGCAGAGA, NQO1 Rev: AGGACCCTTCCGGAGTAAGA, GAPDH Fw: ACCCACTCCTCCACCTTTGA, GAPDH Rev: CTGTTGCTGTAGCCAAATTCGT. PCR amplification was carried out in Step-One Plus thermocycler (Applied Biosystems). Relative quantification was performed using standard curves, GAPDH was used for normalization.

### Viability measurements

Cells were seeded in a transparent-bottom, black 96-well plate, at 15,000 cells per well in sixfold replications, in 90 µL media and let adhere for at least 4 hours. The cells were then treated with the respective compound by adding 10 µL of media-compound mixture for a final volume of 100 µL in each well. Cell-titer Blue reagent was used to measure viability according to the manufacturer’s instructions. Plate reader measurements were performed on a Tecan Spark Control 10 M fluorescent system.

### Immunostaining

For spinning disk confocal, cells were allowed to adhere to a 14mm glass-bottom plate (MatTek P35G-1.5-14-C), followed by fixation with 3.7% Formaldehyde for 5 minutes and permeabilized with 0.1% Triton X-100 for 10 minutes. The cells were then blocked with 5% BSA for 60 min at room temperature. Cells were stained with primary antibody, at a concentration of 1:100, for 3 hours at room temperature, and were then subjected to a secondary antibody conjugated to Alexa Fluor, at a concentration of 1:100-500 for 1 hour at room temperature. Lastly, cells were also stained with DAPI for 10 minutes at room temperature.

### Immunostaining Imaging

Imaging of the cells was performed using a Dragonfly spinning disk confocal system (Andor Technology PLC) connected to an inverted Leica Dmi8 microscope (Leica Microsystems CMS GmbH). The signals were detected by an sCMOS Zyla (Andor) 2048X2048 camera, and the bit depth was 16. At the specified intervals, Z-stacks (0.17 µm) of selected cells were acquired with an HC PL APO 63×/1.3 GLYC CORR CS2 objective (506353, Leica Microsystems CMS GmbH).

### Image Analysis

Total aggregate volume was calculated per cell using Imaris software (version 9.9.1). Automatic aggregates segmentation was done using the surface module with manual threshold fixed for all images of the same experiment (default 2 pixels smoothing, background subtraction with 0.9 μm largest sphere, morphological split with 0.5 μm seed point diameter). Analysis was done per cell by manually selecting regions covering the individual cell and not overlapping with neighboring cells.

### Fluorescence live cell imaging

U2OS cells were seeded on a 35 mm WiLLCo-dish (WILLCO WELLS, HBST-3522) and incubated in growing conditions overnight. Before imaging, the cells were stained with LysoTracker™ Red DND-99 (1:1000 Thermo Fisher Scientific, L7528) for 30 minutes in growing conditions. Compound EL256 was added, followed by the start of imaging. Live-cell imaging was performed using a Lattice Lightsheet 7 microscope (Carl Zeiss Ltd.) under controlled conditions (37°C, 5% CO₂). LysoTracker and EL256’s signals were simultaneously captured in two channels with a dual-camera setup, using bandpass filters of 505–545 nm and 575–618 nm. Excitation was achieved with a combination of 488 nm and 561 nm lasers, using a Sinc3 15 × 650 light sheet. Imaging volumes were acquired with 1001 slices at a 0.2 µm step size and a pixel size of 0.145 µm. Time-lapse recordings were taken at 30-minute intervals over 24 hours. Data processing, including deskewing, cropping, and orthogonal projection, was performed in ZEN Blue 3.10 software, while final visualization and movie export were completed using Imaris 10.2 (Bitplane). The LysoTracker signal was used as a reference for cell location and focus during imaging and analysis, and its signal was not reported in this work.

## Acknowledgments

Research in the London Lab is supported by funding from the European Research Council (ERC_CoG 101125683), the Honey and Dr. Barry Sherman Lab, Dr. Barry Sherman Institute for Medicinal Chemistry, Abisch-Frenkel RNA Therapeutics Center, Moross Integrated Cancer Center, Goldhirsh-Yellin Foundation and Celia Zwillenberg-Fridman. E.D.L. acknowledges support from the European Research Council under the European Union’s Horizon 2020 research and innovation program (grant agreement No. 819318). Microscopy images in this paper were acquired at the de Picciotto Cancer Cell Observatory In memory of Wolfgang and Ruth Lesser of the Moross Integrated Cancer Center in the Department of Life Science Core Facilities, Weizmann Institute of Science

