## Supplemental Figures and Chemistry for "A pharmacological modality to sequester homomeric proteins"

### Supplementary Figures

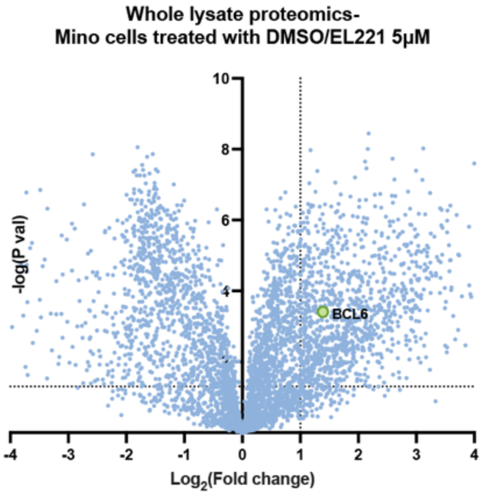


**Figure S1.** Global proteomics analysis of the cell's soluble lysate following treatment with 5μM EL221, comapred to DMSO at a 20h treatment, showing BCL6 as one of the significantly downregulated targets. The dotted lines represent a 2-fold reduction in protein level and *p*=0.05 significance.

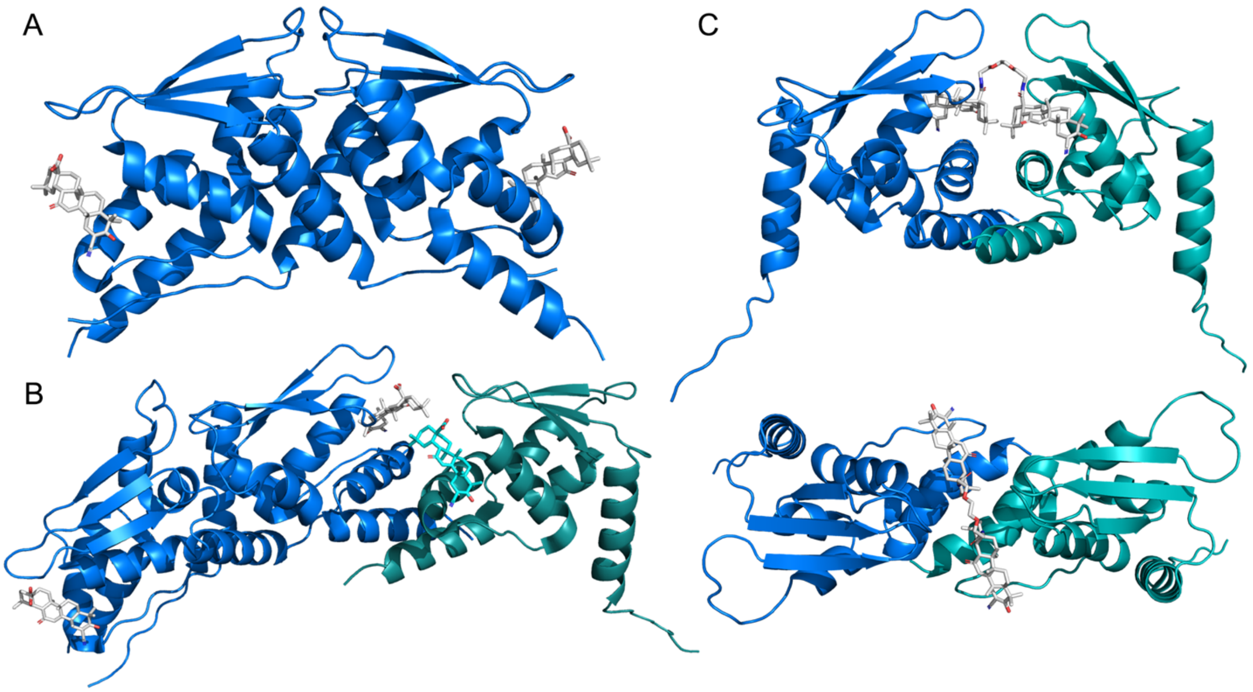


**Figure S2. A.** Keap1-BTB biological homodimeric unit in complex with Bardoxolone (in white sticks). **B.** Keap1-BTB biological homodimeric unit (blue), and an additional crystalographic symmetry mate monomeric Keap1-BTB (teal). **C.** EL133 modeled onto Keap1-BTB crystalographic interface originating from separate biological dimers.

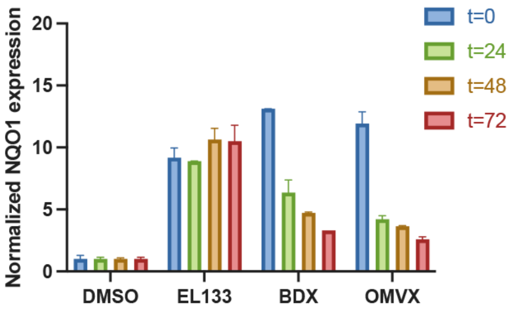


**Figure S3.** rt-PCR analysis of NQO1 expression after a 20h treatment of OCI-AML2 cells with DMSO, EL133, BDX, or OMVX at 250 nM, followed by washing of the cells (representative results following 2 independent experiment replicates).

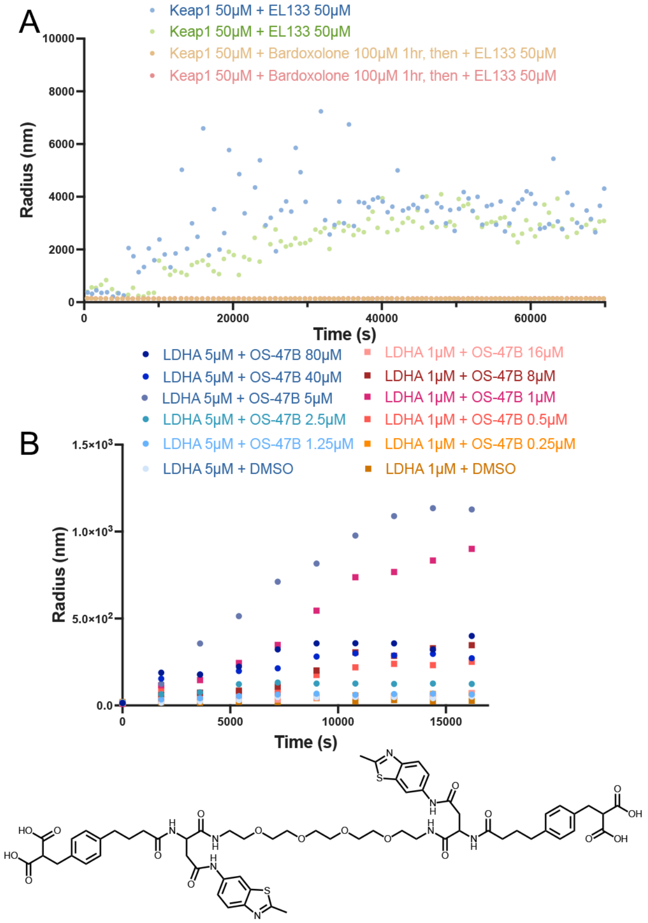


**Figure S4. A.** Dynamic light scattering experiments following recombinant Keap1-BTB and EL133 over time at room temperature, with and without pre-incubation of the protein with excess BDX. **B.** Dynamic light scattering experiments following recombinant LDHA and its PINCH at different ratios over time at room temperature.

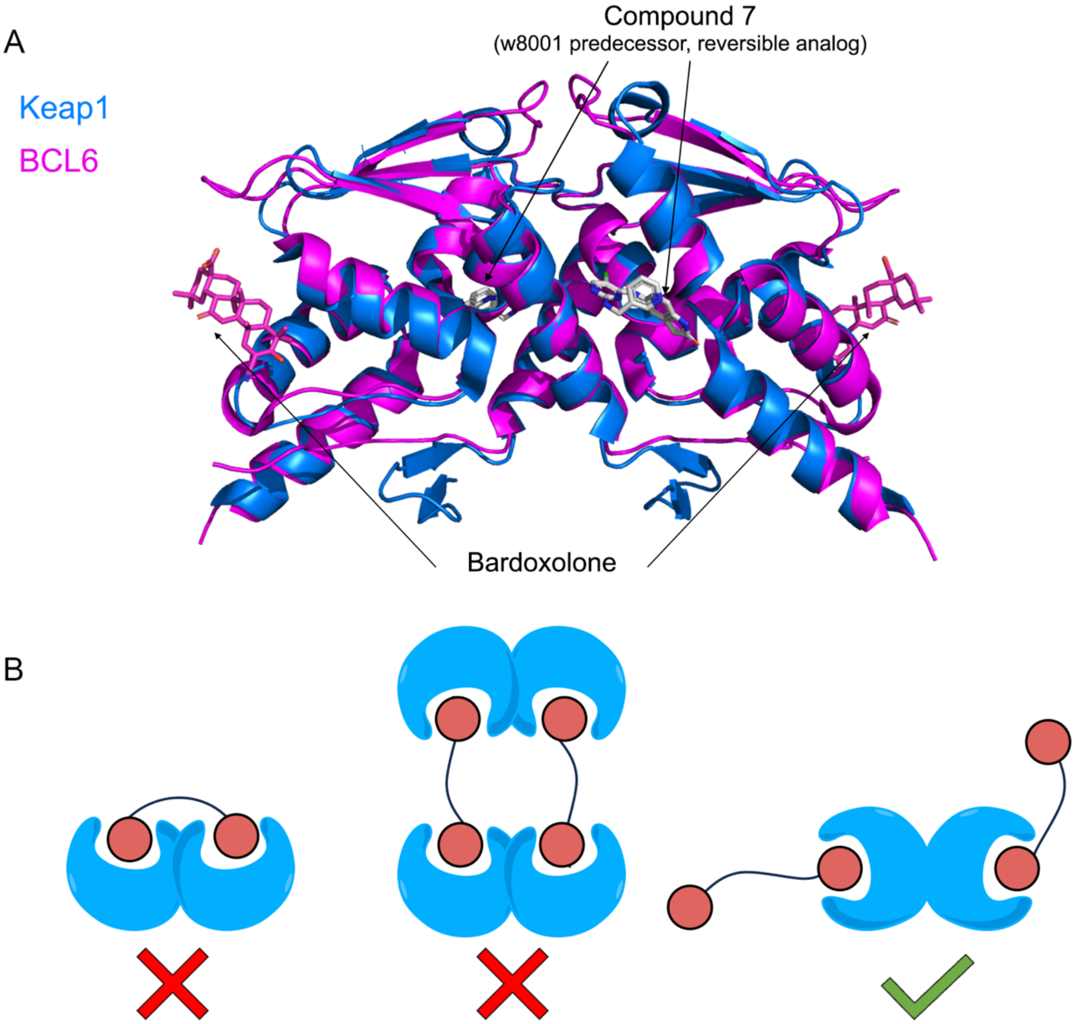


**Figure S5. A.** Overlay of BTB domains of Keap1 (blue, PDB: 4CXT) and BCL6 (magenta, PDB: 5X4Q) and their respective ligands. Compound 7[1] is an early reversible BCL6 binder from which w8001's covalent analog was later optimized, and refelcts our BCL6-PINCH binding site. This show the ligands bind distinct pockets on the BTB domain. However, in both cases, the pockets are on opposite sides of the dimer (left and right for Keap1, front and back for BCL6) **B.** Geometric constraits for effective polymerization by PINCHs. If the binding sites are too close to one another (left panel) or facing the same side of the multimer (middle panel) they may results in a closed oligomer formation. If the binding sites are on opposite sides (right panel) as they are in Keap1 and BCL6 (panel A), they should lead to effective polymerization.

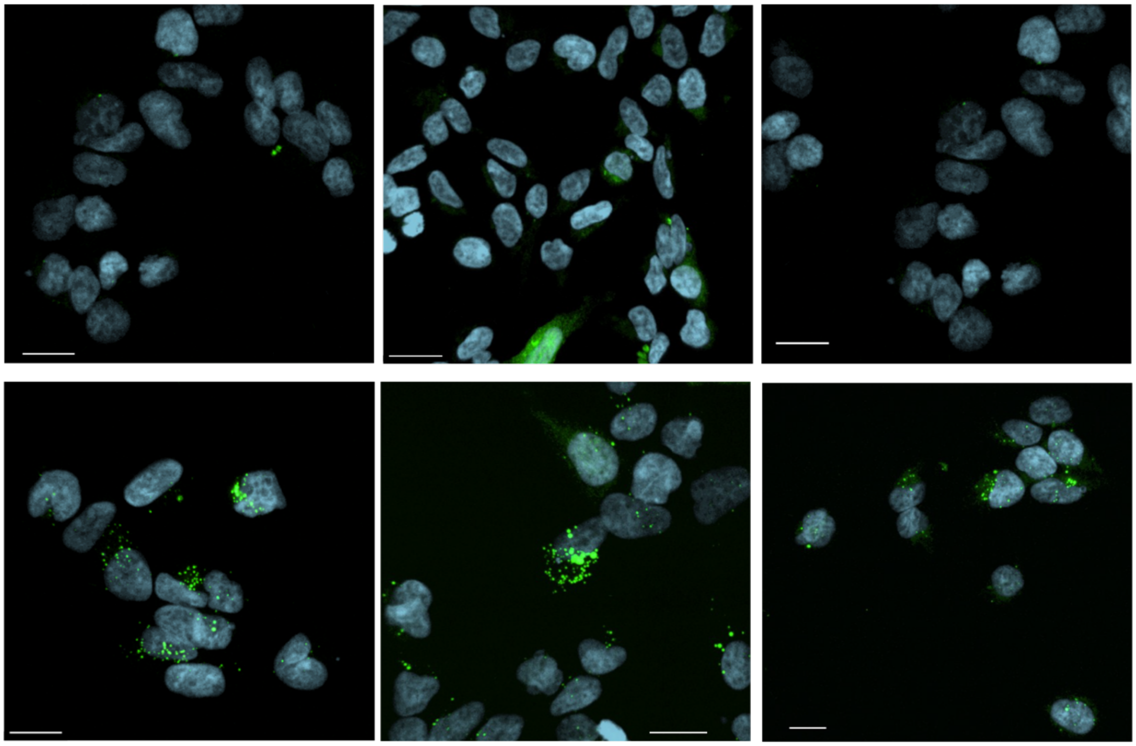

**Figure S6.** Representative images of HEK-293 cells overexpressed with BCL6-GFP(1-275), treated with w8001 (top) or EL221 (bottom), both at 5 μM. Scale bars are at 15 μm.

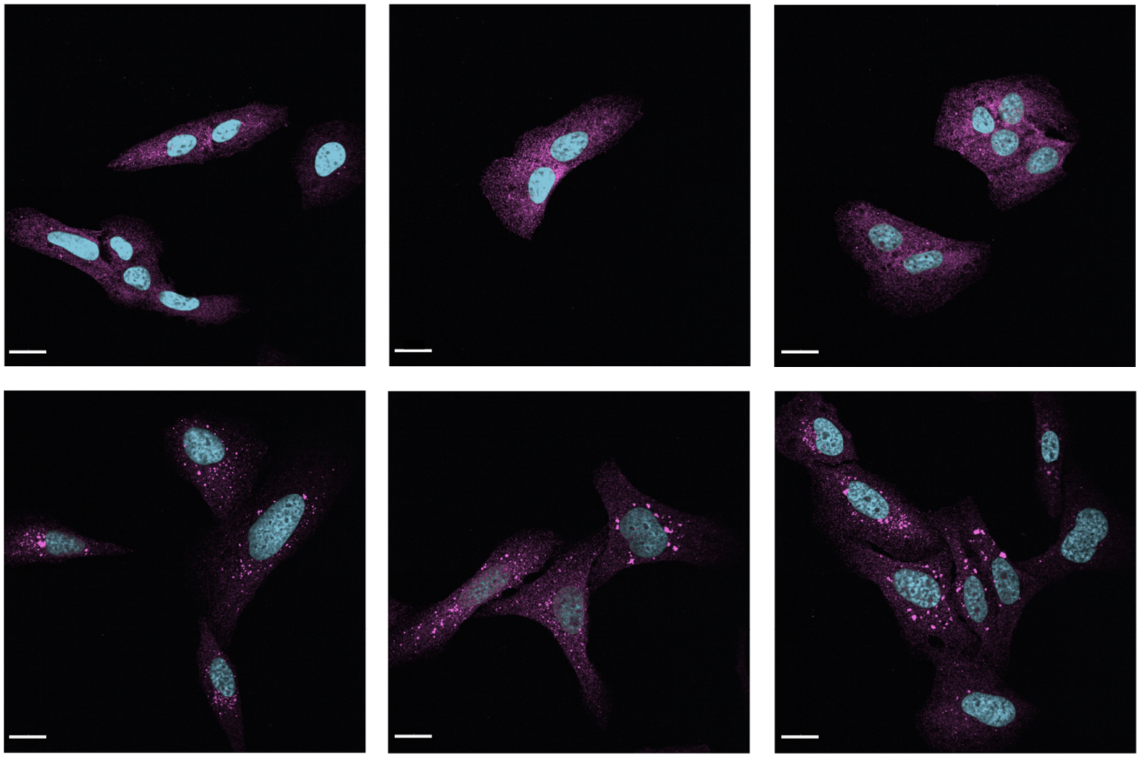

**Figure S7.** Representative images of U-2OS cells treated with Bardoxolone (top) or EL133 (bottom), both at 250 nM, immunostained with Keap1 antibody (Magenta). Scale bars are at 10 μm.

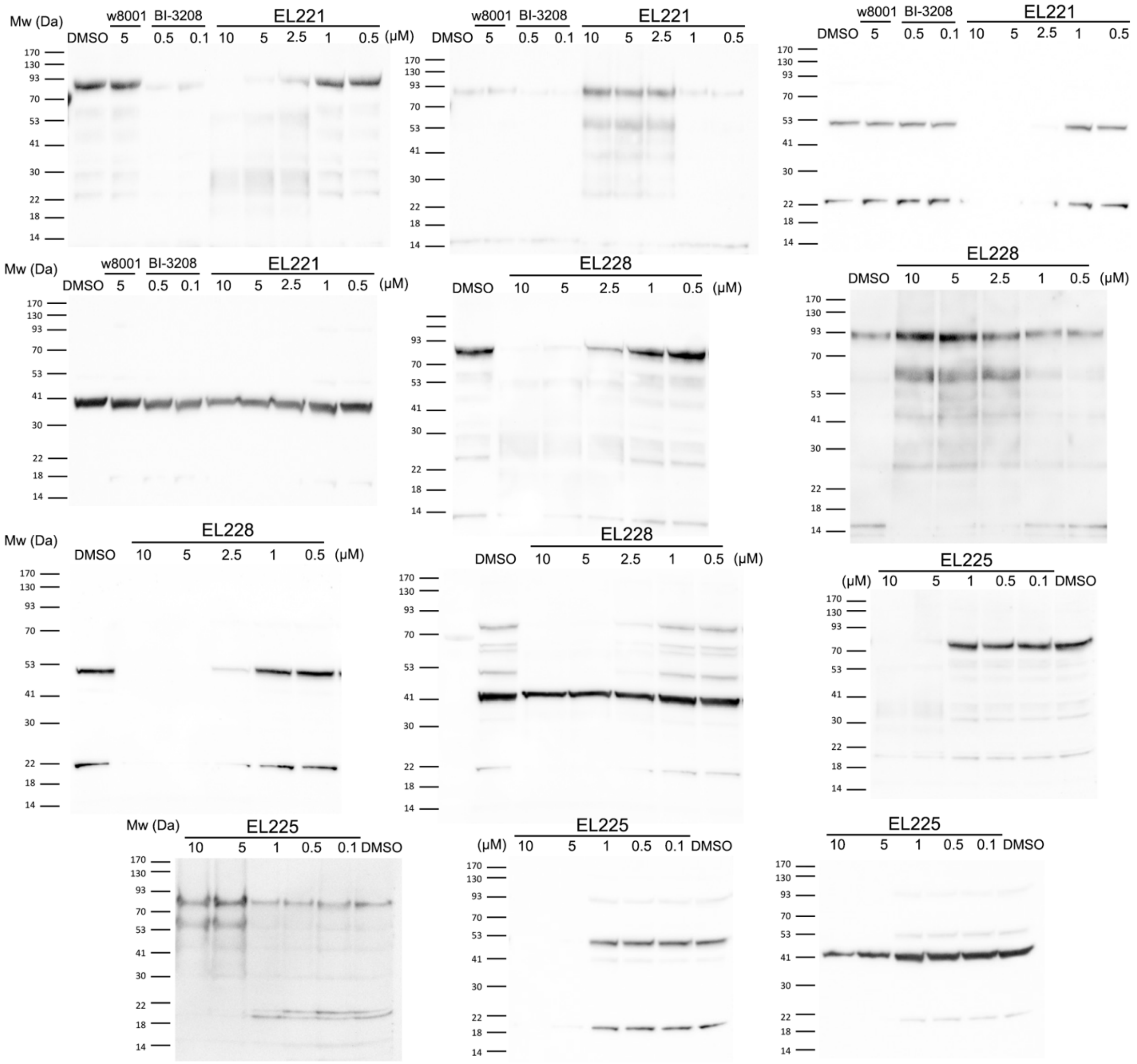


**Figure S8.** Full gels presented in Fig. 1 of the main text.

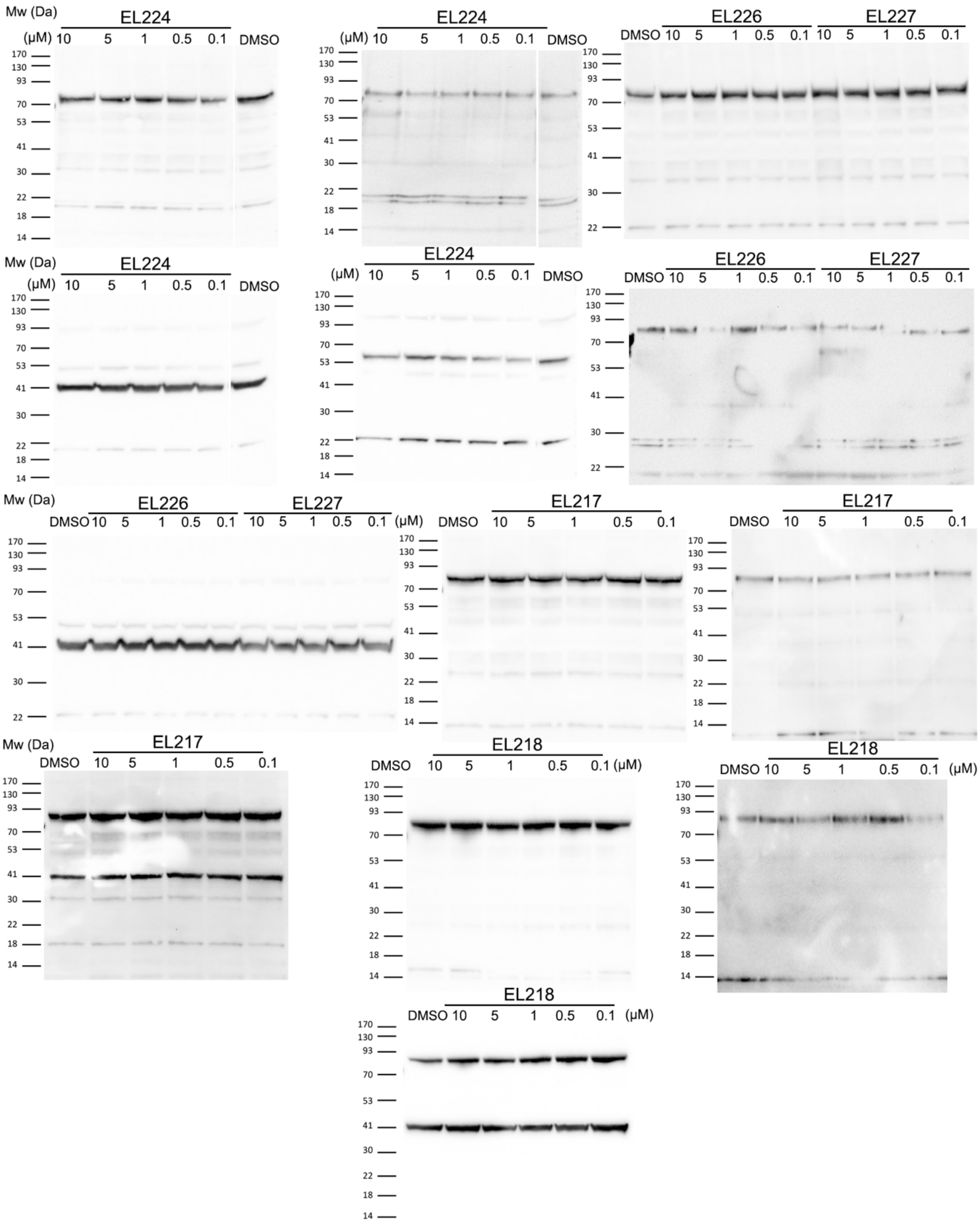


**Figure S9.** Full gels presented in Extended Data Fig. 1

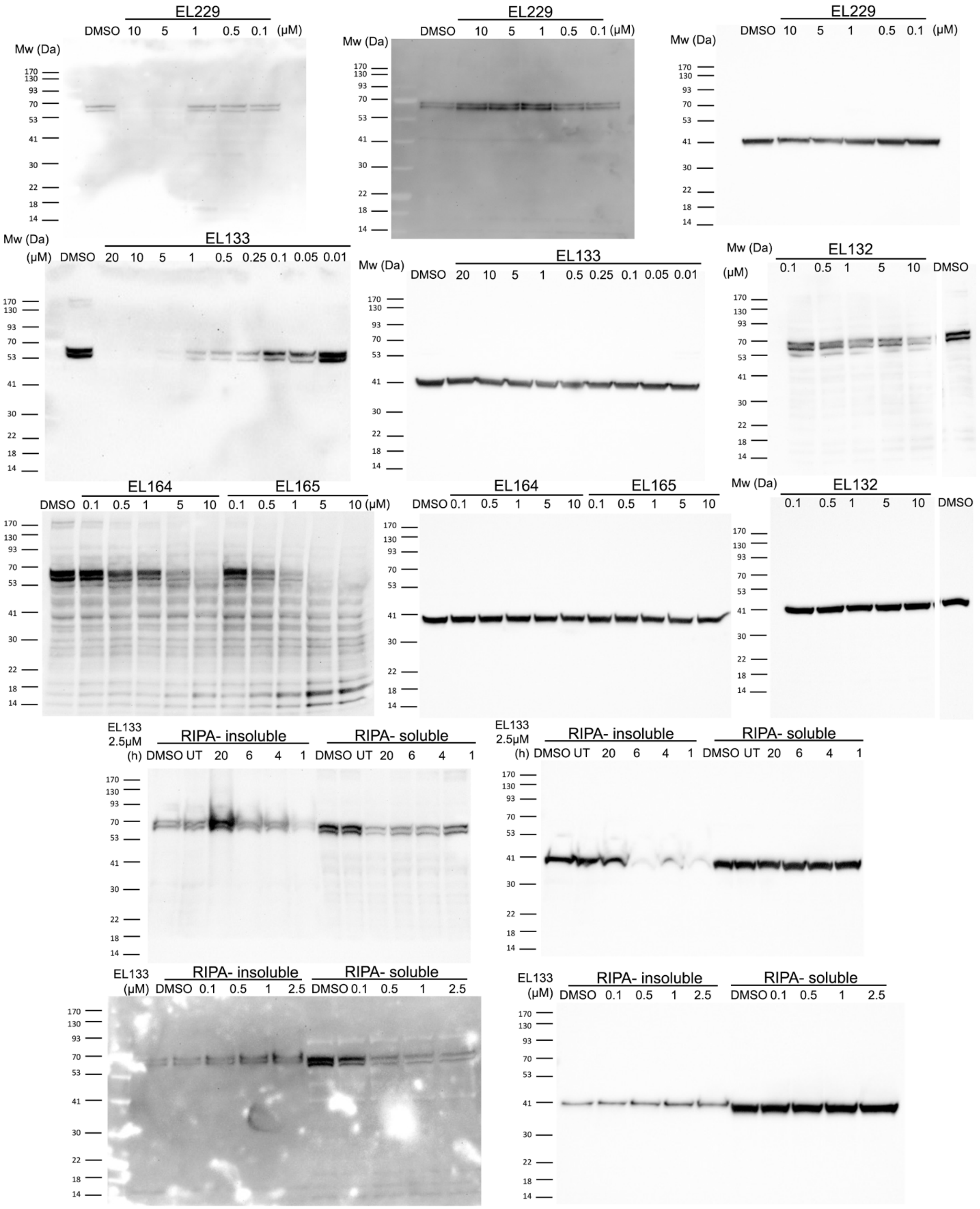


**Figure S10.** Full gels presented in Fig. 2 and Extended Data Fig. 2 of the main text.

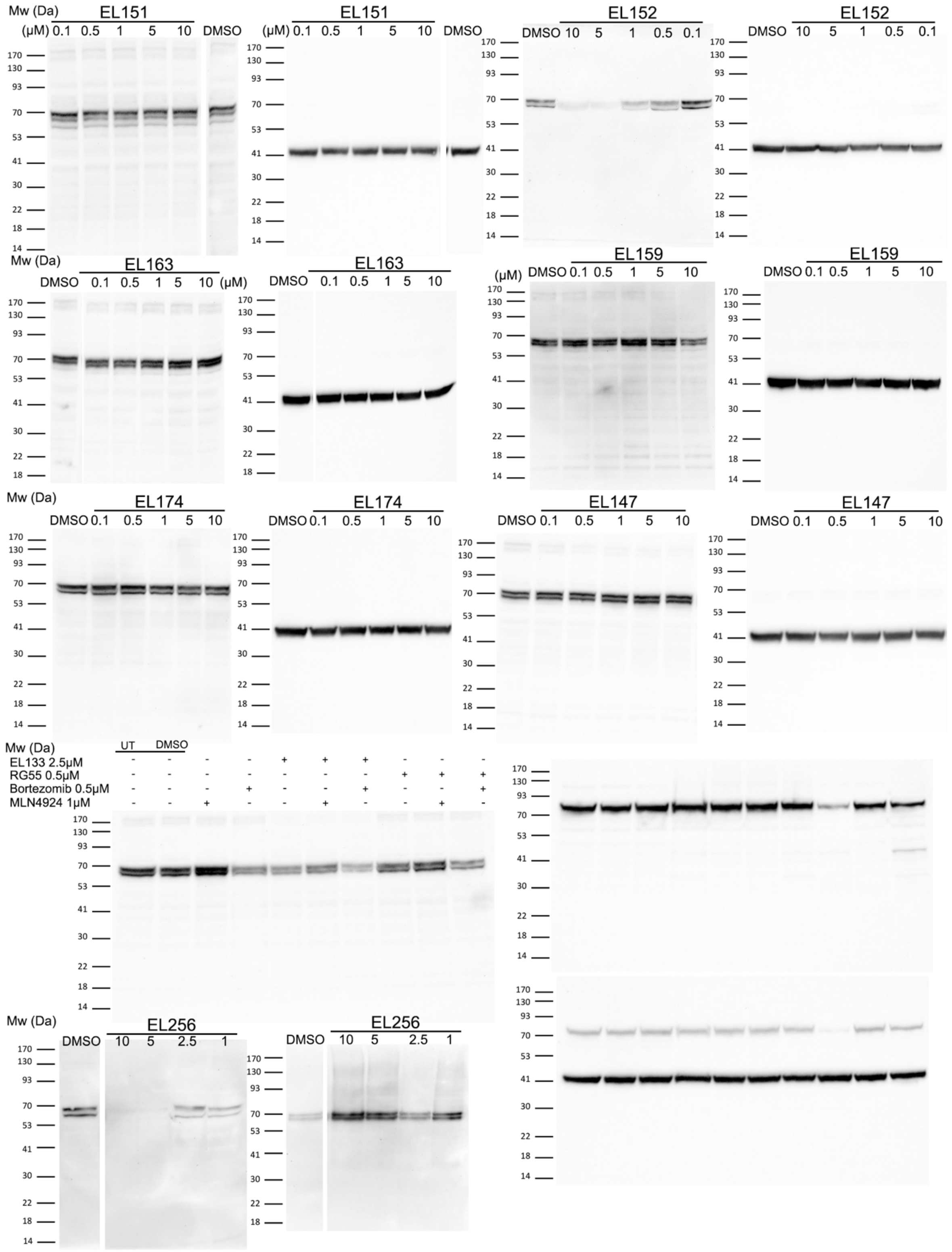


**Figure S11.** Full gels presented in Extended Data Fig. 2

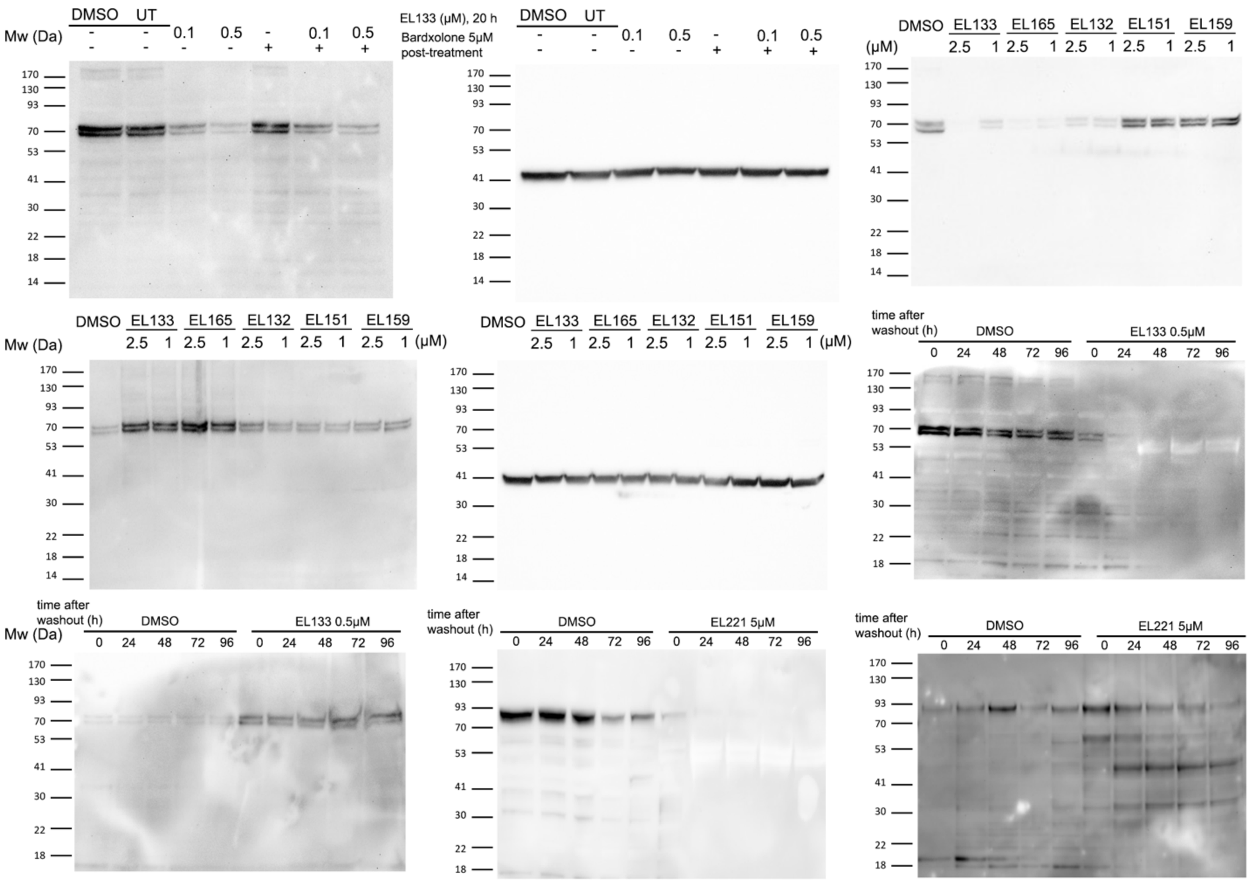


**Figure S12.** Full gels presented in Fig. 3 And Extended Data Fig. 3.

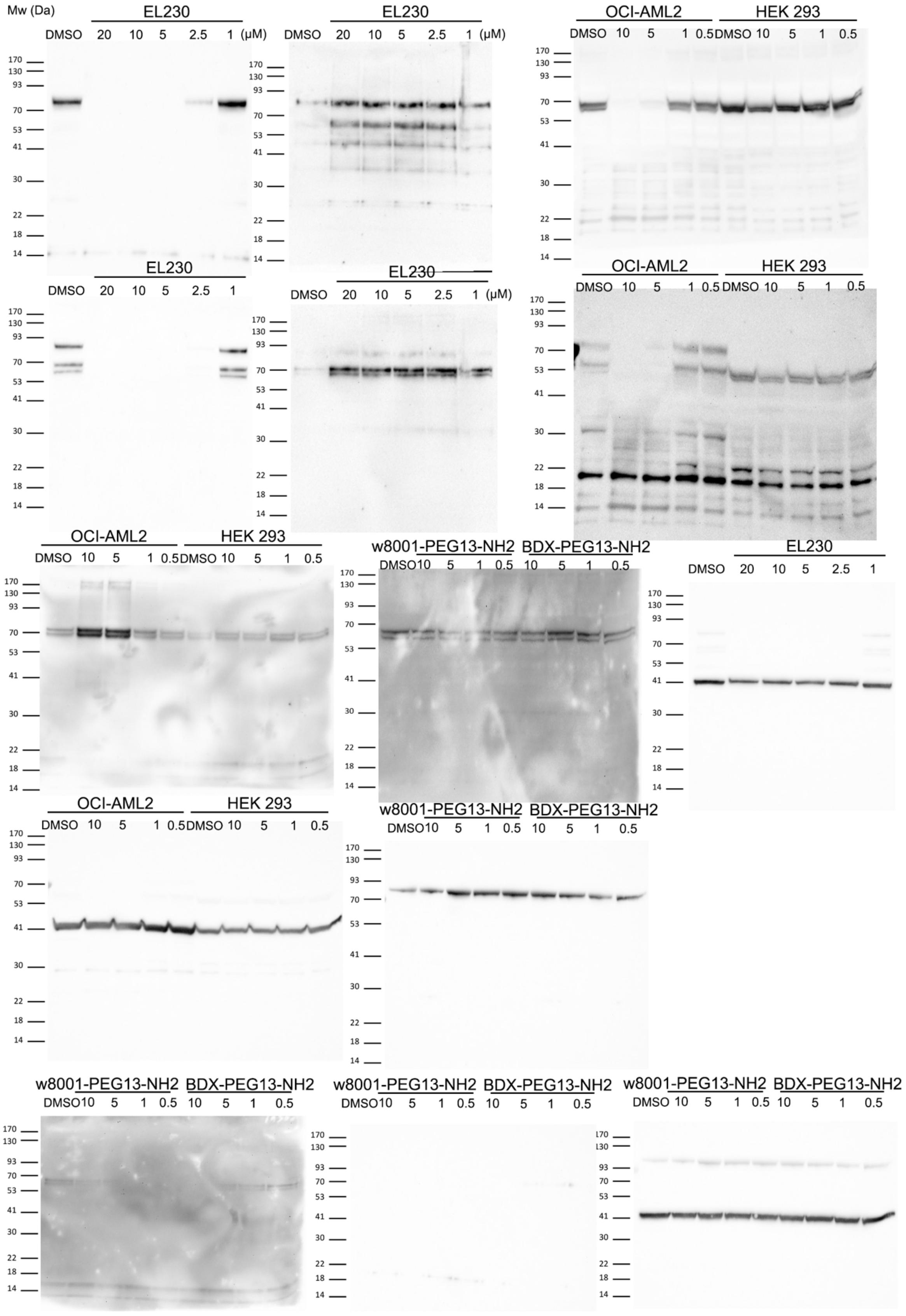


**Figure S13.** Full gels presented in Fig. 4

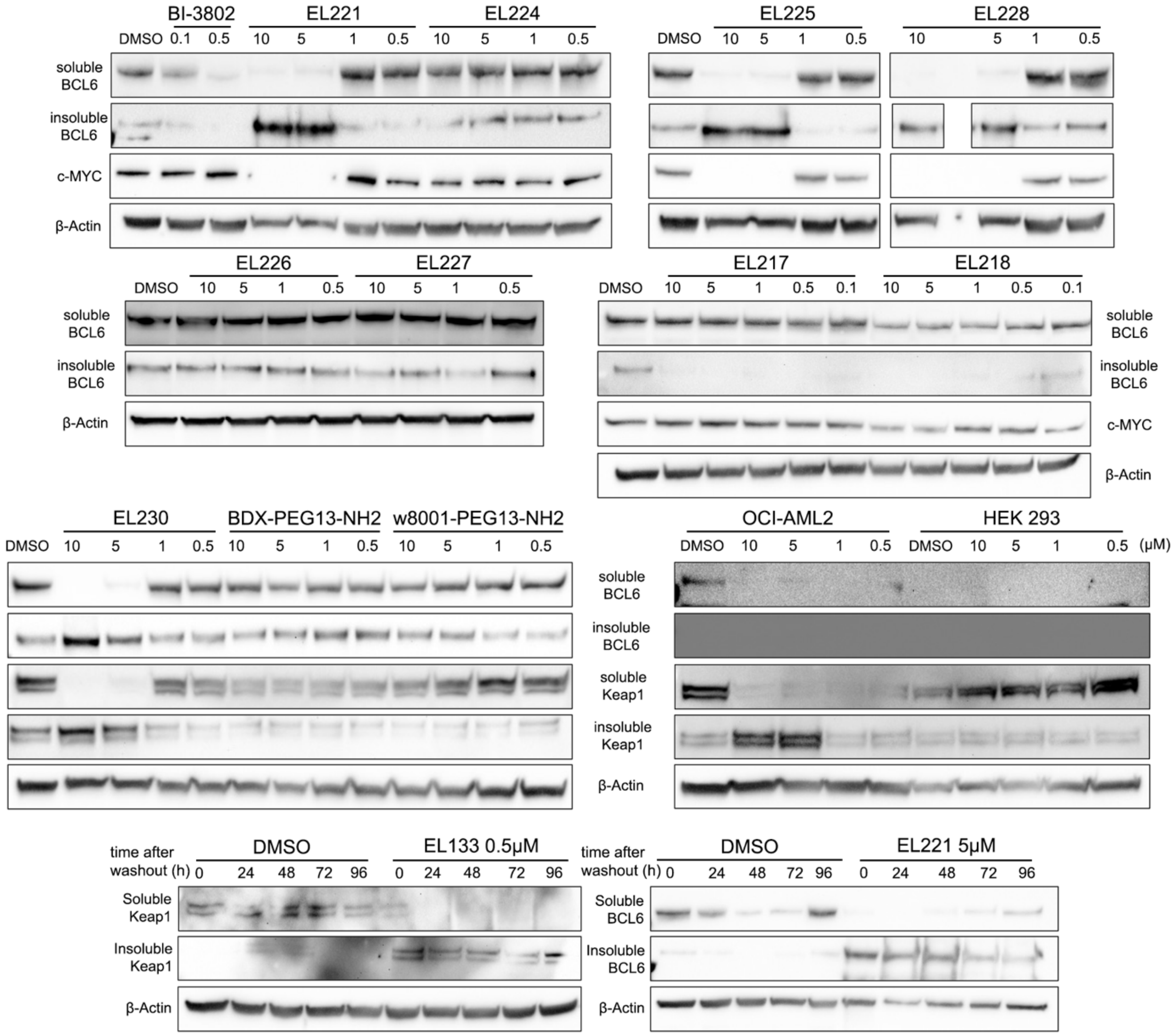


**Figure S14.** Representative repeats of the WBs shown in figures 1, 4 and extended data figures 1, 3.

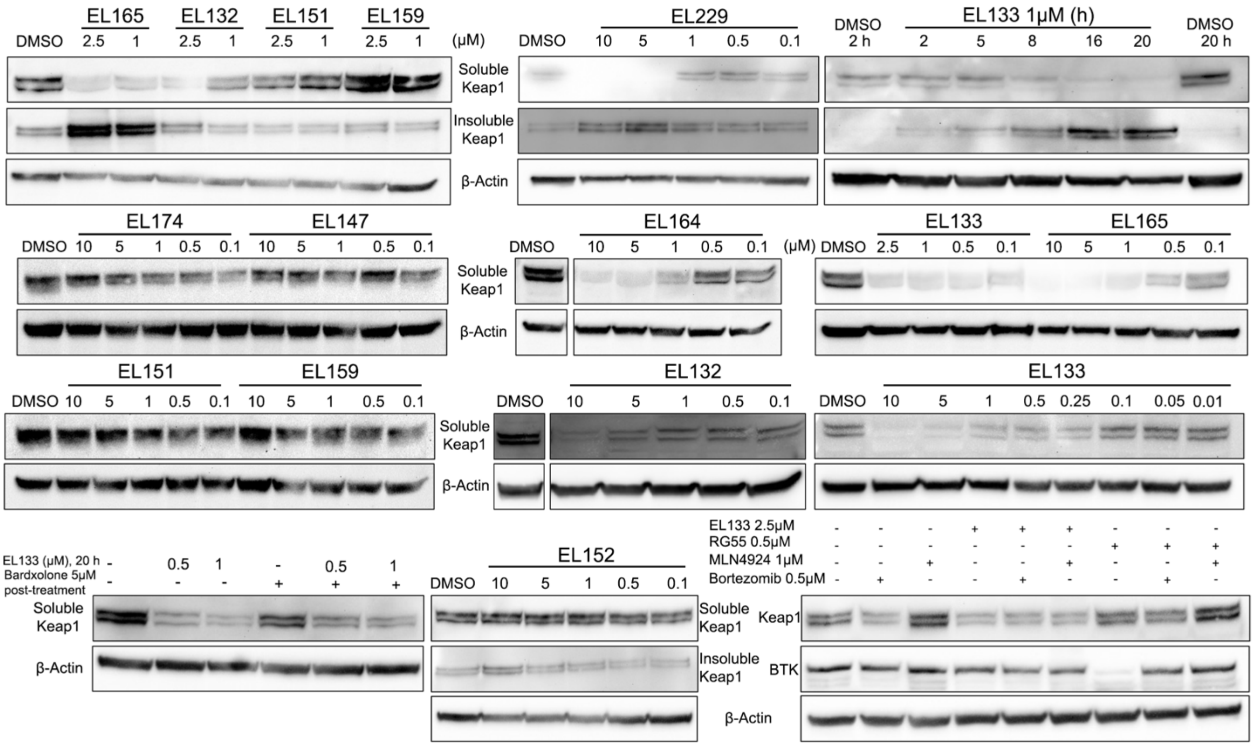


**Figure S15.** Representative repeats of the WBs shown in figures 2, 3 and extended data figure 2.

### Supplementary Note (synthetic procedures and analytical data)

**EL229** - (4a*S*,4a'*S*,6a*R*,6b*S*,6a'*R*,6b'*S*,8a*R*,8a'*R*,12a*S*,12a'*S*,14a*R*,14b*S*,14a'*R*,14b'*S*)-*N*a,*N*a'-(3,6,9,12,15,18,21,24,27,30,33,36,39-tridecaoxahentetracontane-1,41-diyl)bis(11-cyano-2,2,6a,6b,9,9,12a-heptamethyl-10,14-dioxo-1,3,4,5,6,6a,6b,7,8,8a,9,10,12a,14,14a,14b-hexadecahydropicene-4a(2*H*)-carboxamide)

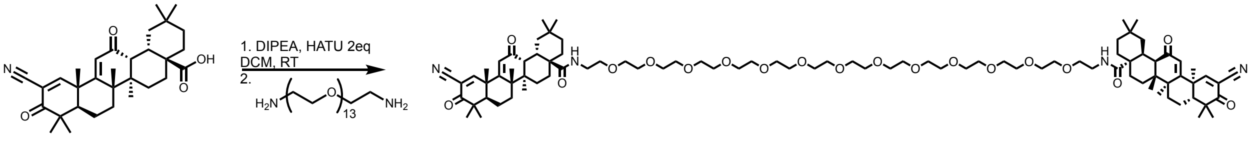


HATU-activation of Bardoxolone: 5 mg of Bardoxolone-COOH were dissolved in DCM, followed by 4eq of DIPEA and 2eq of HATU, and left stirring overnight at room temperature. Then, saturated $NH_4Cl$ solutions was added to the reaction vile to approximately double the volume, followed by 3 times extraction of the activated Bardoxolone acid using DCM. The DCM was then evaporated, and the product dissolved in 200 µL DMF.
To the activated Bardoxolone, 5eq of DIPEA were added, followed by addition of 3,6,9,12,15,18,21,24,27,30,33,36,39-tridecaoxahentetracontane-1,41-diamine (PEG13-diamine), and dissolved in DCM. After overnight reaction in room temperature, the solvent was evaporated, the crude reaction was then dissolved in 50% ACN in $H_2O$ and purified using HPLC.

$^1$H NMR (500 MHz, DMSO) δ 8.65 (s, 1H), 7.79 – 7.69 (m, 1H), 6.20 (s, 1H), 3.53 – 3.47 (m, 30H), 3.43 – 3.39 (m, 3H), 3.17 – 3.09 (m, 2H), 3.02 (dd, *J* = 22.9, 4.5 Hz, 1H), 2.89 – 2.81 (m, 1H), 1.91 – 1.79 (m, 2H), 1.65 (d, *J* = 7.6 Hz, 3H), 1.58 – 1.52 (m, 3H), 1.43 (s, 3H), 1.24 (d, *J* = 6.9 Hz, 3H), 1.17 (d, *J* = 5.4 Hz, 3H), 1.06 (d, *J* = 7.6 Hz, 3H), 0.93 (s, 3H), 0.89 (s, 3H), 0.86 (s, 3H).
$^{13}$C NMR (126 MHz, DMSO) δ 200.00, 197.66, 177.15, 169.25, 168.65, 123.87, 115.61, 113.27, 70.25, 70.00, 69.50, 48.93, 45.96, 45.86, 45.81, 45.54, 44.84, 43.02, 41.99, 41.68, 39.23, 36.01, 34.69, 33.75, 31.24, 30.73, 30.70, 27.92, 26.58, 26.33, 24.71, 24.25, 23.61, 23.29, 22.33, 21.77, 21.58.
HR-MS (m/z): Calculated: 1579.00; Found: 1601.9918 [M+Na]+.

**EL165**- (4a*S*,4a'*S*,6a*R*,6b*S*,6a'*R*,6b'*S*,8a*R*,8a'*R*,12a*S*,12a'*S*,14a*R*,14b*S*,14a'*R*,14b'*S*)-*N*a,*N*a'-(piperazine-1,4-diylbis(ethane-2,1-diyl))bis(11-cyano-2,2,6a,6b,9,9,12a-heptamethyl-10,14-dioxo-1,3,4,5,6,6a,6b,7,8,8a,9,10,12a,14,14a,14b-hexadecahydropicene-4a(2*H*)-carboxamide)

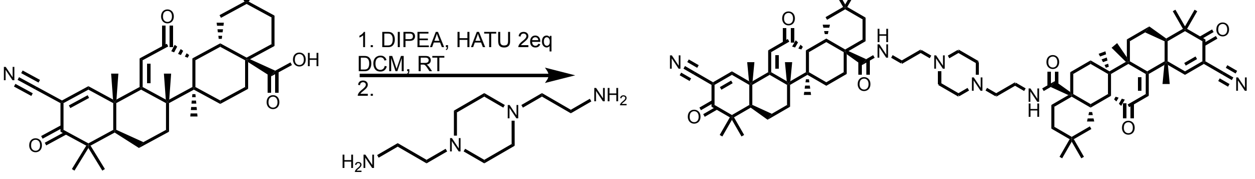


Synthetic procedure performed similar to the one described for EL229, using 2,2'-(piperazine-1,4-diyl)bis(ethan-1-amine) as the linker in the second step.

$^{1}H$ NMR (500 MHz, $CDCl_3$) δ 8.08 (d, *J* = 10.8 Hz, 1H), 8.03 (s, 1H), 6.01 (s, 1H), 3.83 (s, 1H), 3.55 (d, *J* = 4.6 Hz, 4H), 3.15 (d, *J* = 16.1 Hz, 2H), 3.03 (d, *J* = 3.7 Hz, 1H), 2.96 (d, *J* = 12.8 Hz, 1H), 2.03 (t, *J* = 12.7 Hz, 2H), 1.70 (ddd, *J* = 54.5, 26.4, 15.0 Hz, 9H), 1.52 (s, 3H), 1.38 (dd, *J* = 10.7, 8.3 Hz, 2H), 1.33 (s, 3H), 1.28 (s, 3H), 1.19 (s, 3H), 1.04 (d, *J* = 6.6 Hz, 3H), 1.00 (s, 3H), 0.93 (s, 3H).
$^{13}C$ NMR (126 MHz, $CDCl_3$) δ 199.31, 196.57, 178.45, 165.79, 161.95, 124.08, 114.57, 114.46, 57.93, 49.86, 49.47, 47.68, 46.64, 45.83, 45.02, 42.59, 42.02, 35.90, 34.59, 33.56, 33.26, 31.70, 31.62, 30.59, 29.71, 27.80, 26.96, 26.55, 24.77, 23.16, 23.06, 22.61, 21.69, 21.57, 18.22.
HR-MS (m/z): Calculated: 1658.52; Found: 1119.7657 [M+H]+.

**EL133**- (4a*S*,4a'*S*,6a*R*,6b*S*,6a'*R*,6b'*S*,8a*R*,8a'*R*,12a*S*,12a'*S*,14a*R*,14b*S*,14a'*R*,14b'*S*)-*N*a,*N*a'-((ethane-1,2-diylbis(oxy))bis(ethane-2,1-diyl))bis(11-cyano-2,2,6a,6b,9,9,12a-heptamethyl-10,14-dioxo-1,3,4,5,6,6a,6b,7,8,8a,9,10,12a,14,14a,14b-hexadecahydropicene-4a(2*H*)-carboxamide)

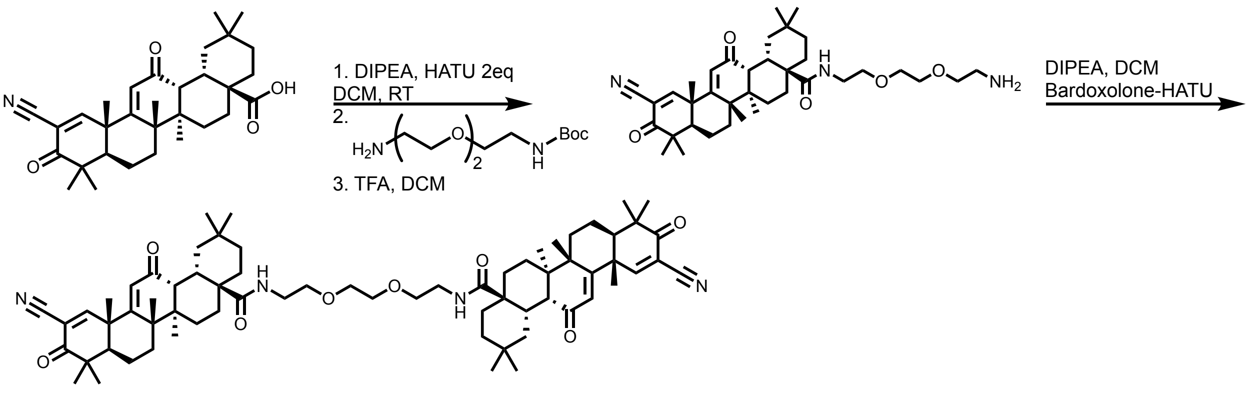


Bardoxolone-COOH was activated using HATU according to the synthetic procedure described for compound EL229, and it was reacted with *tert*-butyl (2-(2-(2-aminoethoxy)ethoxy)ethyl)carbamate in DCM overnight. TFA was added to the reaction to obtain a final solvent ratio of approximately 1:2 (TFA:DCM) and the reaction was left overnight at room temperature. The deprotected intermediate was then evaporated and purified by HPLC, lyophilized and was then used directly in the following reaction. Bardoxolone-HATU was added to the intermediate amine with DIPEA in DCM. After 6h, the reaction was evaporated and the final product was purified using HPLC.

$^{1}H$ NMR (500 MHz, $CDCl_3$) δ 8.27 (s, 1H), 6.64 (s, 1H), 6.17 (s, 1H), 3.60 – 3.48 (m, 5H), 3.03 (s, 1H), 2.91 (d, *J* = 10.3 Hz, 1H), 1.94 – 1.71 (m, 15H), 1.56 (s, 1H), 1.52 (s, 3H), 1.37 (s, 3H), 1.28 (s, 3H), 1.19 (s, 3H), 1.05 (s, 3H), 1.00 (s, 3H), 0.93 (s, 3H).
$^{13}C$ NMR (126 MHz, $CDCl_3$) δ 199.41, 196.56, 177.44, 169.40, 166.12, 124.14, 114.57, 114.46, 70.38, 69.65, 49.72, 47.66, 46.71, 45.91, 45.02, 42.65, 42.15, 39.39, 36.20, 34.63, 33.98, 33.32, 32.11, 31.75, 30.69, 29.70, 27.71, 26.97, 26.54, 24.88, 23.09, 22.58, 21.81, 21.58, 18.25.
HR-MS (m/z): Calculated: 1094.71.; Found: 1095.7185 [M+H]+, 1117.6956 [M+Na]+

**EL159**- (4a*S*,4a'*S*,6a*R*,6b*S*,6a'*R*,6b'*S*,8a*R*,8a'*R*,12a*S*,12a'*S*,14a*R*,14b*S*,14a'*R*,14b'*S*)-*N*a,*N*a'-(cyclohexane-1,4-diylbis(methylene))bis(11-cyano-2,2,6a,6b,9,9,12a-heptamethyl-10,14-dioxo-1,3,4,5,6,6a,6b,7,8,8a,9,10,12a,14,14a,14b-hexadecahydropicene-4a(2*H*)-carboxamide)

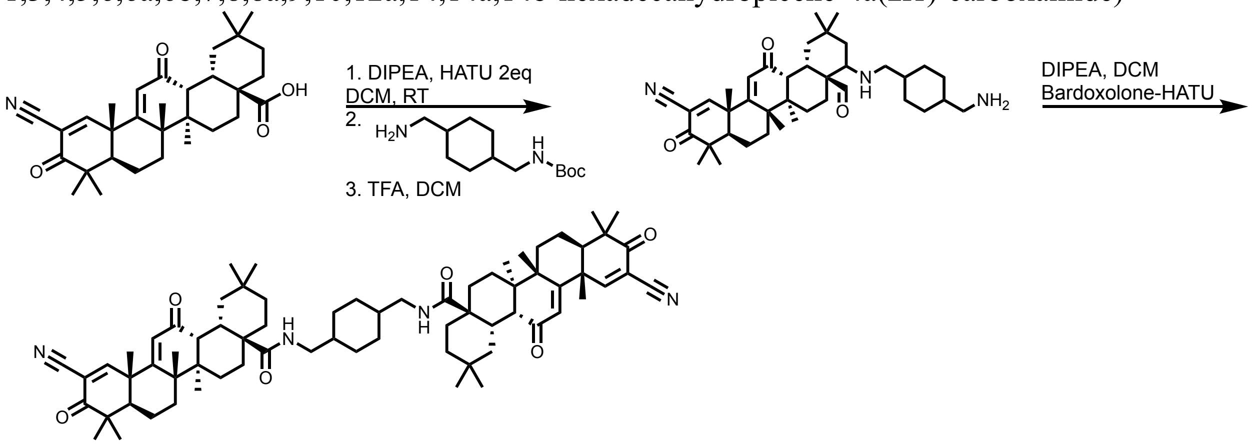


Synthetic procedure performed similar to the one described for EL133, using *tert*-butyl ((4-(aminomethyl)cyclohexyl)methyl)carbamate as the linker in the second step.

HR-MS (m/z): Calculated: 1088.73; Found: 1111.7249 [M+Na]+.

**EL152**- (4a*S*,4a'*S*,6a*R*,6b*S*,6a'*R*,6b'*S*,8a*R*,8a'*R*,12a*S*,12a'*S*,14a*R*,14b*S*,14a'*R*,14b'*S*)-*N*a,*N*a'-(3,6,9,12,15,18-hexaoxaicosane-1,20-diyl)bis(11-cyano-2,2,6a,6b,9,9,12a-heptamethyl-10,14-dioxo-1,3,4,5,6,6a,6b,7,8,8a,9,10,12a,14,14a,14b-hexadecahydropicene-4a(2*H*)-carboxamide)

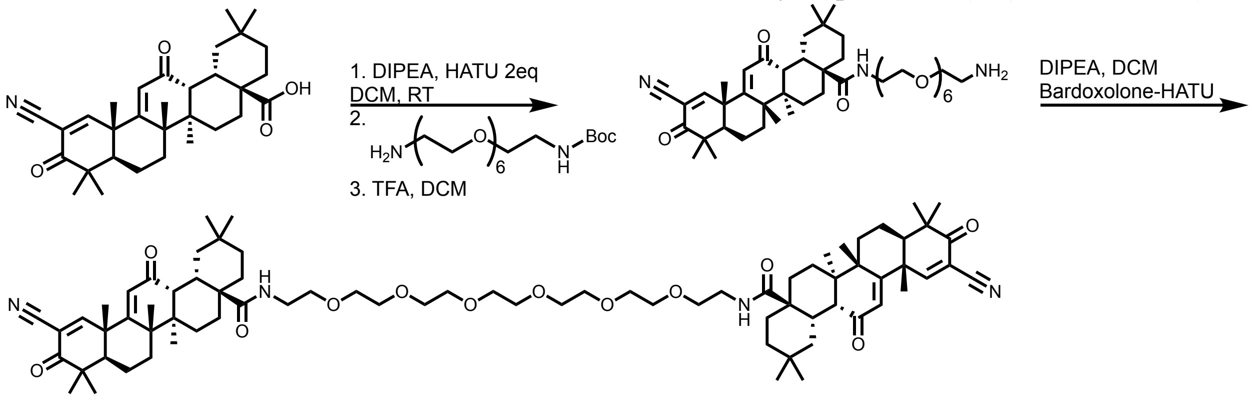


Synthetic procedure performed similar to the one described for EL133, using *tert-butyl (20-amino-3,6,9,12,15,18-hexaoxaicosyl)carbamate* as the linker in the second step.

HR-MS (m/z): Calculated: 1270.81.; Found: 1271.8190 [M+H]+, 1293.8079 [M+Na]+

**EL151**- (4a*S*,6a*R*,6b*S*,8a*R*,12a*S*,14a*R*,14b*S*)-11-cyano-*N*-(1-((4a*S*,6a*R*,6b*S*,8a*R*,12a*S*,14a*R*,14b*S*)-11-cyano-2,2,6a,6b,9,9,12a-heptamethyl-10,14-dioxo-1,3,4,5,6,6a,6b,7,8,8a,9,10,12a,14,14a,14b-hexadecahydropicen-4a(2*H*)-yl)-1-oxo-5,8,12-trioxa-

2-azatetradecan-14-yl)-2,2,6a,6b,9,9,12a-heptamethyl-10,14-dioxo-1,3,4,5,6,6a,6b,7,8,8a,9,10,12a,14,14a,14b-hexadecahydropicene-4a(2*H*)-carboxamide

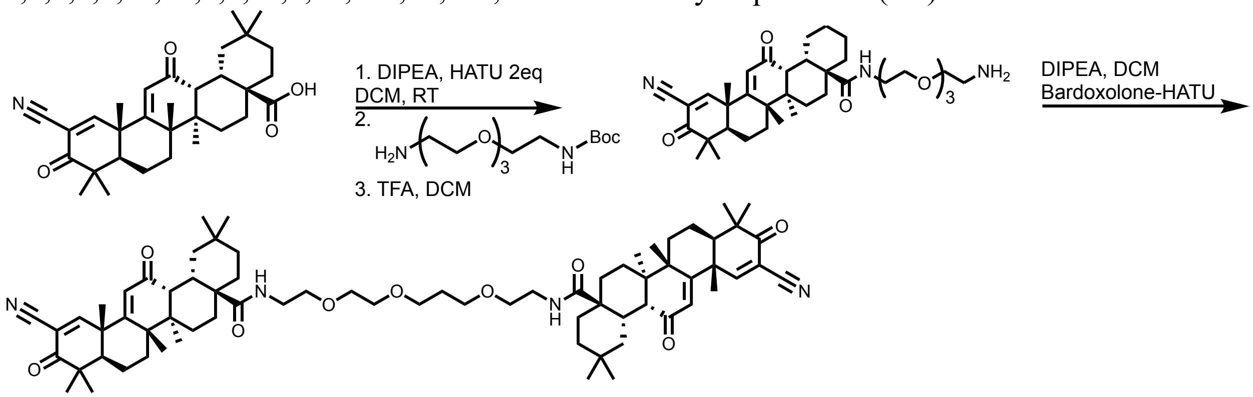


Synthetic procedure performed similar to the one described for EL133, using tert-butyl (2-(2-(3-(2-aminoethoxy)propoxy)ethoxy)ethyl)carbamate as the linker in the second step.

HR-MS (m/z): Calculated: 1138.73; Found: 1139.7429 [M+H]+, 1161.7247 [M+Na]+

**EL147**- (4a*S*,4a'*S*,6a*R*,6b*S*,6a'*R*,6b'*S*,8a*R*,8a'*R*,12a*S*,12a'*S*,14a*R*,14b*S*,14a'*R*,14b'*S*)-*N*a,*N*a'-(oxybis(ethane-2,1-diyl))bis(11-cyano-2,2,6a,6b,9,9,12a-heptamethyl-10,14-dioxo-1,3,4,5,6,6a,6b,7,8,8a,9,10,12a,14,14a,14b-hexadecahydropicene-4a(2*H*)-carboxamide)

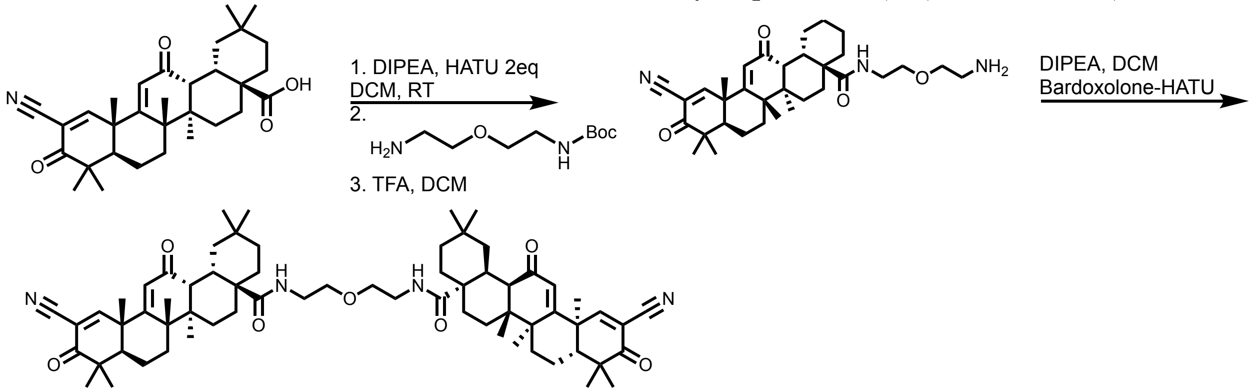


Synthetic procedure performed similar to the one described for EL133, using *tert-butyl (2-(2-aminoethoxy)ethyl)carbamate* as the linker in the second step.

HR-MS (m/z): Calculated: 1050.68; Found: 1051.6923 [M+H]+, 1073.6742 [M+Na]+

**EL164**- (4a*S*,4a'*S*,6a*R*,6b*S*,6a'*R*,6b'*S*,8a*R*,8a'*R*,12a*S*,12a'*S*,14a*R*,14b*S*,14a'*R*,14b'*S*)-*N*a,*N*a'-((ethane-1,2-diylbis(sulfanediyl))bis(ethane-2,1-diyl))bis(11-cyano-2,2,6a,6b,9,9,12a-heptamethyl-10,14-dioxo-1,3,4,5,6,6a,6b,7,8,8a,9,10,12a,14,14a,14b-hexadecahydropicene-4a(2*H*)-carboxamide)

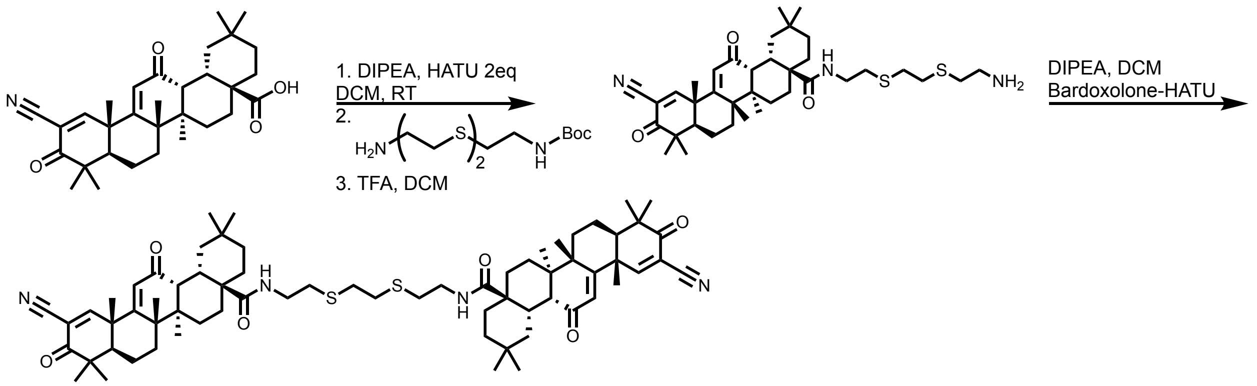


Synthetic procedure performed similar to the one described for EL133, using *tert-butyl (2-((2-((2-aminoethyl)thio)ethyl)thio)ethyl)carbamate* as the linker in the second step.

[1]H NMR (500 MHz, DMSO) δ 8.65 (s, 1H), 7.84 (t, *J* = 5.2 Hz, 1H), 6.20 (s, 1H), 3.36 (dd, *J* = 12.3, 5.7 Hz, 1H), 3.30 (s, 1H), 3.21 – 3.13 (m, 1H), 3.05 (d, *J* = 4.4 Hz, 1H), 2.85 (d, *J* = 13.3 Hz, 1H), 2.71 (s, 2H), 2.66 – 2.57 (m, 2H), 1.94 – 1.76 (m, 3H), 1.61 (ddd, *J* = 21.3, 17.4, 8.2 Hz, 5H), 1.44 (d, *J* = 3.1 Hz, 1H), 1.43 (s, 3H), 1.25 (d, *J* = 6.0 Hz, 3H), 1.17 (s, 3H), 1.06 (s, 3H), 0.94 (s, 3H), 0.88 (d, *J* = 8.8 Hz, 3H), 0.86 (s, 3H).
[13]C NMR (126 MHz, DMSO) δ 199.67, 197.79, 177.13, 169.45, 168.83, 123.86, 115.58, 113.27, 49.16, 46.68, 45.89, 45.87, 44.84, 43.02, 42.02, 40.59, 39.17, 35.81, 34.61, 33.71, 33.60, 31.66, 31.37, 31.22, 31.02, 30.69, 27.79, 26.58, 26.35, 24.84, 23.63, 22.36, 21.77, 21.61, 17.92.
HR-MS (m/z): Calculated: 1126.66; Found: 1127.6669 [M+H]+, 1149.6544 [M+Na]+

**EL132**- ((4a*S*,4a'*S*,6a*R*,6b*S*,6a'*R*,6b'*S*,8a*R*,8a'*R*,12a*S*,12a'*S*,14a*R*,14b*S*,14a'*R*,14b'*S*)-*N*a,*N*a'-(hexane-1,6-diyl)bis(11-cyano-2,2,6a,6b,9,9,12a-heptamethyl-10,14-dioxo-1,3,4,5,6,6a,6b,7,8,8a,9,10,12a,14,14a,14b-hexadecahydropicene-4a(2*H*)-carboxamide)

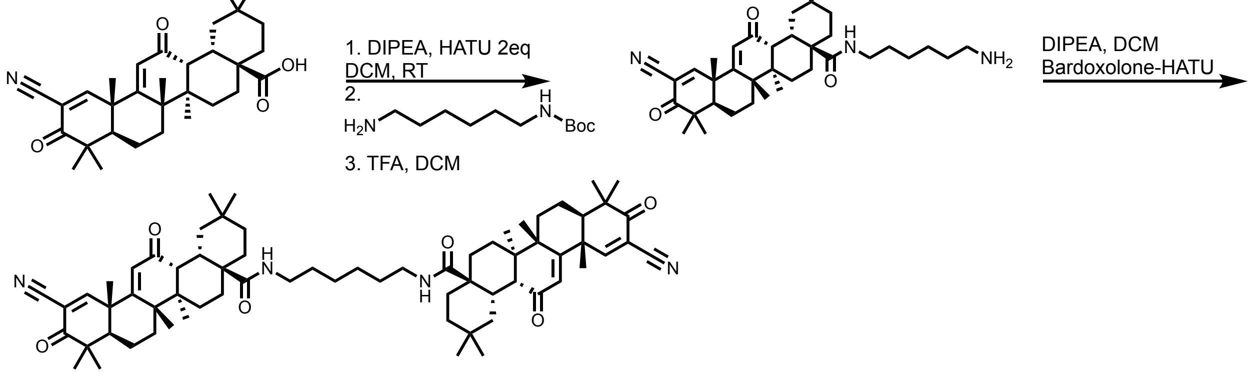


Synthetic procedure performed similar to the one described for EL133, using *tert-butyl (6-aminohexyl)carbamate* as the linker in the second step.

[1]H NMR (500 MHz, $CDCl_3$) δ 8.07 (s, 1H), 5.99 (s, 1H), 5.97 (t, *J* = 5.9 Hz, 1H), 3.26 (qd, *J* = 13.3, 6.4 Hz, 2H), 3.05 (d, *J* = 4.4 Hz, 1H), 2.92 – 2.85 (m, 1H), 1.82 – 1.78 (m, 4H), 1.60 (s,

14H), 1.51 (s, 3H), 1.36 – 1.33 (m, $J$ = 9.7 Hz, 4H), 1.28 (s, 3H), 1.19 (s, 3H), 1.04 (s, 3H), 1.02 (s, 3H), 0.93 (s, 3H).
$^{13}$C NMR (126 MHz, $CDCl_3$) δ 199.04, 196.52, 177.13, 168.91, 165.63, 123.97, 114.65, 114.37, 49.58, 47.71, 46.51, 45.92, 45.03, 42.56, 42.10, 39.16, 36.12, 34.61, 34.14, 33.27, 32.05, 31.69, 30.64, 29.73, 27.77, 27.00, 26.64, 26.14, 24.91, 23.13, 22.92, 21.78, 21.57, 18.25.
HR-MS (m/z): Calculated: 1062.72; Found: 1085.7102 [M+H]+

**EL163**- (4a*S*,4a'*S*,6a*R*,6b*S*,6a'*R*,6b'*S*,8a*R*,8a'*R*,12a*S*,12a'*S*,14a*R*,14b*S*,14a'*R*,14b'*S*)-*N*a,*N*a'-(octane-1,8-diyl)bis(11-cyano-2,2,6a,6b,9,9,12a-heptamethyl-10,14-dioxo-1,3,4,5,6,6a,6b,7,8,8a,9,10,12a,14,14a,14b-hexadecahydropicene-4a(2*H*)-carboxamide)

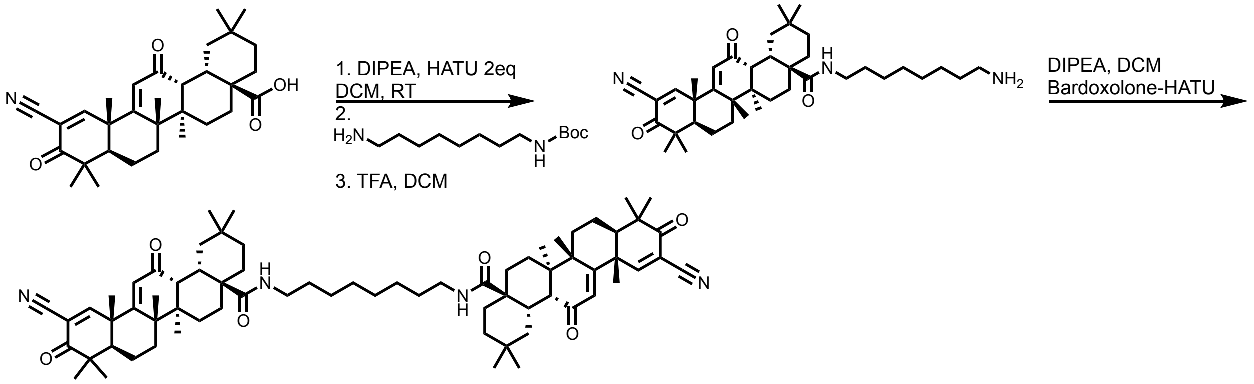


Synthetic procedure performed similar to the one described for EL133, using *tert-butyl (8-aminooctyl)carbamate* as the linker in the second step.

HR-MS (m/z): Calculated: 1090.75; Found: 1113.7347 [M+Na]+

**EL174**- (4a*S*,4a'*S*,6a*R*,6b*S*,6a'*R*,6b'*S*,8a*R*,8a'*R*,12a*S*,12a'*S*,14a*R*,14b*S*,14a'*R*,14b'*S*)-*N*a,*N*a'-(piperazine-1,4-diylbis(2-oxoethane-2,1-diyl))bis(11-cyano-2,2,6a,6b,9,9,12a-heptamethyl-10,14-dioxo-1,3,4,5,6,6a,6b,7,8,8a,9,10,12a,14,14a,14b-hexadecahydropicene-4a(2*H*)-carboxamide)

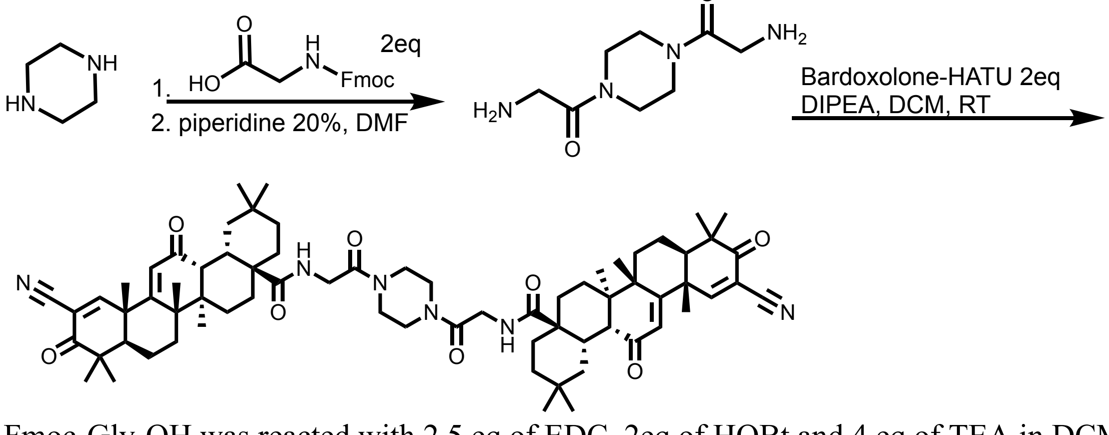


Fmoc-Gly-OH was reacted with 2.5 eq of EDC, 2eq of HOBt and 4 eq of TEA in DCM for 15 min on ice. Next, 30 mg of piperazine was added to the reaction which was allowed to warm to room temperature and kept overnight. The solvent was evaporated, and the crude product was dissolved in 20% piperidine in DMF, to allow Fmoc deprotection for 2 h in room temperature. The crude deprotected product was directly used for the next reaction, where it was reacted with HATU-Bardoxolone as described in the procedure for compound EL229.

HR-MS (m/z): Calculated: 1046.71; Found: 1169.7006 [M+Na]+

**EL221**- 3,3'-(((((((2,46-dioxo-6,9,12,15,18,21,24,27,30,33,36,39,42-tridecaoxa-3,45-diazaheptatetracontane-1,47-diyl)bis(oxy))bis(8-methoxy-1-methyl-2-oxo-1,2-dihydroquinoline-3,6-diyl))bis(azanediyl))bis(4-chloro-3,1-phenylene))bis(azanediyl))bis(methylene))dibenzenesulfonyl fluoride

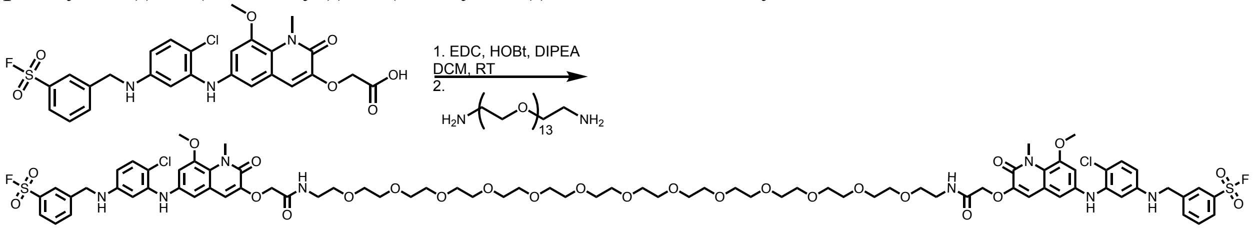


w8002 free acid was reacted with 2 eq EDC and 2 eq HOBt, along with 1.1 eq DIPEA in DCM for 15 min in room temperature. Next, 3,6,9,12,15,18,21,24,27,30,33,36,39-tridecaoxahentetracontane-1,41-diamine was dissolved in 100μL DMF, then added to the reaction mixture and allowed to react for 3 h in room temperature. The crude reaction was evaporated, dissolved in 40% ACN in $H_2O$, and purified by HPLC.

$^1$H NMR (500 MHz, DMSO) δ 8.45 (s, 1H), 8.03 – 7.95 (m, 1H), 7.86 (d, $J$ = 7.9 Hz, 1H), 7.74 – 7.69 (m, 1H), 7.41 (s, 1H), 7.10 – 7.06 (m, 2H), 7.05 (s, 1H), 6.87 (s, 1H), 6.75 (s, 1H), 6.53 (d, $J$ = 10.4 Hz, 2H), 6.18 (d, $J$ = 6.7 Hz, 2H), 4.54 (s, 3H), 4.38 (d, $J$ = 5.5 Hz, 2H), 3.83 (s, 3H), 3.78 (s, 3H), 3.48 (s, 28H).
$^{13}$C NMR (126 MHz, DMSO) δ 167.62, 158.06, 158.06, 148.93, 148.93, 148.35, 146.86, 146.86, 143.87, 141.00, 139.10, 135.58, 135.58, 130.92, 130.49, 127.14, 126.83, 122.77, 121.37, 114.42, 110.70, 108.60, 106.67, 105.29, 102.76, 70.22, 70.03, 69.30, 56.94, 46.16, 38.78, 35.44.
HR-MS (m/z): Calculated: 1746.57; Found: 1747.5796 [M+H]+, 1769.5701 [M+Na]+

**w8001**- 3-(((4-chloro-3-((8-methoxy-1-methyl-3-(2-(methylamino)-2-oxoethoxy)-2-oxo-1,2-dihydroquinolin-6-yl)amino)phenyl)amino)methyl)benzenesulfonyl fluoride

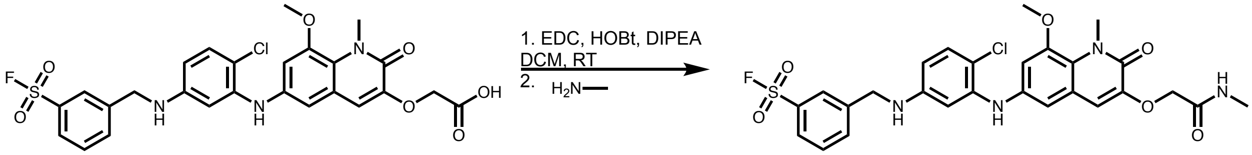


w8002 free acid was reacted with 2 eq EDC and 2 eq HOBt, along with 1.1 eq DIPEA in DCM for 15 min in room temperature. Next, 1 eq of methylamine was added to the reaction mixture and allowed to react for 3 h in room temperature. The crude reaction was evaporated, dissolved in 30% ACN in $H_2O$, and purified by HPLC.

HR-MS (m/z): Calculated: 588.12; Found: 611.1154 [M+Na]+

**EL217**- 3,3'-(((((((2,13-dioxo-6,9-dioxa-3,12-diazatetradecane-1,14-diyl)bis(oxy))bis(8-methoxy-1-methyl-2-oxo-1,2-dihydroquinoline-3,6-diyl))bis(azanediyl))bis(4-chloro-3,1-phenylene))bis(azanediyl))bis(methylene))dibenzenesulfonyl fluoride

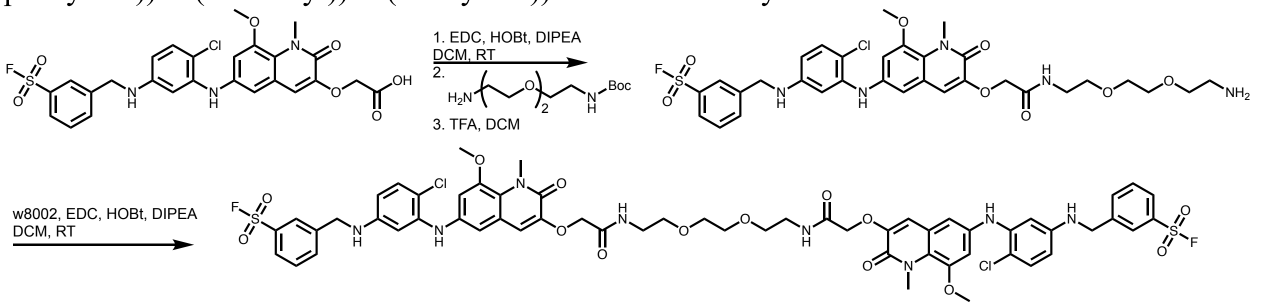


w8002 free acid was reacted with 2 eq EDC and 2 eq HOBt, along with 1.1 eq DIPEA in DCM for 15 min in room temperature. Next, *tert*-butyl (2-(2-(2-aminoethoxy)ethoxy)ethyl)carbamate was added to the reaction mixture and allowed to react for 3 h in room temperature. TFA was then added to double the solvent volume and left to deprotect overnight. The deprotected intermediate was then evaporated and purified by HPLC, lyophilized and was then used directly in the following reaction. W8002 was reacted with EDC, HOBt and DIPEA as described in the first step. The intermediate amine was dissolved in 100 µL DMF and added to the reaction mixture. After 2 h, the crude reaction was evaporated, dissolved in 40% ACN in $H_2O$, and purified by HPLC.

HR-MS (m/z): Calculated: 1262.29; Found: 1285.2777 [M+Na]+

**EL218**- 3,3'-(((((((2,19-dioxo-6,9,12,15-tetraoxa-3,18-diazaicosane-1,20-diyl)bis(oxy))bis(8-methoxy-1-methyl-2-oxo-1,2-dihydroquinoline-3,6-diyl))bis(azanediyl))bis(4-chloro-3,1-phenylene))bis(azanediyl))bis(methylene))dibenzenesulfonyl fluoride

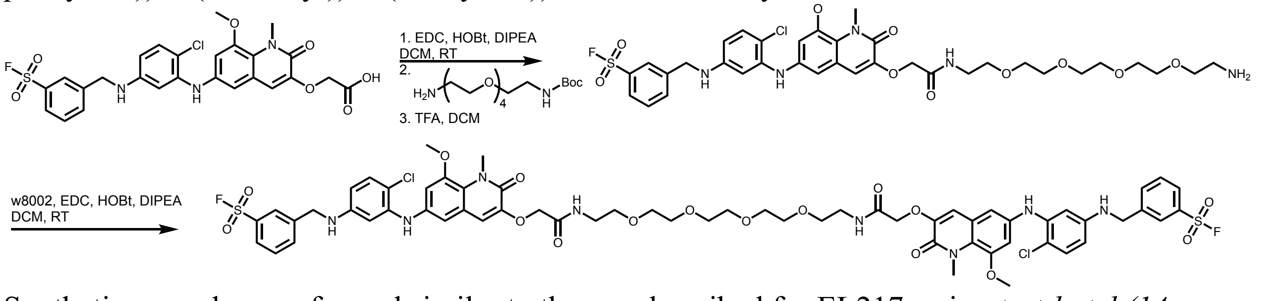


Synthetic procedure performed similar to the one described for EL217, using *tert-butyl (14-amino-3,6,9,12-tetraoxatetradecyl)carbamate* as the linker in the second step.

HR-MS (m/z): Calculated: 1350.34; Found: 1373.3314 [M+Na]+

**EL224**- 3,3'-(((((((2,37-dioxo-6,9,12,15,18,21,24,27,30,33-decaoxa-3,36-diazaoctatriacontane-1,38-diyl)bis(oxy))bis(8-methoxy-1-methyl-2-oxo-1,2-dihydroquinoline-3,6-diyl))bis(azanediyl))bis(4-chloro-3,1-phenylene))bis(azanediyl))bis(methylene))dibenzenesulfonyl fluoride

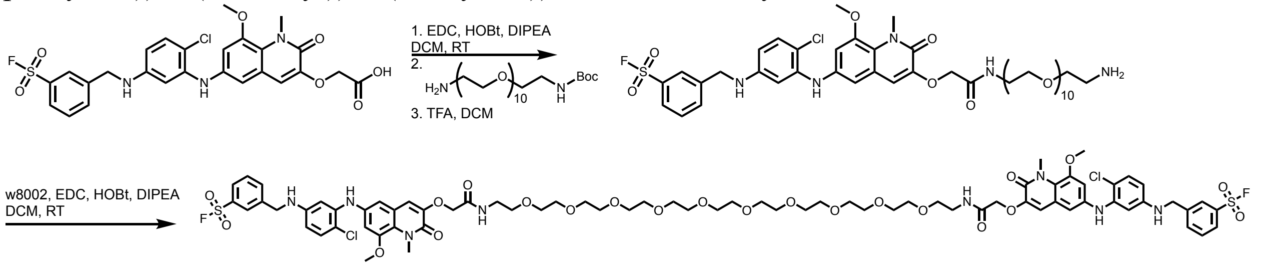


Synthetic procedure performed similar to the one described for EL217, using *tert-butyl (32-amino-3,6,9,12,15,18,21,24,27,30-decaoxadotriacontyl)carbamate* as the linker in the second step.

HR-MS (m/z): Calculated: 1614.50; Found: 1637.4777 [M+Na]+

**EL226**- 3,3'-(((((((2,34-dioxo-6,9,12,15,18,21,24,27,30-nonaoxa-3,33-diazapentatriacontane-1,35-diyl)bis(oxy))bis(8-methoxy-1-methyl-2-oxo-1,2-dihydroquinoline-3,6-diyl))bis(azanediyl))bis(4-chloro-3,1-phenylene))bis(azanediyl))bis(methylene))dibenzenesulfonyl fluoride

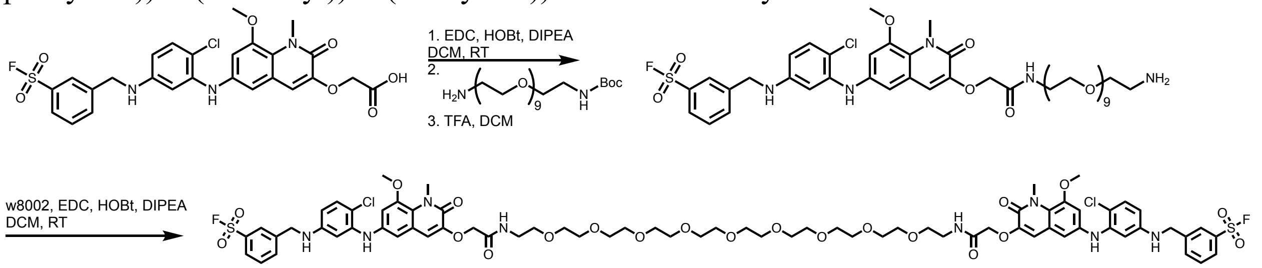


Synthetic procedure performed similar to the one described for EL217, using *tert-butyl (29-amino-3,6,9,12,15,18,21,24,27-nonaoxanonacosyl)carbamate* as the linker in the second step.

HR-MS (m/z): Calculated: 1570.47; Found: 1593.4633 [M+Na]+

**EL227**- 3,3'-(((((((2,31-dioxo-6,9,12,15,18,21,24,27-octaoxa-3,30-diazadotriacontane-1,32-diyl)bis(oxy))bis(8-methoxy-1-methyl-2-oxo-1,2-dihydroquinoline-3,6-diyl))bis(azanediyl))bis(4-chloro-3,1-phenylene))bis(azanediyl))bis(methylene))dibenzenesulfonyl fluoride

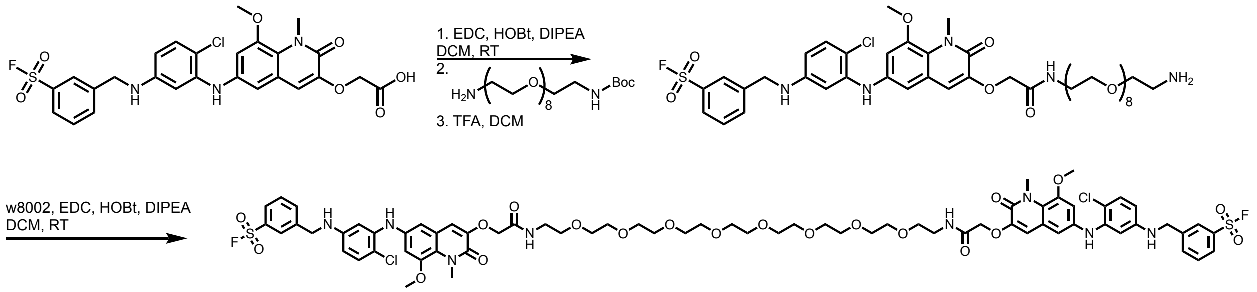


Synthetic procedure performed similar to the one described for EL217, using *tert-butyl (26-amino-3,6,9,12,15,18,21,24-octaoxahexacosyl)carbamate* as the linker in the second step.

HR-MS (m/z): Calculated: 1526.44; Found: 1549.4396 [M+Na]+

**EL225**- 3,3'-(((((((2,40-dioxo-6,9,12,15,18,21,24,27,30,33,36-undecaoxa-3,39-diazahentetracontane-1,41-diyl)bis(oxy))bis(8-methoxy-1-methyl-2-oxo-1,2-dihydroquinoline-3,6-diyl))bis(azanediyl))bis(4-chloro-3,1-phenylene))bis(azanediyl))bis(methylene))dibenzenesulfonyl fluoride

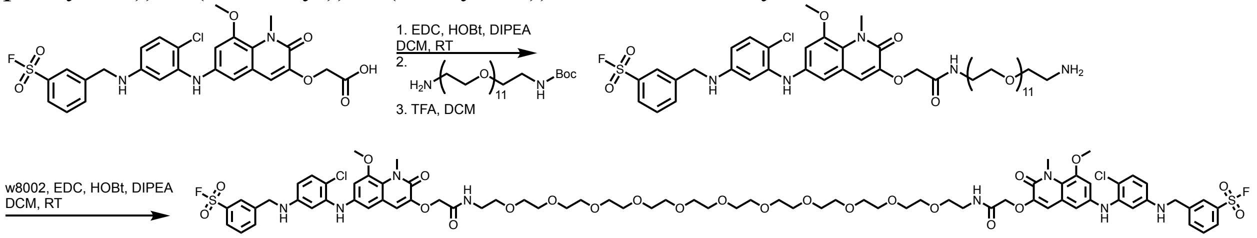


Synthetic procedure performed similar to the one described for EL217, using *tert-butyl (35-amino-3,6,9,12,15,18,21,24,27,30,33-undecaoxapentatriacontyl)carbamate* as the linker in the second step.

$^{1}$H NMR (500 MHz, DMSO) δ 8.47 (s, 1H), 8.01 (s, 1H), 7.99 – 7.96 (m, 1H), 7.86 (d, *J* = 7.7 Hz, 1H), 7.72 (t, *J* = 7.8 Hz, 1H), 7.41 (s, 1H), 7.08 (d, *J* = 8.7 Hz, 2H), 7.05 (s, 1H), 6.87 (d, *J* = 1.9 Hz, 1H), 6.75 (d, *J* = 1.9 Hz, 1H), 6.56 – 6.53 (m, 1H), 6.52 (d, *J* = 2.4 Hz, 1H), 6.18 (dd, *J* = 8.7, 2.3 Hz, 1H), 4.54 (s, 2H), 4.38 (d, *J* = 5.8 Hz, 2H), 3.82 (d, *J* = 5.4 Hz, 2H), 3.78 (s, 3H), 3.48 (s, 22H).
$^{13}$C NMR (126 MHz, DMSO) δ 167.63, 158.06, 148.94, 148.40, 146.86, 143.88, 141.00, 139.11, 135.58, 130.92, 130.52, 127.14, 126.83, 122.77, 121.36, 114.42, 110.70, 108.59, 106.67, 105.29, 102.77, 70.21, 70.15, 70.03, 69.30, 68.24, 56.94, 46.16, 38.78, 35.44.
HR-MS (m/z): Calculated: 1658.52; Found: 1681.5098 [M+Na]+

**EL228**- 3,3'-(((((((2,43-dioxo-6,9,12,15,18,21,24,27,30,33,36,39-dodecaoxa-3,42-diazatetratetracontane-1,44-diyl)bis(oxy))bis(8-methoxy-1-methyl-2-oxo-1,2-dihydroquinoline-3,6-diyl))bis(azanediyl))bis(4-chloro-3,1-

phenylene))bis(azanediyl))bis(methylene))dibenzenesulfonyl fluoride

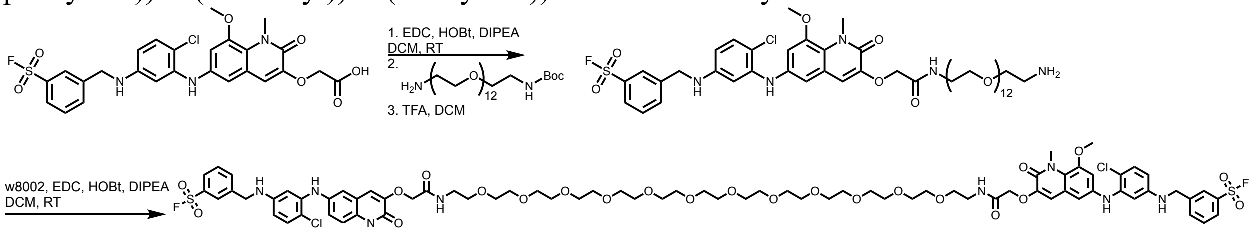


Synthetic procedure performed similar to the one described for EL217, using *tert-butyl (38-amino-3,6,9,12,15,18,21,24,27,30,33,36-dodecaoxaoctatriacontyl)carbamate* as the linker in the second step.

HR-MS (m/z): Calculated: 1702.55; Found: 1725.5461 [M+Na]+

**EL230**- 3-(((4-chloro-3-((3-((1-((4a*S*,6a*R*,6b*S*,8a*R*,12a*S*,14a*R*,14b*S*)-11-cyano-2,2,6a,6b,9,9,12a-heptamethyl-10,14-dioxo-1,3,4,5,6,6a,6b,7,8,8a,9,10,12a,14,14a,14b-hexadecahydropicen-4a(2*H*)-yl)-1,42-dioxo-5,8,11,14,17,20,23,26,29,32,35,38-dodecaoxa-2,41-diazatritetracontan-43-yl)oxy)-8-methoxy-1-methyl-2-oxo-1,2-dihydroquinolin-6-yl)amino)phenyl)amino)methyl)benzenesulfonyl fluoride

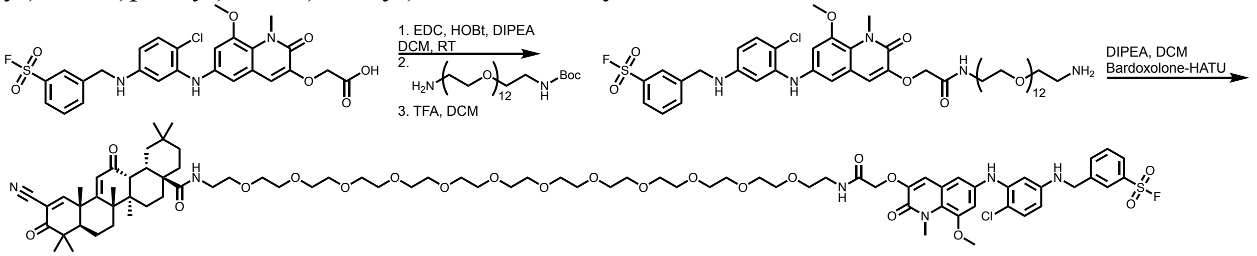


Synthetic procedure performed similar to the one described for EL217, using *tert-butyl (38-amino-3,6,9,12,15,18,21,24,27,30,33,36-dodecaoxaoctatriacontyl)carbamate* as the linker in the second step. The intermediate deprotected amine was purified by HPLC, lyophilized and was then used directly in the following reaction. Bardoxlone-HATU, prepared as described for compound EL229 was added to the reaction along with 3 eq of DIPEA in DCM. After 2 h in room temperature, the crude reaction was evaporated, dissolved in 50% ACN in $H_2O$, and purified by HPLC.

$^{1}$H NMR (500 MHz, DMSO) δ 8.65 (s, 1H), 8.01 (d, *J* = 7.7 Hz, 1H), 7.98 (d, *J* = 6.8 Hz, 1H), 7.86 (d, *J* = 7.7 Hz, 1H), 7.73 (dd, *J* = 14.8, 6.9 Hz, 2H), 7.41 (s, 1H), 7.09 (d, *J* = 8.7 Hz, 1H), 7.05 (s, 1H), 6.87 (d, *J* = 2.0 Hz, 1H), 6.75 (d, *J* = 1.8 Hz, 1H), 6.55 (d, *J* = 5.9 Hz, 1H), 6.54 – 6.49 (m, 2H), 6.23 – 6.15 (m, 2H), 4.54 (s, 2H), 4.38 (d, *J* = 5.8 Hz, 2H), 3.83 (s, 3H), 3.79 (s, 3H), 3.49 (dd, *J* = 5.7, 3.0 Hz, 48H), 3.47 – 3.39 (m, 6H), 3.30 (t, *J* = 5.6 Hz, 4H), 1.92 – 1.75 (m, 3H), 1.65 (dd, *J* = 27.7, 15.1 Hz, 5H), 1.55 (dd, *J* = 13.6, 3.9 Hz, 2H), 1.43 (s, 3H), 1.24 (s, 3H), 1.17 (s, 3H), 1.06 (s, 3H), 0.93 (s, 3H), 0.89 (s, 3H), 0.85 (s, 3H).
$^{13}$C NMR (126 MHz, DMSO) δ 199.73, 197.79, 177.10, 168.82, 167.62, 158.06, 148.94, 148.40, 146.87, 143.88, 141.00, 139.11, 135.58, 132.16, 130.93, 130.52, 127.14, 126.83, 123.83, 122.77, 121.37, 114.43, 113.27, 110.71, 108.58, 106.68, 105.29, 102.77, 70.23, 70.16, 70.04, 69.99, 69.46, 69.30, 68.25, 56.94, 49.13, 46.68, 46.16, 45.86, 45.80, 44.83, 43.01, 41.98, 39.23, 38.78,

35.80, 35.45, 34.62, 33.72, 33.46, 31.22, 30.69, 27.68, 26.57, 26.32, 24.70, 23.60, 22.23, 21.76, 21.59, 17.94.
HR-MS (m/z): Calculated: 1618.76; Found: 1641.7552 [M+Na]+

**OS-47B**- 2,2'-(((6,25-bis(2-((2-methylbenzo[*d*]thiazol-6-yl)amino)-2-oxoethyl)-4,7,24,27-tetraoxo-11,14,17,20-tetraoxa-5,8,23,26-tetraazatriacontane-1,30-diyl)bis(4,1-phenylene))bis(methylene))dimalonic acid

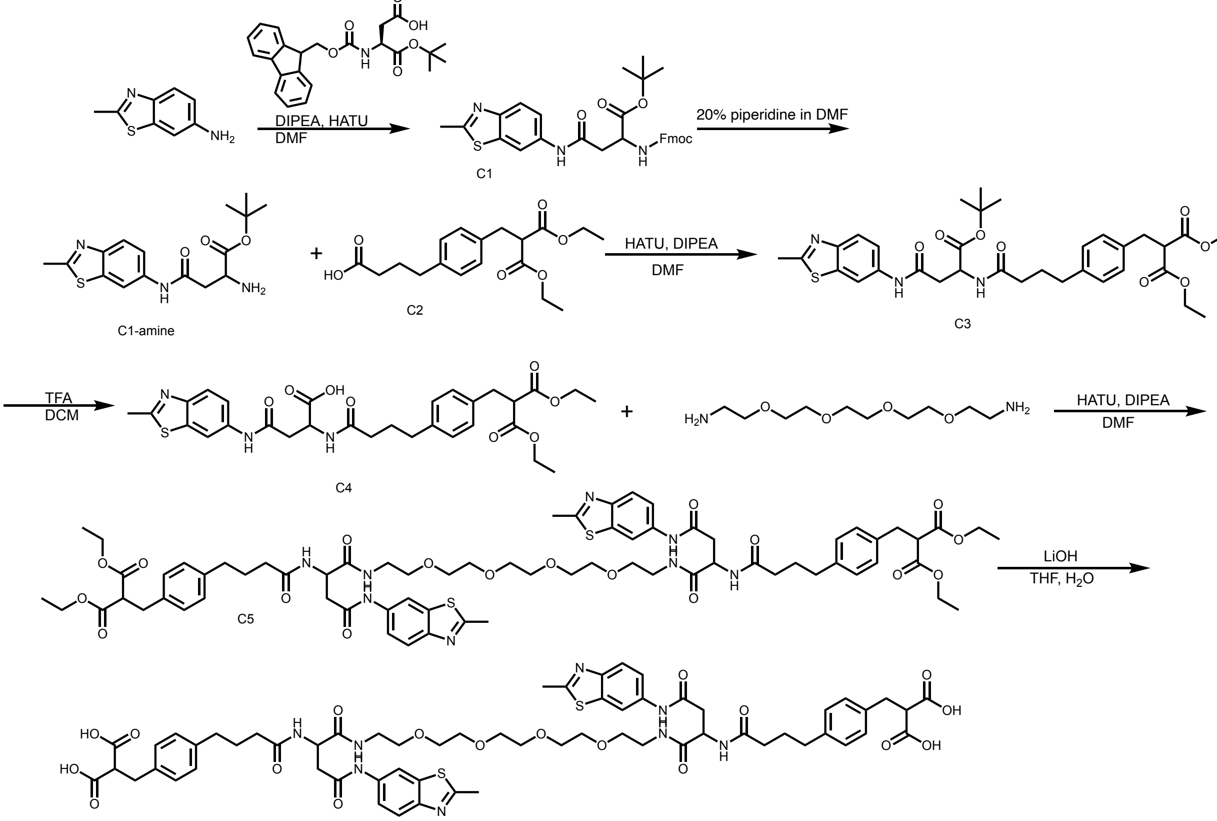


2-methylbenzo[d]thiazol-6-amine was coupled to Fmoc-L-aspartic acid-α-tert-butylester in the presence of DIPEA and HATU, in DMF for overnight at room temperature. The mixture was evaporated to dryness and the product was extracted using EtOAc, followed by sequential washes with water and brine. Finally, the organic phase was dried over $Na_2SO_4$ and concentrated under reduced pressure, and the crude was purified using flash chromatography (DCM:MeOH). Next, the Fmoc protecting group was removed using 20% piperidine in DMF for 1 h at room temperature, and the product was purified using flash chromatography. The resulting amine was then reacted with compound C2 which was synthesized according to the previously reported procedure[2], using DIPEA and HATU, in DMF for overnight at room temperature. The solvent was evaporated to dryness, and the crude C3 was redissolved in DCM and washed with water, 0.5 M HCl, and brine. The organic phase was dried over $Na_2SO_4$, evaporated under reduced pressure, and purified using flash chromatography. Deprotection of Boc group was performed in 1:1 TFA/DCM to afford compound C4. The free acid was then coupled on both ends of the diamine linker 3,6,9,12-tetraoxatetradecane-1,14-diamine in a single step using DIPEA and HATU, in DMF for overnight at room temperature. Resulting compound C5 was purified by

HPLC, then reacted with 1 M LiOH in a 1:1 THF/H2O solution to afford the final compound, which was purified by HPLC.

$^{1}$H NMR (500 MHz, DMSO) δ 12.72 (s, 2H), 10.11 (s, 1H), 8.39 (d, J = 1.8 Hz, 1H), 8.08 (d, J = 8.0 Hz, 1H), 7.83 (t, J = 5.6 Hz, 1H), 7.80 (d, J = 8.8 Hz, 1H), 7.47 (dd, J = 8.8, 1.9 Hz, 1H), 7.03 (dd, J = 25.5, 8.0 Hz, 4H), 4.69 (dd, J = 14.0, 7.9 Hz, 1H), 3.52 (t, J = 7.8 Hz, 2H), 3.46 (d, J = 5.2 Hz, 6H), 3.38 (t, J = 6.1 Hz, 2H), 3.20 (qd, J = 13.5, 7.0 Hz, 2H), 2.97 (d, J = 7.7 Hz, 2H), 2.79 (dt, J = 10.0, 5.1 Hz, 1H), 2.75 (s, 3H), 2.62 (dd, J = 15.2, 8.1 Hz, 1H), 2.49 – 2.43 (m, 2H), 2.12 (t, J = 7.3 Hz, 2H), 1.73 (dt, J = 13.9, 7.0 Hz, 2H), 1.29 – 1.23 (m, 1H).
$^{13}$C NMR (126 MHz, DMSO) δ 172.42, 171.44, 170.71, 168.90, 165.84, 149.37, 140.20, 136.66, 136.23, 136.20, 129.00, 128.64, 122.27, 118.69, 111.69, 70.20, 70.14, 70.03, 69.31, 53.82, 50.12, 39.17, 39.14, 35.24, 34.58, 34.26, 27.41, 20.08.
HR-MS (m/z): Calculated: 1281.4460; Found: 1281.4456 [M+H]+

**NMR spectra**

EL133:

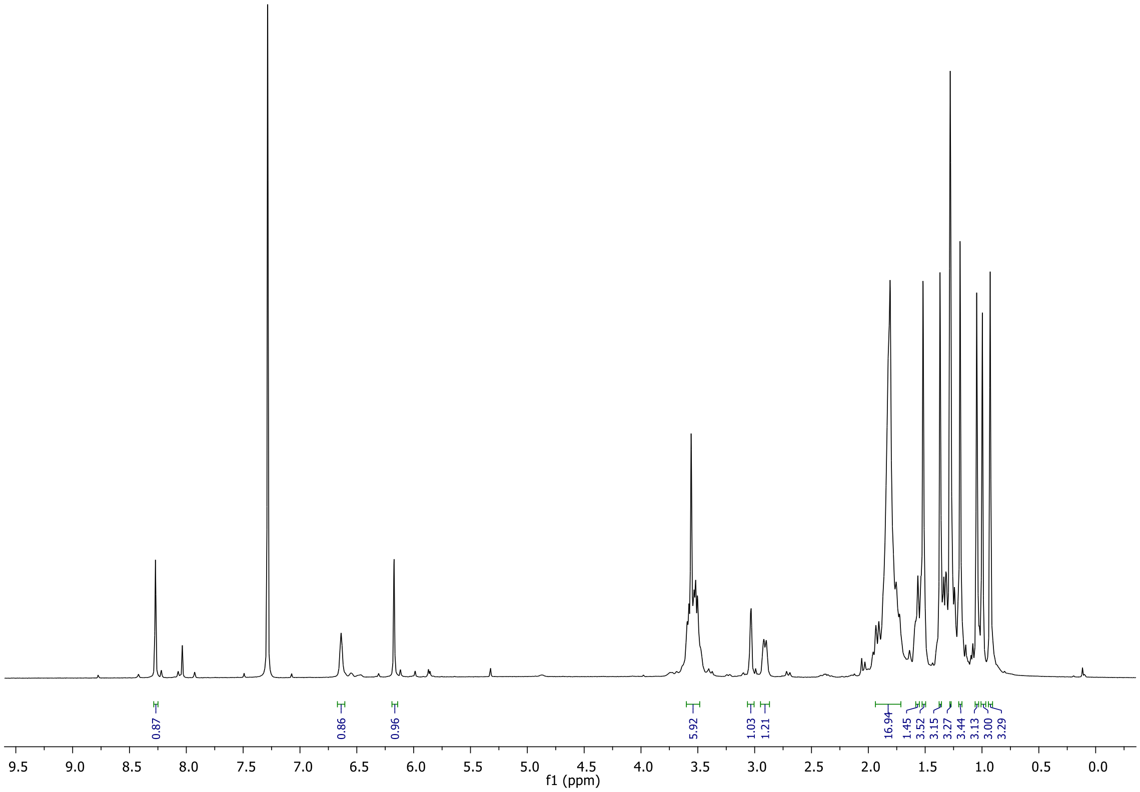


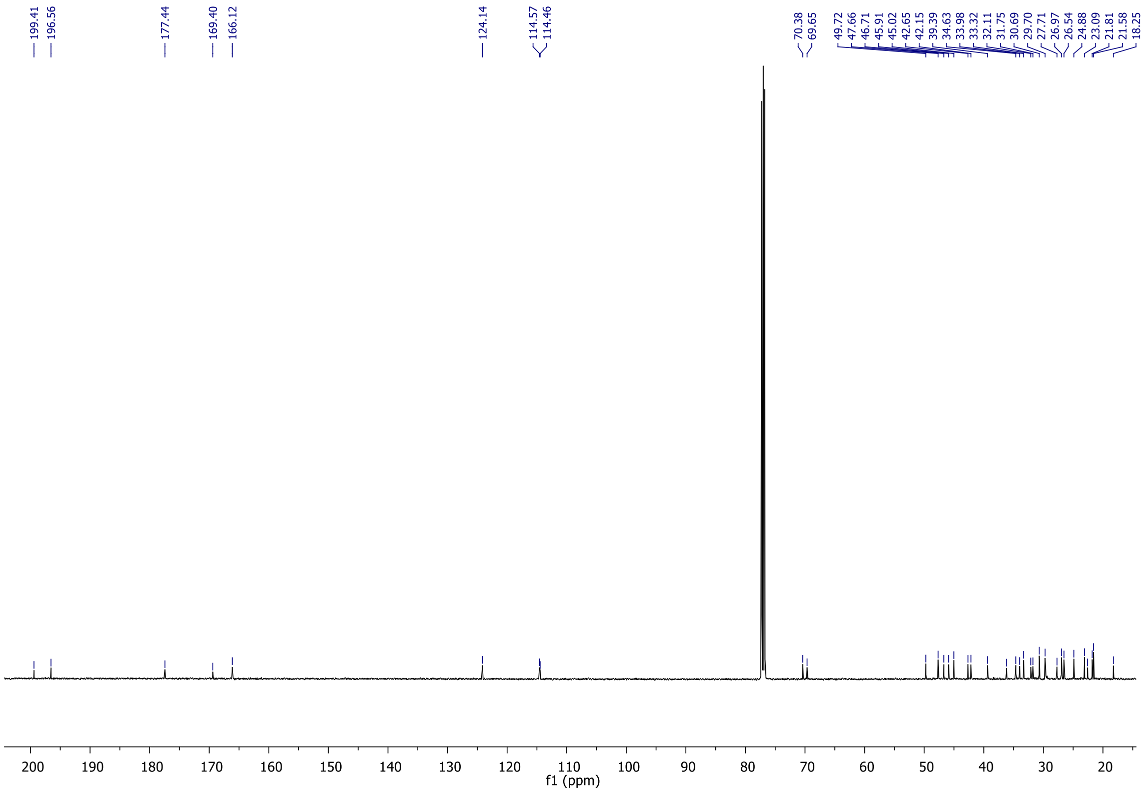

EL164:

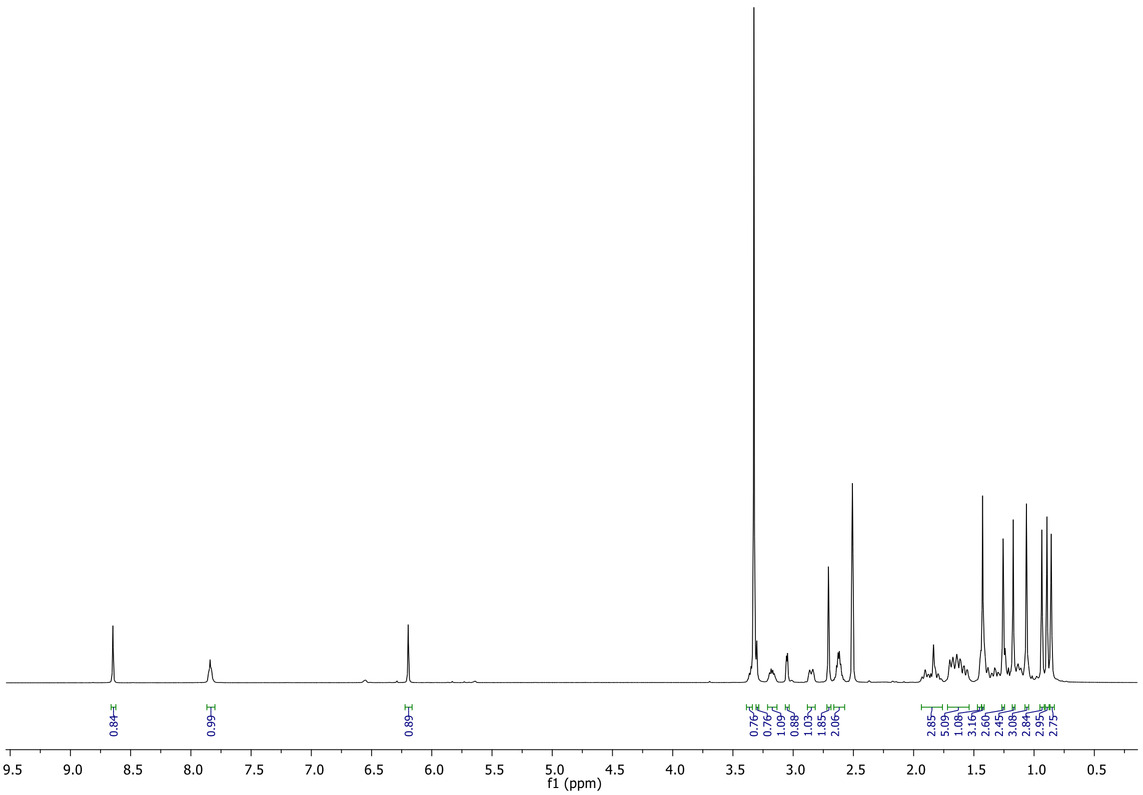
0.84
0.99
0.89
0.76
0.76
1.09
0.88
1.03
1.85
2.06
2.85
5.09
1.08
3.16
2.60
2.45
3.08
2.84
2.95
2.75
f1 (ppm)

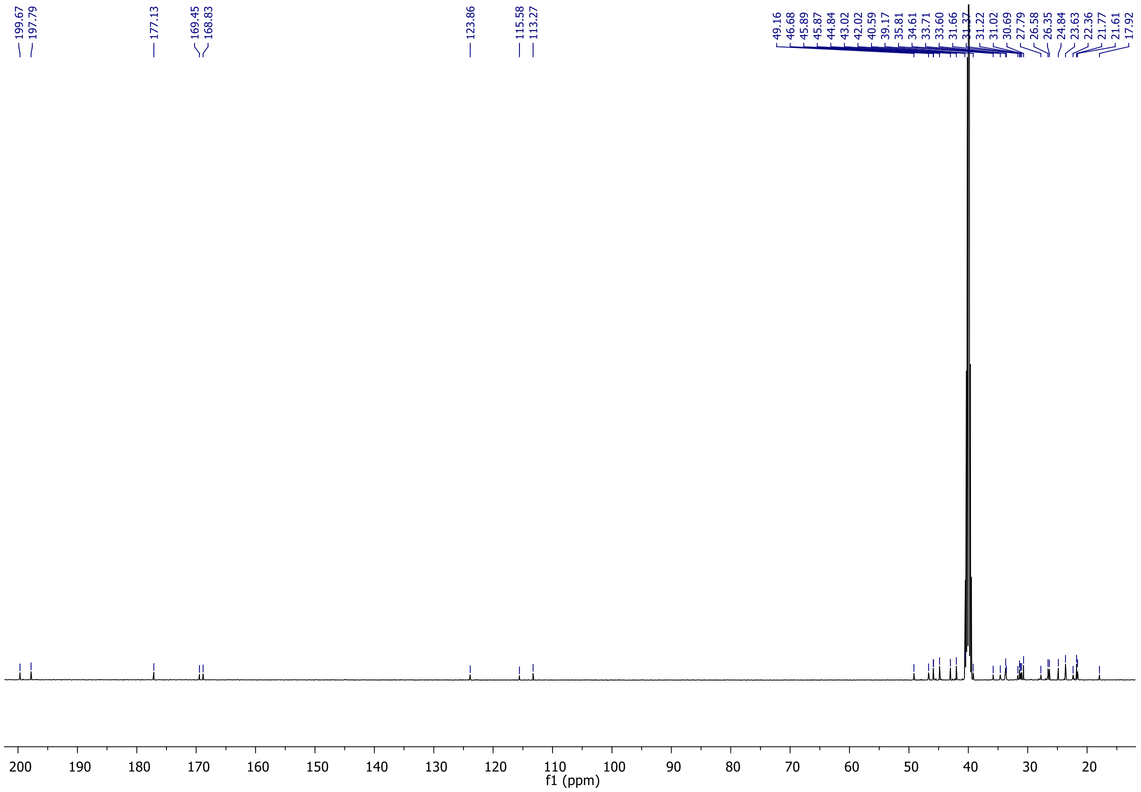
199.67
197.79
177.13
169.45
168.83
123.86
115.58
113.27
49.16
46.68
45.89
45.87
44.84
43.02
42.02
40.59
39.17
35.81
34.61
33.71
33.60
31.66
31.37
31.22
31.02
30.69
27.79
26.58
26.35
24.84
23.63
22.36
21.77
21.61
17.92
f1 (ppm)

EL165:

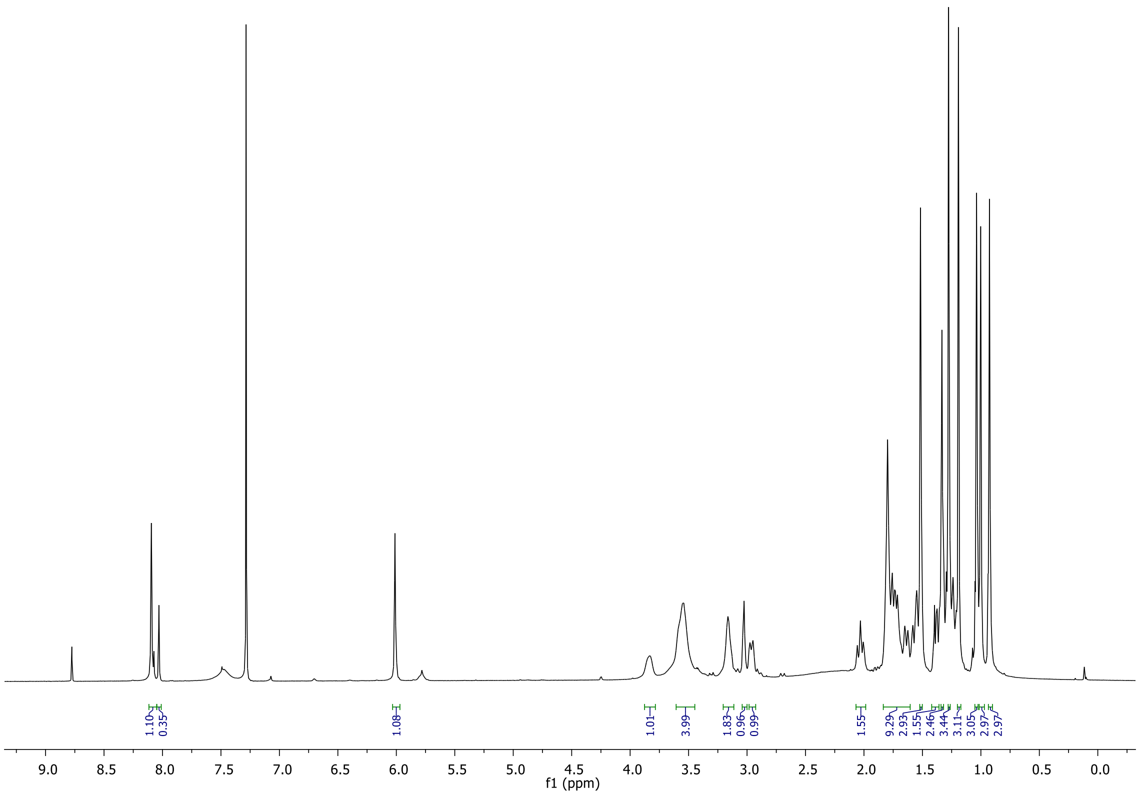

1.10
0.35
1.08
1.01
3.99
1.83
0.96
0.99
1.55
9.29
2.93
1.55
2.46
3.44
3.11
3.05
2.97
2.97
f1 (ppm)


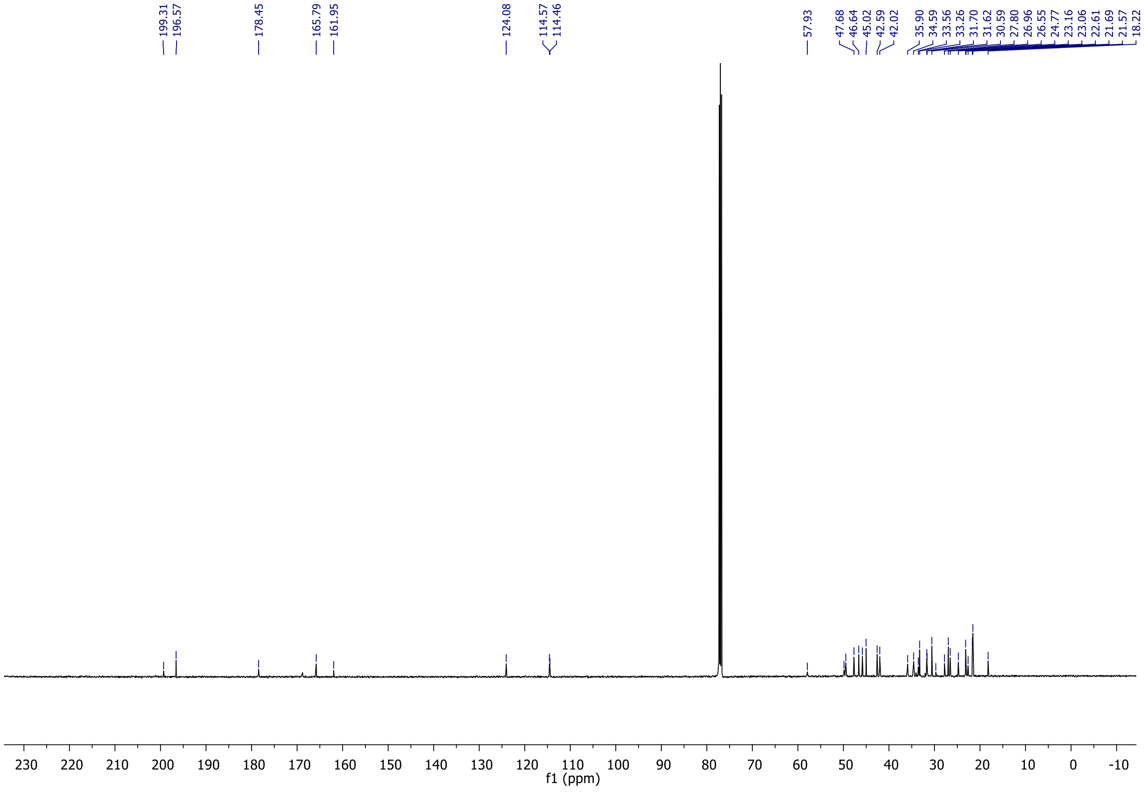

199.31
196.57
178.45
165.79
161.95
124.08
114.57
114.46
57.93
47.68
46.64
45.02
42.59
42.02
35.90
34.59
33.56
33.26
31.70
31.62
30.59
27.80
26.96
26.55
24.77
23.16
23.06
22.61
21.69
21.57
18.22
f1 (ppm)

EL229:

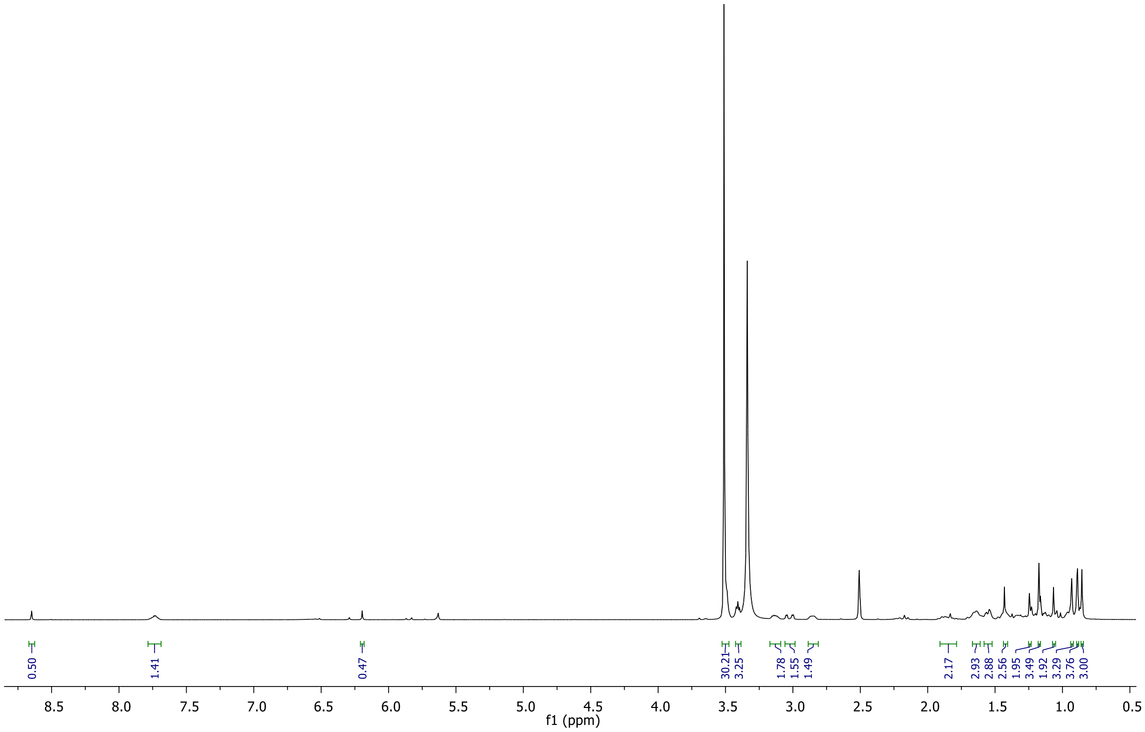

0.50
1.41
0.47
30.21
3.25
1.78
1.55
1.49
2.17
2.93
2.88
2.56
1.95
3.49
1.92
3.29
3.76
3.00
f1 (ppm)


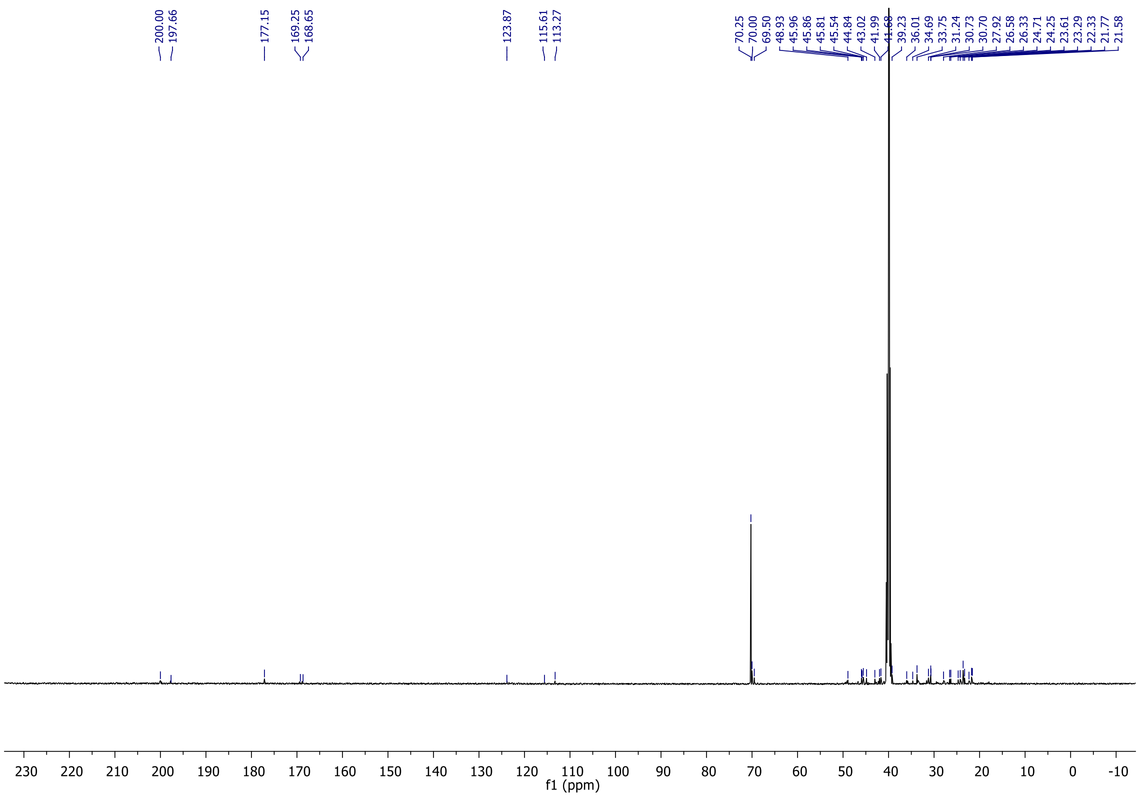

200.00
197.66
177.15
169.25
168.65
123.87
115.61
113.27
70.25
70.00
69.50
48.93
45.96
45.86
45.81
45.54
44.84
43.02
41.99
39.23
36.01
34.69
33.75
31.24
30.73
30.70
27.92
26.58
26.33
24.71
24.25
23.61
23.29
22.33
21.77
21.58
f1 (ppm)

EL228:

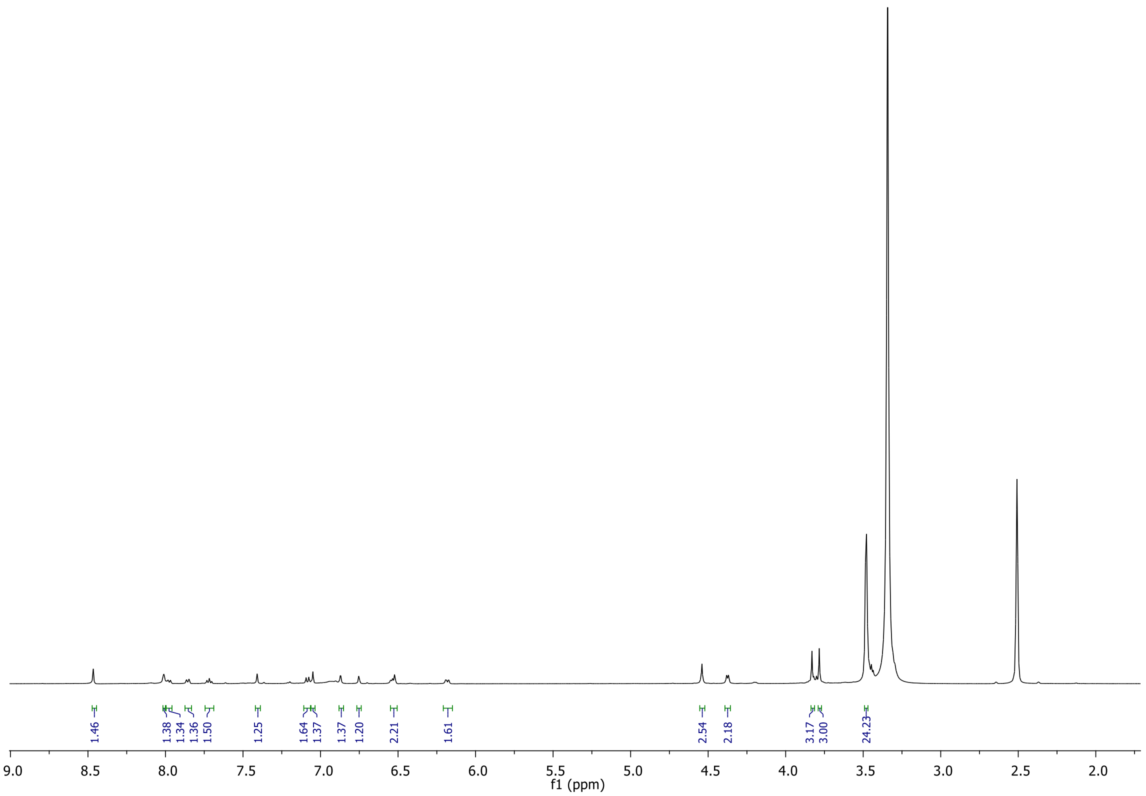
1.46
1.38
1.34
1.36
1.50
1.25
1.64
1.37
1.37
1.20
2.21
1.61
2.54
2.18
3.17
3.00
24.23
9.0
8.5
8.0
7.5
7.0
6.5
6.0
5.5
5.0
4.5
4.0
3.5
3.0
2.5
2.0
f1 (ppm)

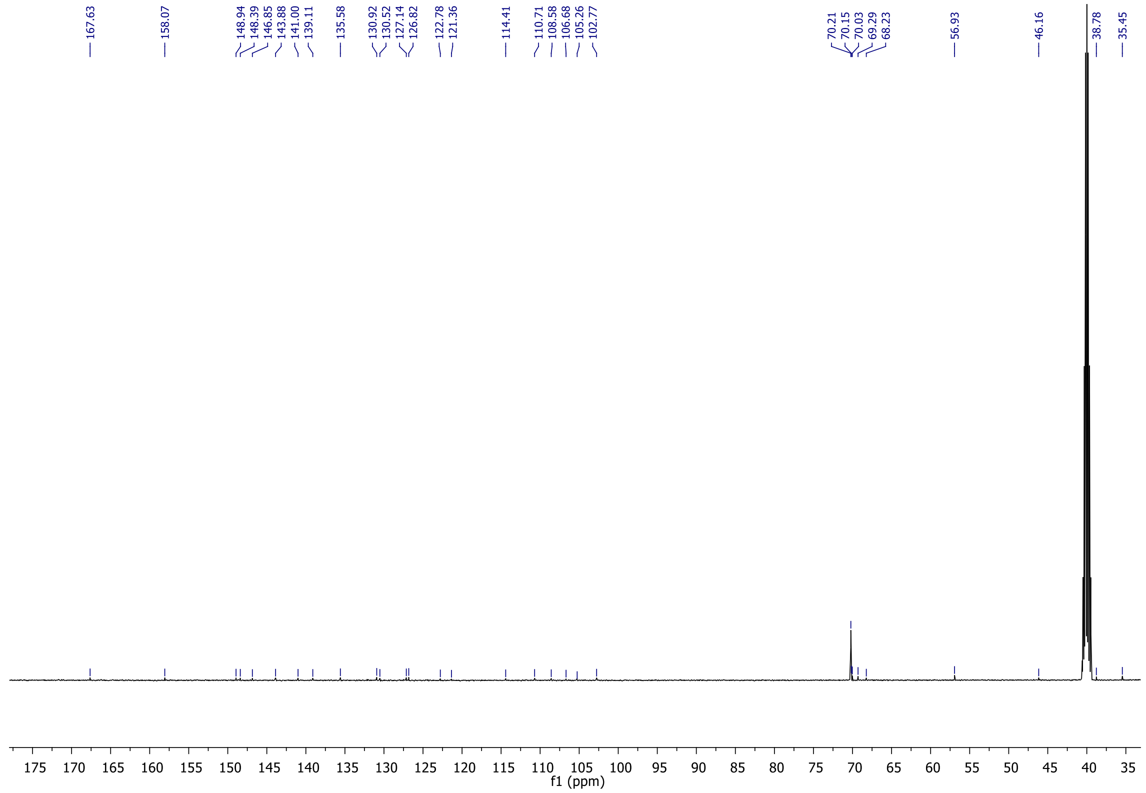
167.63
158.07
148.94
148.39
146.85
143.88
141.00
139.11
135.58
130.92
130.52
127.14
126.82
122.78
121.36
114.41
110.71
108.58
106.68
105.26
102.77
70.21
70.15
70.03
69.29
68.23
56.93
46.16
38.78
35.45
175
170
165
160
155
150
145
140
135
130
125
120
115
110
105
100
95
90
85
80
75
70
65
60
55
50
45
40
35
f1 (ppm)

EL221:

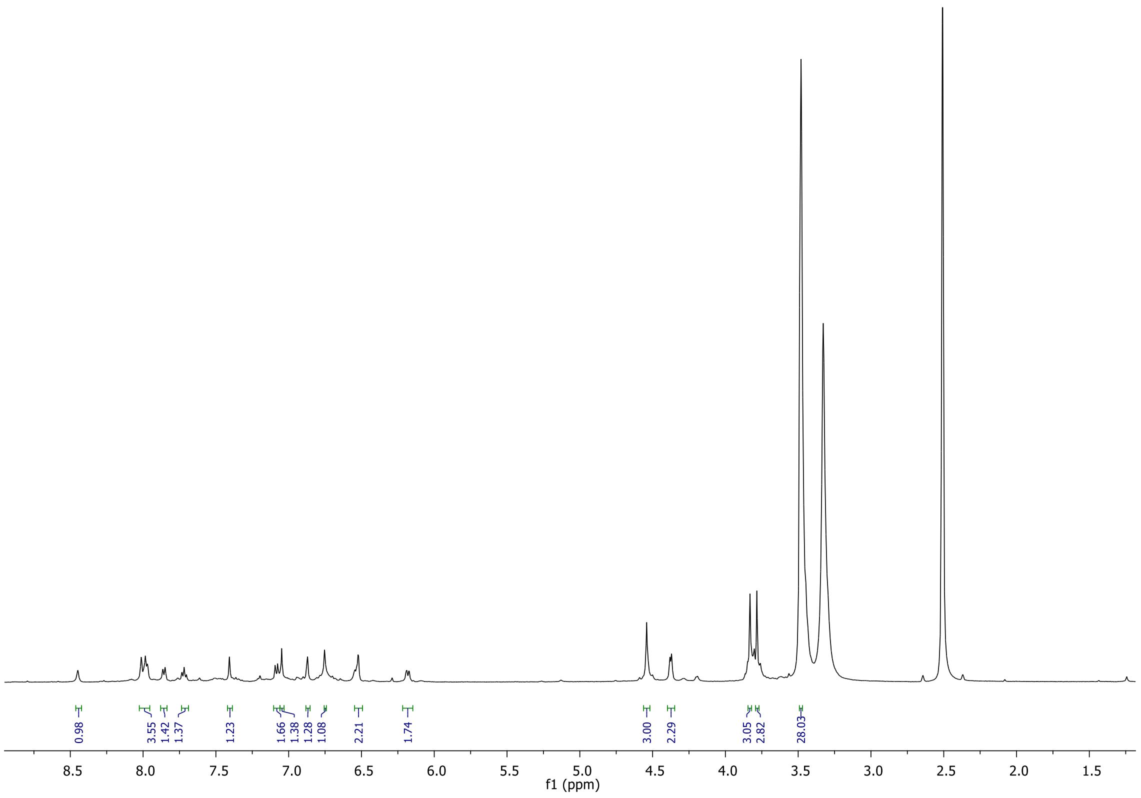
0.98
3.55
1.42
1.37
1.23
1.66
1.38
1.28
1.08
2.21
1.74
3.00
2.29
3.05
2.82
28.03
f1 (ppm)

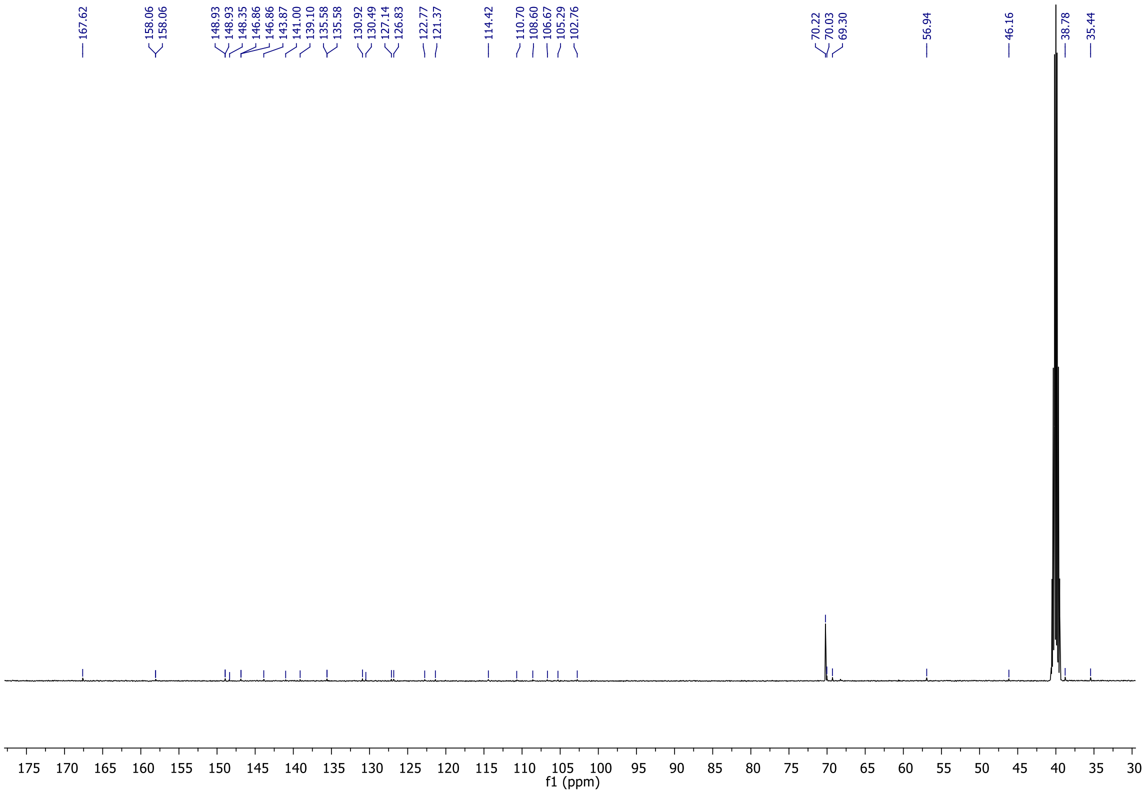
167.62
158.06
158.06
148.93
148.93
148.35
146.86
146.86
143.87
141.00
139.10
135.58
135.58
130.92
130.49
127.14
126.83
122.77
121.37
114.42
110.70
108.60
106.67
105.29
102.76
70.22
70.03
69.30
56.94
46.16
38.78
35.44
f1 (ppm)

EL225:

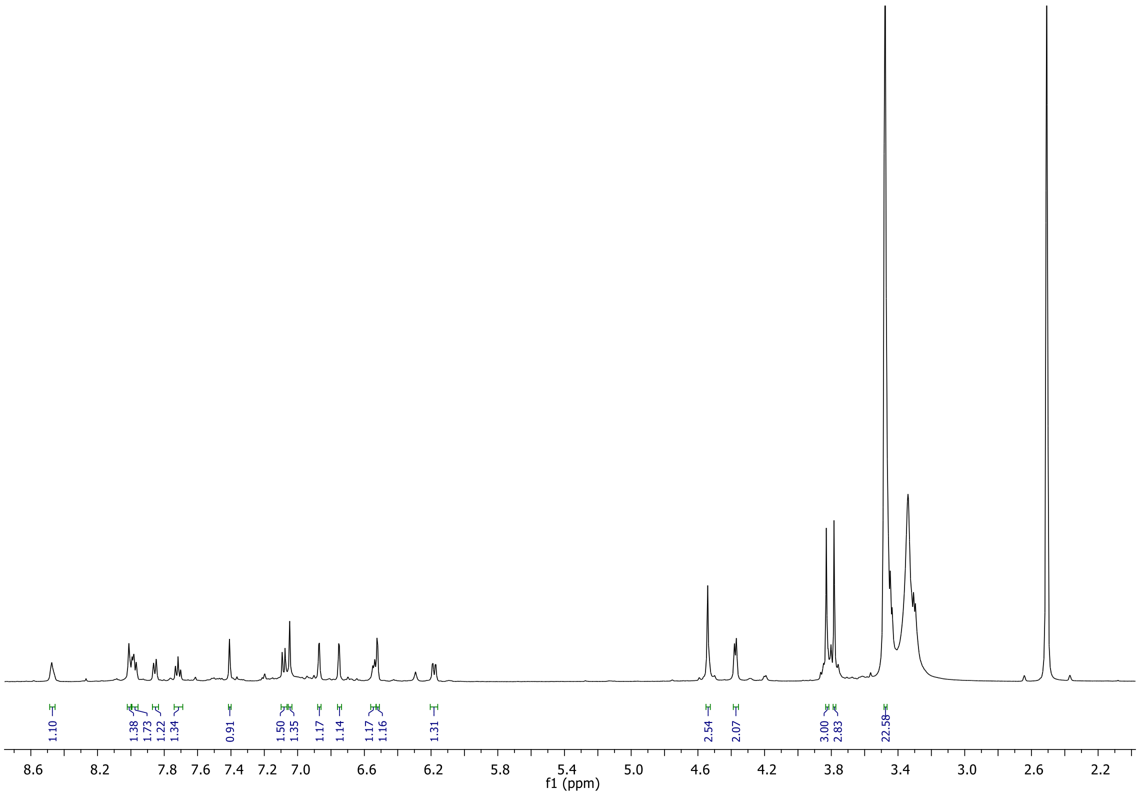
1.10
1.38
1.73
1.22
1.34
0.91
1.50
1.35
1.17
1.14
1.17
1.16
1.31
2.54
2.07
3.00
2.83
22.58
f1 (ppm)

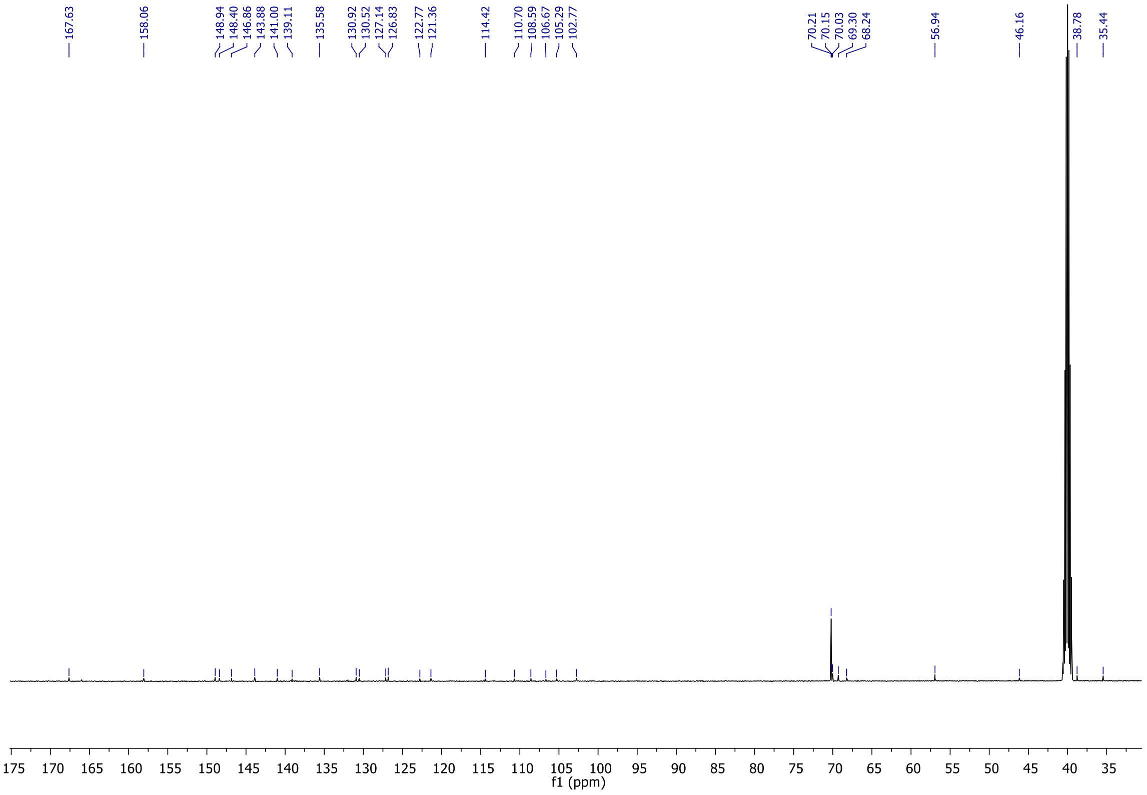
167.63
158.06
148.94
148.40
146.86
143.88
141.00
139.11
135.58
130.92
130.52
127.14
126.83
122.77
121.36
114.42
110.70
108.59
106.67
105.29
102.77
70.21
70.15
70.03
69.30
68.24
56.94
46.16
38.78
35.44
f1 (ppm)

OS-47B:

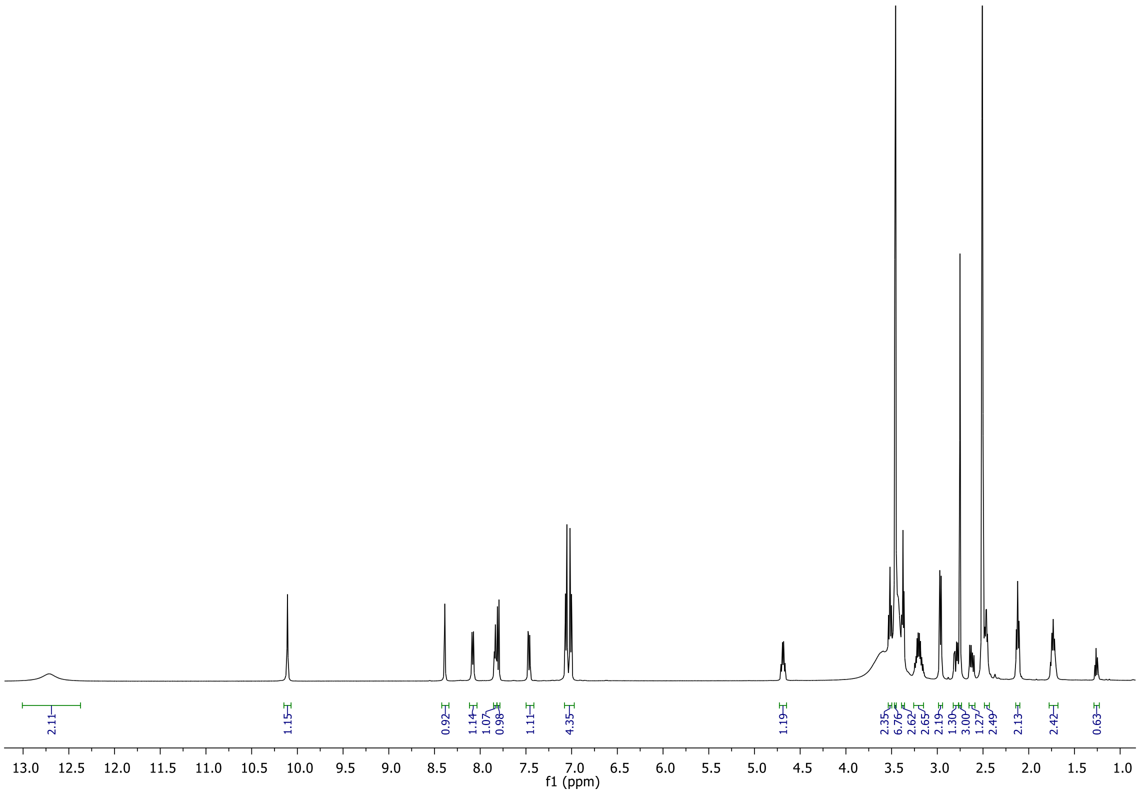

2.11
1.15
0.92
1.14
1.07
0.98
1.11
4.35
1.19
2.35
6.76
2.62
2.65
2.19
1.30
3.00
1.27
2.49
2.13
2.42
0.63
f1 (ppm)


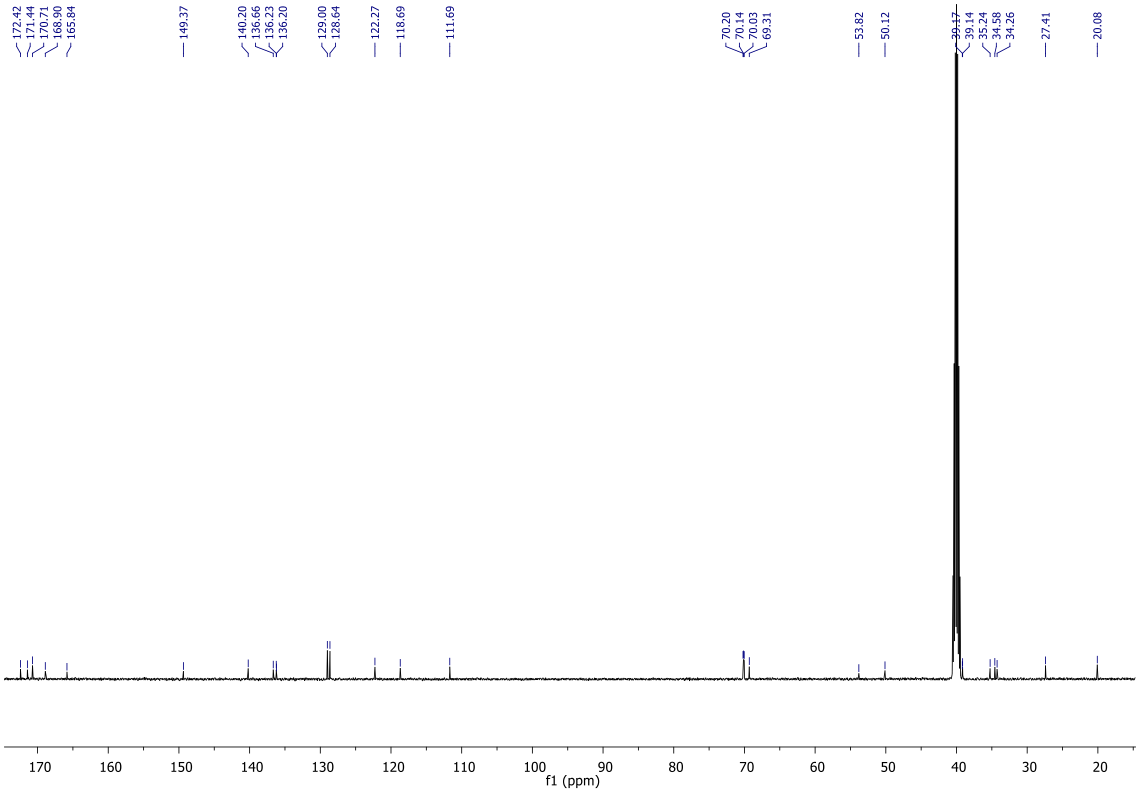

172.42
171.44
170.71
168.90
165.84
149.37
140.20
136.66
136.23
136.20
129.00
128.64
122.27
118.69
111.69
70.20
70.14
70.03
69.31
53.82
50.12
39.17
39.14
35.24
34.58
34.26
27.41
20.08
f1 (ppm)

EL132:

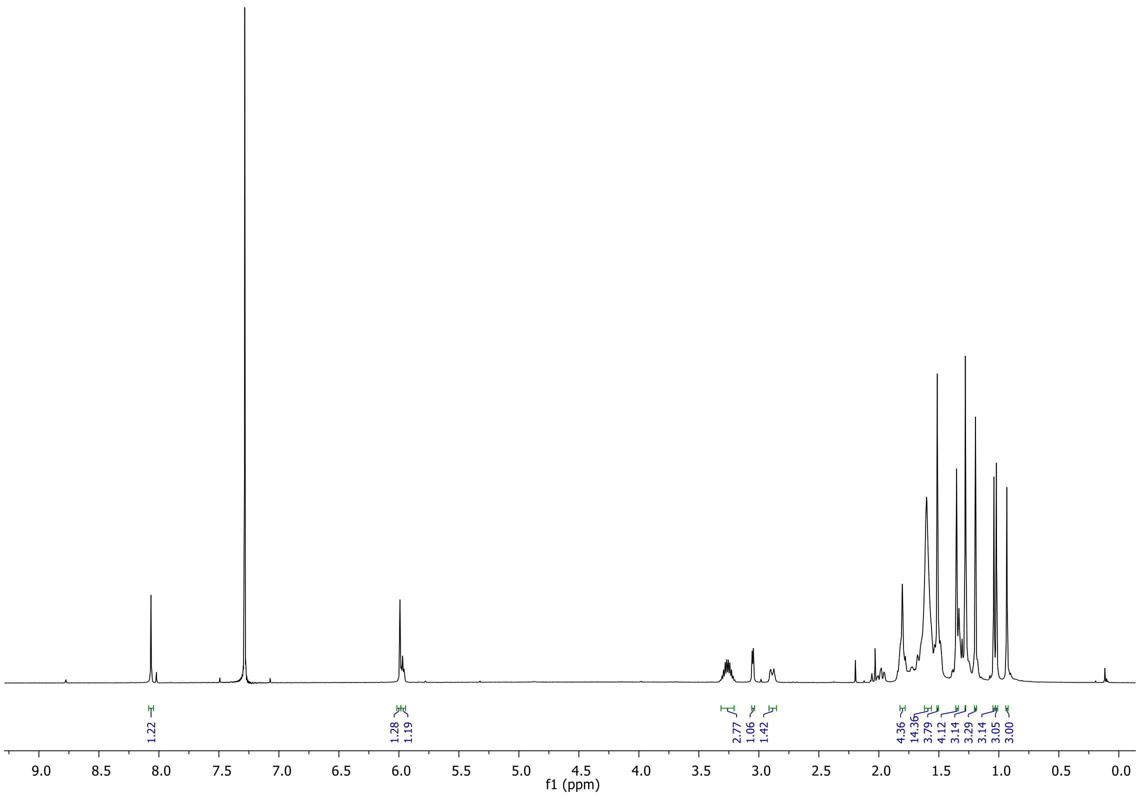
1.22
1.28
1.19
2.77
1.06
1.42
4.36
14.36
3.79
4.12
3.14
3.29
3.14
3.05
3.00
9.0
8.5
8.0
7.5
7.0
6.5
6.0
5.5
5.0
4.5
4.0
3.5
3.0
2.5
2.0
1.5
1.0
0.5
0.0
f1 (ppm)

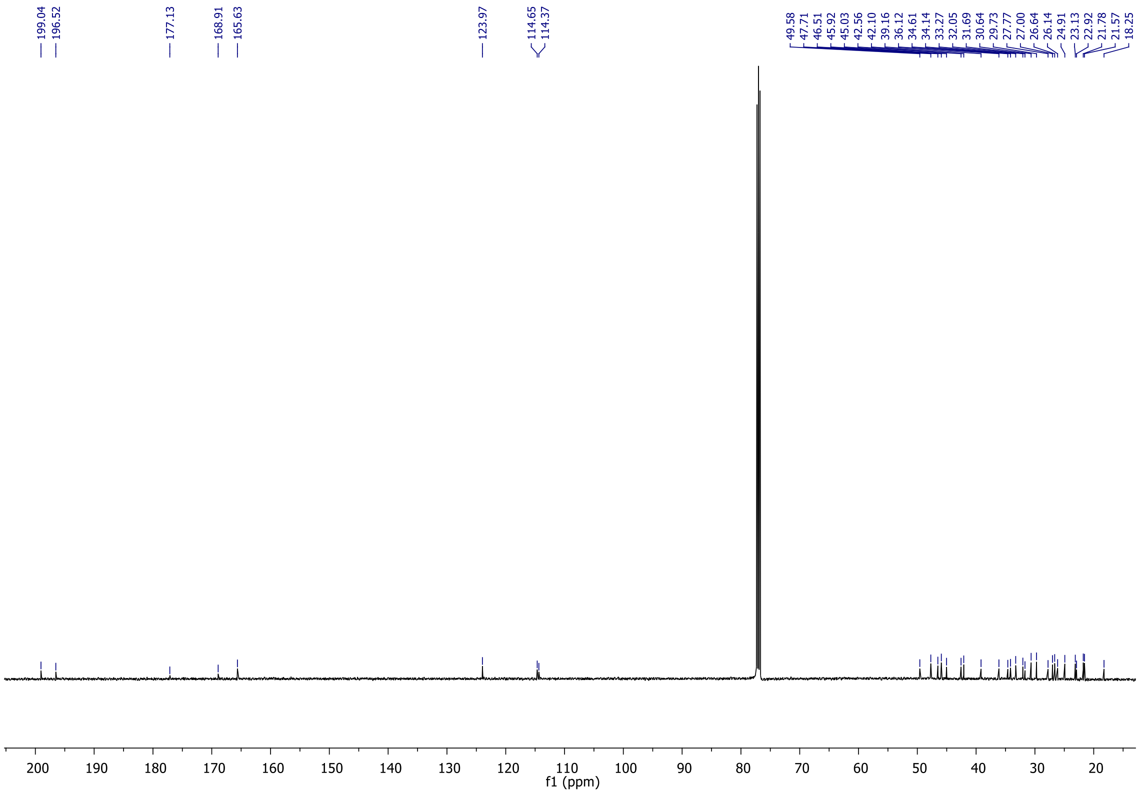
199.04
196.52
177.13
168.91
165.63
123.97
114.65
114.37
49.58
47.71
46.51
45.92
45.03
42.56
42.10
39.16
36.12
34.61
34.14
33.27
32.05
31.69
30.64
29.73
27.77
27.00
26.64
26.14
24.91
23.13
22.92
21.78
21.57
18.25
200
190
180
170
160
150
140
130
120
110
100
90
80
70
60
50
40
30
20
f1 (ppm)

EL230:

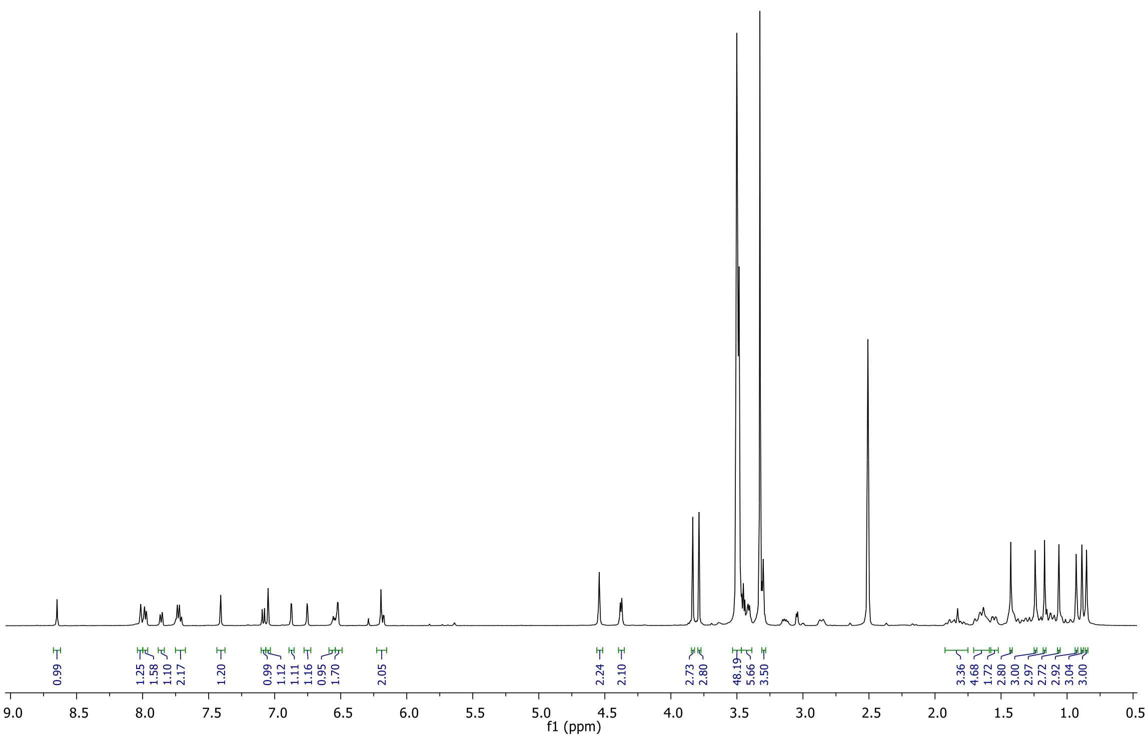
0.99
1.25
1.58
1.10
2.17
1.20
0.99
1.12
1.11
1.16
0.95
1.70
2.05
2.24
2.10
2.73
2.80
48.19
5.66
3.50
3.36
4.68
1.72
2.80
3.00
2.97
2.72
2.92
3.04
3.00
9.0 8.5 8.0 7.5 7.0 6.5 6.0 5.5 5.0 4.5 4.0 3.5 3.0 2.5 2.0 1.5 1.0 0.5
f1 (ppm)

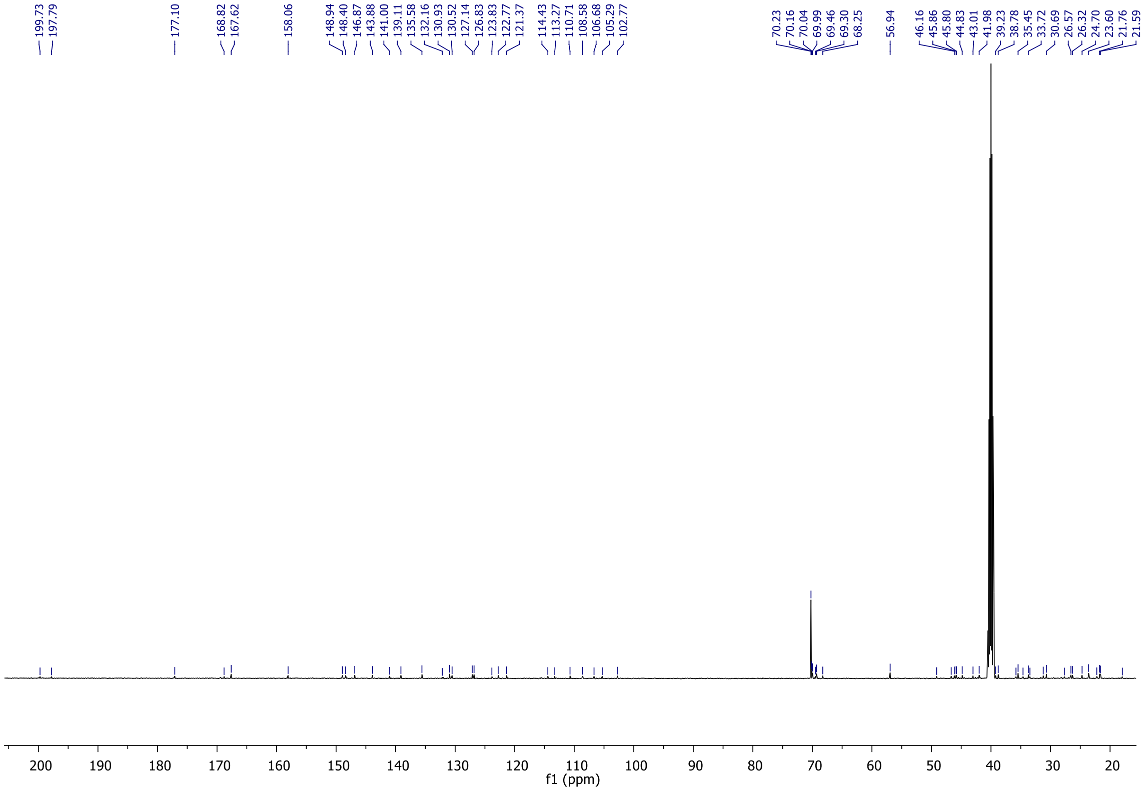
199.73
197.79
177.10
168.82
167.62
158.06
148.94
148.40
146.87
143.88
141.00
139.11
135.58
132.16
130.93
130.52
127.14
126.83
123.83
122.77
121.37
114.43
113.27
110.71
108.58
106.68
105.29
102.77
70.23
70.16
70.04
69.99
69.46
69.30
68.25
56.94
46.16
45.86
45.80
44.83
43.01
41.98
39.23
38.78
35.45
33.72
30.69
26.57
26.32
24.70
23.60
21.76
21.59
200 190 180 170 160 150 140 130 120 110 100 90 80 70 60 50 40 30 20
f1 (ppm)
